# A normative reference for large-scale human brain dynamics across the lifespan

**DOI:** 10.64898/2026.03.06.710057

**Authors:** Yanwu Yang, Nitin Sharma, Sicheng Dai, Francesco Mallus, Guinan Su, Atharva Kand, Mariam Zabihi, Dag Alnæs, Jan K. Buitelaar, Ilona Croy, Richard Dinga, Daniel Durstewitz, Andreas J. Fallgatter, Christian Gaser, Corina Greven, Seyed Mostafa Kia, Milin Kim, Marieke Klein, Max Korbmacher, Jens Krüger, Vinod Kumar Jangir, Esten Leonardsen, Torgeir Moberget, Daan Van Roij, Didac Vidal-Piñeiro, Martin Walter, Yunpeng Wang, Lars T. Westlye, Barbara Franke, Saige Rutherford, Thomas Wolfers

**Author notes:** Corresponding authors **Dr. Yanwu Yang** Humboldt Research Fellow Laboratory for Mental Health Mapping (MHM-Lab) Department of Psychiatry and Psychotherapy University Hospital Tübingen, Tübingen, Germany **Prof. Dr. Thomas Wolfers** Carl-Zeiss Professor for AI in Neural Systems Imaging Laboratory for Mental Health Mapping (MHM-Lab) Friedrich Schiller University Jena, Jena, Germany.

## Abstract

Human brain function emerges from dynamic reconfigurations of large-scale neural networks. While population-level reference charts have transformed the study of static brain structure and connectivity, an equivalent normative framework for intrinsic brain dynamics has been lacking. This gap has limited our ability to characterize individual variability, development, ageing, and mental health conditions at scale. Here, we establish a population-level normative reference for large-scale human brain dynamics using resting-state fMRI data from more than 10,000 individuals spanning the lifespan and 91 scanning sites. We derive a compact set of recurring brain-state configurations that are reproducible across scanners and acquisition paradigms and that generalize to previously unseen cohorts. Anchoring these dynamic states to normative lifespan models enables the quantification of individual deviations relative to population reference distributions. We show that intrinsic brain dynamics undergo systematic reorganization across development and ageing, with pronounced changes before early adulthood and more gradual modulation thereafter. Applying this framework across multiple mental health conditions reveals disorder-specific and highly heterogeneous deviations in brain dynamics that are not captured by static neuroimaging measures. Robust transfer to independent cohorts and longitudinal analyses demonstrate that normative brain dynamics can be reliably assessed out of distribution. These results delineate a population-scale dynamic architecture of the human brain and extend normative brain mapping from static phenotypes to the temporal domain, providing a reference framework for studying brain function across the lifespan in health and disease.

## Introduction

A central aim of neuroscience is to understand how large-scale brain organization varies across development, ageing, and mental health conditions^1–3^. Over the past decade, population-level normative brain charts have reshaped this effort by providing reference distributions for static brain phenotypes, enabling individual-level deviation mapping across the lifespan^4–7^. Such frameworks have proven powerful for studying heterogeneity in both typical development and mental health conditions ^7,8^. However, these advances have focused almost exclusively on static measures of brain structure and connectivity, leaving the dynamic processes that govern moment-to-moment brain function largely uncharted. Accumulating evidence indicates, however, that cognitive functions emerge from the dynamic coordination and competition of large-scale brain network configurations over time^9–11^. Despite their central importance, this brain dynamics remain largely underexplored across brain development and aging, leaving key aspects of the brain function elusive.

Brain function unfolds through metastable transitions between coordinated and decoupled configurations of distributed systems, reflected in time-varying coupling patterns^10,12,13^. Converging evidence from magnetoencephalography (MEG)^14,15^, electroencephalography (EEG)^16,17^, and functional magnetic resonance imaging (fMRI)^18,19^ have shown brain dynamics unfold as structured, recurring patterns of large-scale coordination that continuously reorganize over time^20–22^. Specifically, the concept of data-driven brain states provides a quantitative framework that reduces high-dimensional, continuously fluctuating neural signals into a finite set of recurring and interpretable configurations. By representing brain activity as a sequence of macro-states, brain dynamics can be characterized using temporal measures such as fractional occupancy, dwell time, and state transition probabilities, which capture how network configurations persist and reorganize over time. A range of data-driven approaches, including Hidden Markov Models (HMM)^14,22,23^ and clustering methods^18,24,25^ such as Leading Eigenvector Dynamics Analysis (LEiDA)^26^ and Eigenvector Dynamics Analysis (EiDA)^9^, have identified transient brain states and their temporal transitions within individual datasets. However, how such states relate across cohorts and studies remains poorly understood, owing to the scarcity of well-harmonized large-scale datasets and the lack of brain-state modelling approaches, which together impede the development of reliable and transferable brain-state representations.

Here, we assembled and harmonized a large-scale resting-state fMRI cohort comprising more than 10,000 individuals across 91 scanning sites to establish a transferable normative reference for large-scale human brain network organization. We introduce NeuroLex, a tokenization framework that compresses continuous functional activity into a compact repertoire of 12 recurring brain-state configurations. Using fractional occupancy and dwell time to index state prevalence and temporal stability, we chart normative lifespan trajectories and quantify individual deviations from age-expected patterns. Applied across individuals with attention-deficit/hyperactivity disorder (ADHD), autism spectrum disorder (ASD), major depressive disorder (MDD), anxiety disorders (ANX) and schizophrenia (SCZ), this framework reveals a shared core repertoire of dynamic brain configurations alongside disorder-specific shifts in state expression, delineating both transdiagnostic and condition-specific signatures of altered brain organization. By extending normative modelling from static brain measures to dynamic network organization, this framework reframes brain-state variability as quantifiable deviation from population norms. Generalization to independent, previously unseen cohorts supports transferability and highlights normative deviation in network dynamics as a scalable, clinically relevant marker of neurobiological vulnerability.

**Fig. 1.**
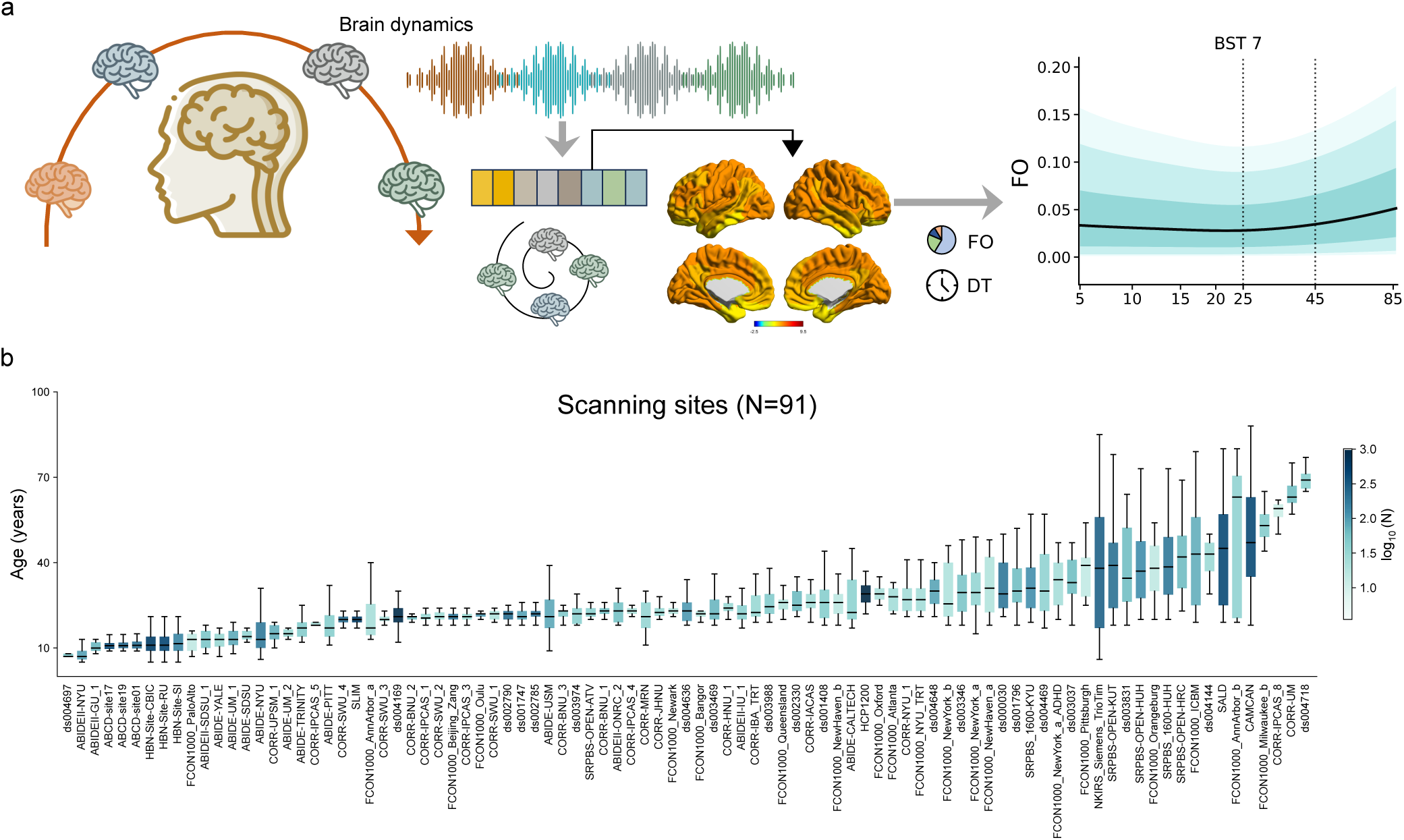
Charting human brain network dynamics at population scale. **a**, Brain dynamics are characterized by recurring large-scale network configurations (referred to as brain state tokens), which are used to quantify individual-level features of brain dynamics, including fractional occupancy (FO) and dwell time (DT). Normative modelling is then applied to chart lifespan trajectories of brain state expressions. An example normative trajectory for a representative brain state (BST7) is shown, illustrating age-related variation in fractional occupancy. Shaded regions denote the variation in healthy reference cohort expressed in percentiles. **b,** Large-scale resting state fMRI data was aggregated across 91 scanner sites, comprising 10,182 individuals spanning an age range from 5 to 88 years. Box plots depict the age distribution of each study, with colour indicating relative sample size. This multi-site reference cohort provides a population-level foundation for learning normative brain-state trajectories that generalize to unseen individuals and out-of-distribution datasets, supporting translational applications.

## Results

### NeuroLex enables reliable quantization of brain network dynamics at scale

To enable population-level quantization of brain dynamics, we used time-resolved phase-locking matrices to model large-scale phase synchronization as the basis of our dynamic representation. We summarized these phase-locking patterns at the network level by projecting brain activity onto the Yeo-7 atlas^27^, yielding a compact description of synchronization within and between seven canonical networks. To ensure cross-study comparability and scalability, we aggregated large-scale resting-state fMRI datasets spanning 91 scanners and harmonized temporal resolution to 2 s. Table 1 summarizes the demographic characteristics of the samples. We then developed NeuroLex, a self-supervised framework that delineates recurrent phase-locking configurations and discretizes them into brain-state tokens (BSTs) (Fig. 2a). Built on the training set, NeuroLex identified 12 recurrent BSTs (Fig. 2b). The number of states was selected by jointly optimizing state separability and split-half reproducibility (Supplementary Fig. S1). These states organized into three higher-order modes reflecting systematic fluctuations between globally synchronized configurations, transient network-specific decoupling, and intermediate hybrid states (Supplementary Fig. S2).

**Table 1.**
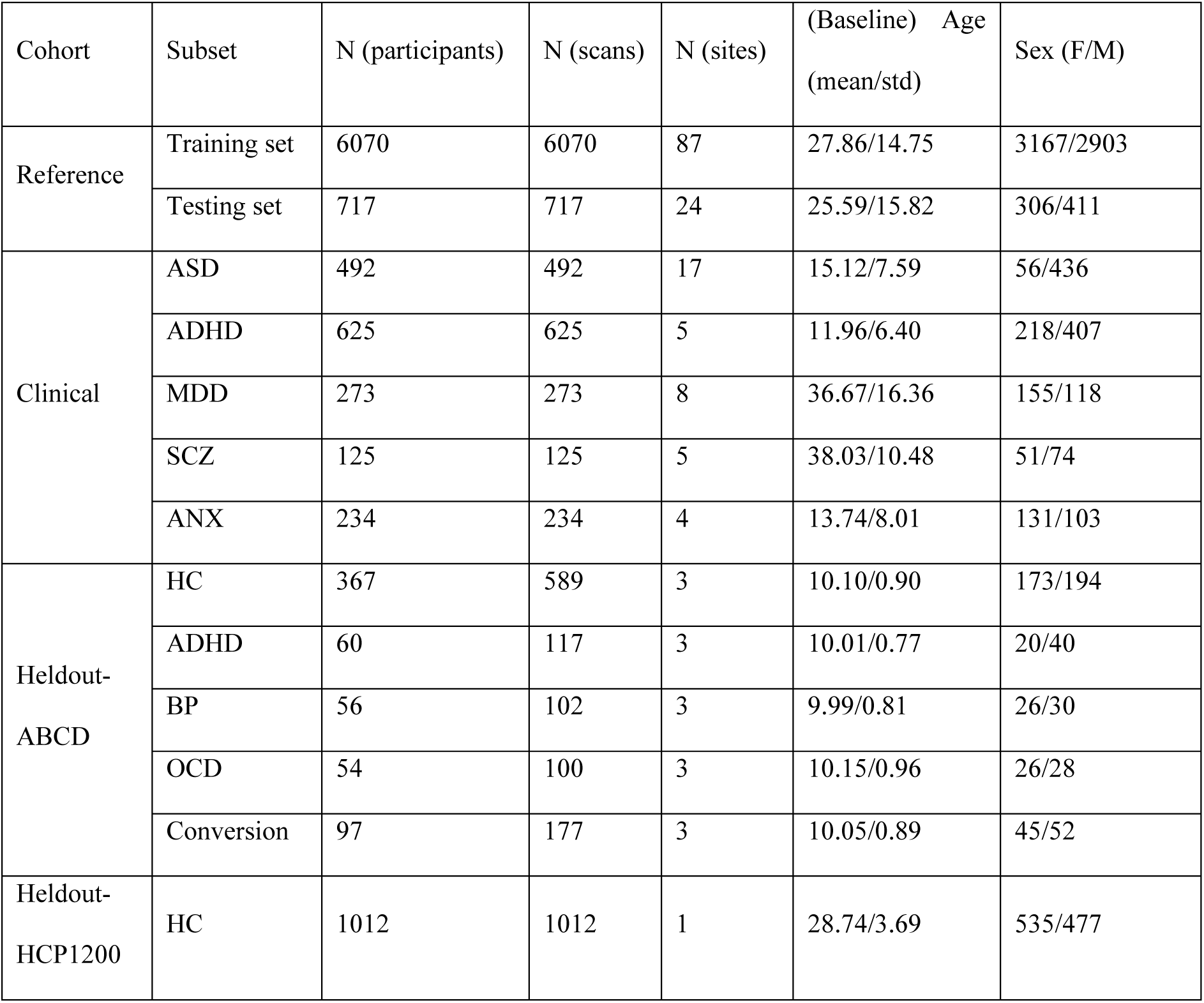
Sample description and demographics.

| Cohort | Subset | N (participants) | N (scans) | N (sites) | (Baseline) Age<br>(mean/std) | Sex (F/M) |
| --- | --- | --- | --- | --- | --- | --- |
| Reference | Training set | 6070 | 6070 | 87 | 27.86/14.75 | 3167/2903 |
|  | Testing set | 717 | 717 | 24 | 25.59/15.82 | 306/411 |
| Clinical | ASD | 492 | 492 | 17 | 15.12/7.59 | 56/436 |
|  | ADHD | 625 | 625 | 5 | 11.96/6.40 | 218/407 |
|  | MDD | 273 | 273 | 8 | 36.67/16.36 | 155/118 |
|  | SCZ | 125 | 125 | 5 | 38.03/10.48 | 51/74 |
|  | ANX | 234 | 234 | 4 | 13.74/8.01 | 131/103 |
| Heldout-ABCD | HC | 367 | 589 | 3 | 10.10/0.90 | 173/194 |
|  | ADHD | 60 | 117 | 3 | 10.01/0.77 | 20/40 |
|  | BP | 56 | 102 | 3 | 9.99/0.81 | 26/30 |
|  | OCD | 54 | 100 | 3 | 10.15/0.96 | 26/28 |
|  | Conversion | 97 | 177 | 3 | 10.05/0.89 | 45/52 |
| Heldout-HCP1200 | HC | 1012 | 1012 | 1 | 28.74/3.69 | 535/477 |

We next evaluated robustness across acquisition heterogeneity and compared NeuroLex with several well-estimated approaches, including GaussianHMM^15,19^, LEiDA^26^, and EiDA^9^. Across scanning sites, NeuroLex yielded more distinctly organized state structure and higher within-subject reproducibility than GaussianHMM, LEiDA and EiDA. In detail, NeuroLex achieved consistently higher silhouette scores across sites (0.474 vs. 0.144/0.143/0.238 for GaussianHMM, LEiDA and EiDA) (Fig. 2c). In within-subject split-half analyses, token occurrence distributions showed strong correspondence between independent halves (ICC = 0.820), exceeding reproducibility of alternative approaches (ICC = 0.716/0.805/0.783; Fig. 2d). Additional site-wise comparisons are provided in Supplementary Fig. S3-S5. These indicate a stable and geometry-consistent brain-state discretization across individuals and sites. The learned representation was highly stable across independent runs (mean R² = 0.89; Fig. 2e). Brain-state tokens retained stable semantic correspondence and similar organization (Supplementary Fig. S6), indicating a robust embedding geometry. Fractional occupancy and dwell time were consistent between training and test cohorts (Supplementary Fig. S7), supporting generalizability.

Having established that BSTs are more robust across sites and cohorts, we next tested whether the quantized states retain interpretable behavioural relevance at the population level. We assessed the generalizability of NeuroLex in an independent HCP1200 cohort (Supplementary Fig. S8) and conducted partial correlation analyses between cognitive measures and the fractional occupancy and dwell time of each BST, controlling for age and sex. Cognitive performance was indexed by the g-factor^28^, including processing speed, crystallized ability, visuospatial ability and memory. Associations between BST expression and cognitive measures were small in magnitude and reached nominal significance (P < 0.05) (Fig. 2f). Only the limbic-decoupling state (BST7) remained significant after false discovery rate correction (fractional occupancy: r= -0.102, corrected P = 0.014; dwell time: r = -0.097, corrected P = 0.026), showing consistency between limbic network function and memory performance. The globally high-coupling state showed a negative association with crystallized ability, whereas the corresponding weakly coupled state showed a positive association. Overall, associations between BST expression and behaviour were modest and distributed across states. Although detectable, effect sizes were small and not concentrated in a subset of strongly cognition-linked configurations, suggesting that the quantized representation captures interpretable behavioural signals without being strongly driven by stable cognitive differences. Finally, to further assess whether the rest-derived BST representation remains informative under a strong distribution shift, we tested whether it could discriminate intrinsically organized rest from externally driven task conditions in unseen participants, without any task-specific training. Rest-task classification using logistic regression on the HCP1200 cohort across seven tasks (rest vs. task) demonstrated that NeuroLex achieved the highest average classification accuracy (73.11%) and the highest area under the curve (80.31%) compared to GaussianHMM (68.45%/74.89%), EiDA (69.57%/76.11%), and LEiDA (71.48%/78.31%) (Fig. 2g and Supplementary Table S3). The superior ability to separate rest from multiple task conditions without task-specific training indicates that the rest-derived BST representation captures more structured organization of large-scale brain dynamics that remain informative under distributional shift. Overall, we find that NeuroLex enables reliable, transferable, and interpretable quantization of brain network dynamics at scale, maintaining stability across sites, cohorts, and cognitive contexts while preserving structured behavioural relevance.

**Fig. 2.**
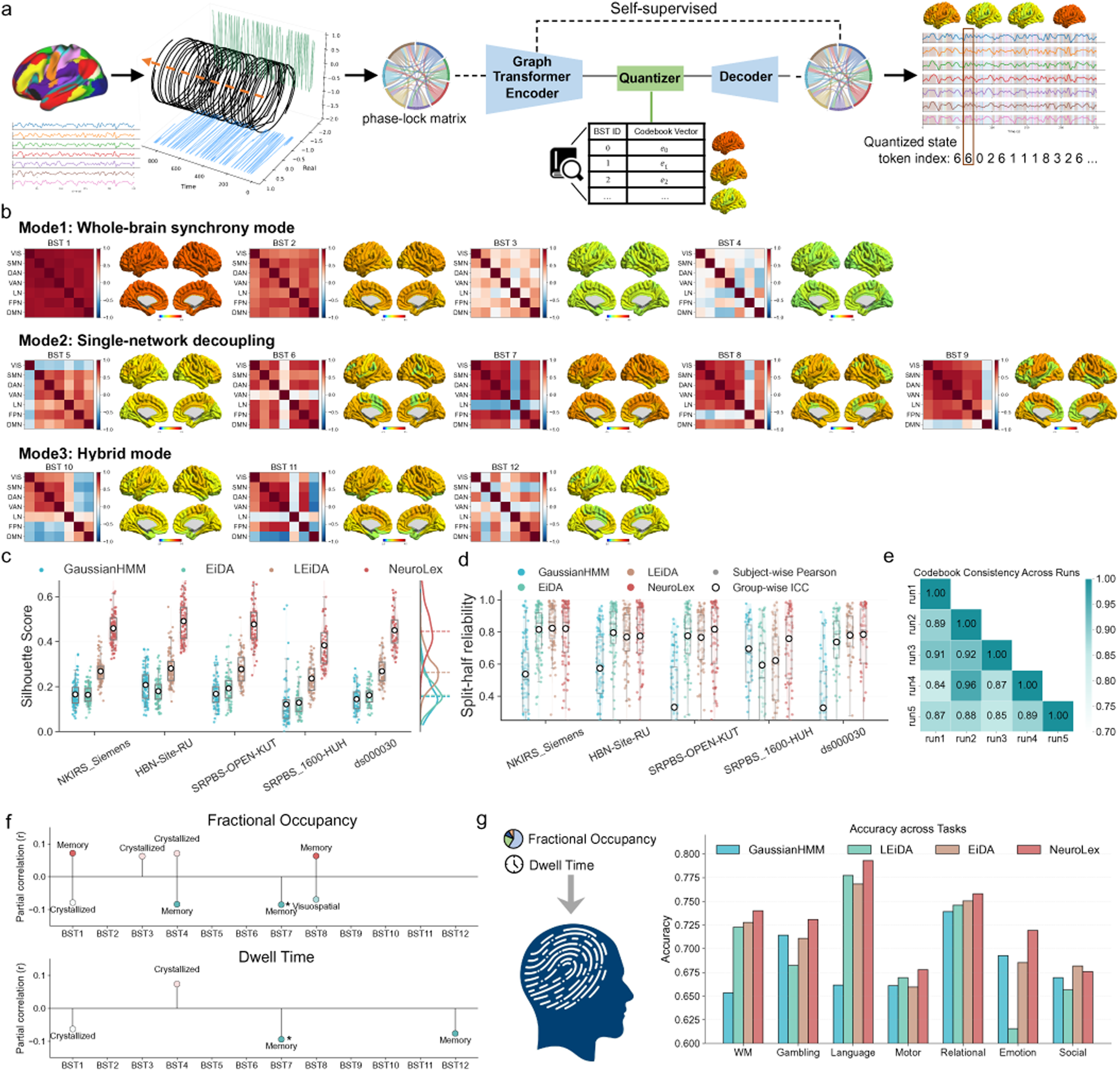
NeuroLex enables reliable identification and characterization of recurring brain network dynamics. **a**, Overview of the NeuroLex method. fMRI signals are transformed into phase-lock matrices, which are then encoded using a graph Transformer and discretized via a learned quantization module in a self-supervised manner. This yields a compact sequence of brain state indices that capture recurring dynamic network configurations over time. **b,** Characteristic brain state patterns identified by NeuroLex, grouped into three dominant modes. Mode1: whole-brain synchrony mode (BST1–BST4); Mode2: a single-network decoupling mode (BST5–BST9); Mode3: a hybrid mode (BST10–BST12). For each state, the phase-lock matrix and corresponding spatial patterns are shown, together with global phase-locking strength. **c,** Clustering quality across five independent sites, quantified by silhouette scores. NeuroLex exhibits consistently stronger cluster separation across sites. **d,** Split-half reliability of brain state representations, evaluated at the individual level via split-half similarity (Pearson correlation) and at the group level using intraclass correlation (ICC). **e,** Codebook consistency across independent runs, indicating stable state representations learned by NeuroLex. **f,** Associations between the general factor (g-factor) and BST temporal measures (fractional occupancy and dwell time) were assessed using age- and sex-adjusted regression models. Results shown reached nominal significance (uncorrected P < 0.05). The limbic-decoupling state (BST7) survived false discovery rate correction. **g,** Rest–task discrimination on the held-out HCP1200 cohort, spanning seven tasks (emotion, gambling, motor, language, social cognition, relational reasoning, and working memory), provided an operational test of cross-context stability under task engagements.

### Brain state tokens reveal functional trajectories across the lifespan

We examined lifespan trajectories of fractional occupancy and dwell time for each brain state token using generalized additive models for location, scale and shape (GAMLSS)^29,30^. This framework enables flexible modelling of non-linear developmental and ageing effects while accounting for site- and study-specific variability through random-effect terms. GAMLSS models were fitted to fractional occupancy and dwell time features derived from the brain state tokens to show lifespan trajectory reference (Supplementary Table S4–S5).

The lifespan curves in Fig. 3 revealed significant nonlinear age effects that were modulated by sex in a subset of brain states, with consistent age-by-sex interactions observed for both fractional occupancy and dwell time after multiple-comparison correction (corrected P < 0.05). BST5, indicating visual-network decoupling, showed a decreasing fractional occupancy across the lifespan. In contrast, BST7, showing limbic decoupling, displayed a distinct non-linear trajectory, where fractional occupancy showed a slight decrease before early adulthood, followed by a marked increase after 23 years of age. This suggests an age-dependent increase in the probability of limbic system disengagement from large-scale networks^31,32^. The hybrid state BST10 showed a slight increase before the age of 25, after which it remained relatively stable. Lifespan trajectories of fractional occupancy across all brain state tokens are described in Supplementary Figs. S9–S11.

In addition, age-related effects in dwell time were generally weaker than those observed in fractional occupancy, indicating that ageing primarily modulates the probability of state occurrence rather than the persistence of individual states. Nevertheless, significant dwell time effects were observed for BST5, BST7, and BST10 (corrected P < 0.05). Notably, BST7 showed a non-linear trajectory, with a mild reduction in dwell time before early adulthood followed by an increase thereafter, mirroring its fractional occupancy profile and indicating longer persistence of limbic-decoupling configurations in later life. Lifespan trajectories of dwell time across all brain state tokens are described in Supplementary Figs. S12–S14. Besides the above state with significant age-related effects, several brain state tokens such as the default-mode-network decoupling state (BST9) exhibited a pronounced increase during early childhood (before age 10), followed by a plateau across later developmental stages, suggesting early stabilization of DMN-related disengagement dynamics^38^.

In addition, dashed lines in the figure ere used to illustrate representative trajectories of structural change, including white-matter volume (WMV), subcortical grey-matter volume (sGMV) and grey-matter volume (GMV), providing an anatomical benchmark for the lifespan dynamics (Supplementary Fig. S15). Structural and functional measures were derived from the same individuals, enabling direct cross-modal comparison. Structural indices exhibited distinct and modality-specific developmental timing, with WMV peaking in early adulthood (29 years), whereas sGMV reached its maximum earlier, around adolescence (14 years), consistent with previous studies^7^. In contrast, functional dynamics showed their most prominent changes around young adulthood (20–25 years). For example, BST7 displayed a non-monotonic pattern in both fractional occupancy and dwell time characterized by an initial decrease followed by a subsequent increase. These results indicate a temporal dissociation between structural maturation and functional reorganization, showing heterogeneous structural–functional trajectories across the lifespan. Functional connectivity provides converging support for these heterogeneous structural–functional trajectories (Supplementary Fig. S16–S17).

Sex-related differences in brain-state dynamics were selectively observed in specific tokens. In particular, BST1, corresponding to a globally coupled brain state, showed higher fractional occupancy and longer dwell time in males than in females (Supplementary Fig. S9). This is consistent with prior evidence^34,35^ that males exhibit stronger global network integration and spend more time in highly synchronized functional configurations, whereas females tend to show more differentiated and dynamic network organization. Additional sex differences were observed in BST4, BST6, BST11, and BST12 for both fractional occupancy and dwell time, as well as in BST7, BST9, and BST10 for fractional occupancy (Supplementary Table S6).

Despite clear qualitative trends in fractional occupancy and dwell time, age-related effects were modest, reflecting the substantial inter-individual variability inherent in resting-state functional dynamics, which limits variance explained at the group level. Importantly, rather than optimizing mean fit, our models establish a lifespan normative reference distribution, enabling calibrated characterization of individual-level deviations.

**Fig. 3.**
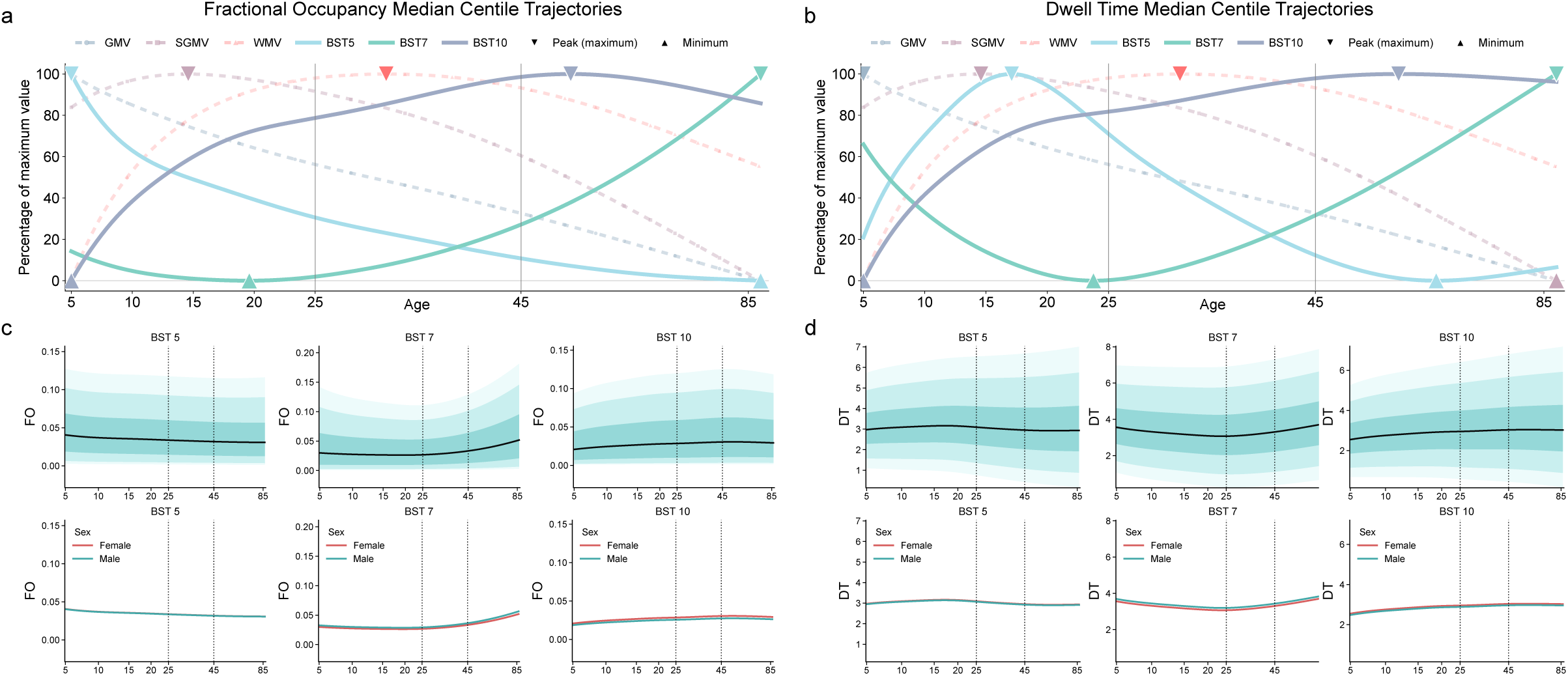
Lifespan organization of recurring human brain network dynamics. Building on the brain state tokens identified by NeuroLex, we characterized how brain state expression changes systematically across the lifespan. **a, b,** Normative lifespan trajectories of fractional occupancy (FO) and dwell time (DT) for the brain-state tokens (5, 7, 10) show significant changes over the lifespan. GMV, sGMV and WMV trajectories are additionally presented for lifespan comparison. **c, d,** Corresponding lifespan trajectories of FO and DT, with shaded regions indicating normative centiles (5th, 25th, 50th, 75th, and 95th percentiles; upper panels) and sex-specific differences shown in the lower panels.

### Brain-state tokens reveal heterogeneous alterations across mental disorders

We quantified individual-level deviations in brain-state dynamics relative to population-based reference norms. Building on this framework, we extended the analysis to five clinical diagnoses, autism spectrum disorder (ASD), attention-deficit/hyperactivity disorder (ADHD), major depressive disorder (MDD), anxiety disorders (ANX), and schizophrenia (SCZ), and evaluated deviations in each diagnosis relative to unseen control individuals without such diagnosis stemming from the test set. A key strength of this framework lies in its ability to enable individual-level characterization relative to a normative reference (Fig. 4a, c). We further performed group-wise comparisons of Z-scores derived from fractional occupancy and dwell time features using Wilcoxon–Mann–Whitney tests (Fig. 4b and d). ASD was primarily characterized by increased fractional occupancy and dwell time in the Limbic-decoupling state (BST7), suggesting enhanced persistence of affective-limbic segregation from distributed cortical systems. Notably, ANX, ADHD, and ASD all exhibited elevated fractional occupancy in this state, potentially reflecting a shared transdiagnostic tendency toward heightened limbic dominance and altered salience-emotion processing dynamics. ANX and ADHD further displayed positive deviations in fractional occupancy for BST2, accompanied by widespread negative deviations in dwell time across multiple states, suggesting reduced state persistence and more frequent transitions, consistent with increased dynamical instability. SCZ showed prolonged dwell time in globally weakly coupled states (BST3), indicating diminished adaptive network segregation combined with a collapse toward low-integration regimes. In contrast, MDD was primarily characterized by alterations in globally coupled states (BST1 and BST3), with reduced fractional occupancy and dwell time, reflecting decreased occupancy and persistence of highly synchronized configurations and a shift toward moderately or weakly integrated regimes. Full statistical results of the group-wise comparisons of normative Z-scores are provided in Supplementary Tables S7–S11.

We next examined the extent to which brain-state dynamics complement conventional structural and static functional markers in characterizing psychiatric disorders. Specifically, we compared dynamic signatures with four global structural MRI measures (GMV, WMV, sGMV, and Ventricular cerebrospinal fluid volume) and inter-network functional connectivity profiles (Fig. 4e). ANX and SCZ showed pronounced sensitivity to structural alterations, with consistent reductions in GMV, WMV, and sGMV, and additional ventricular enlargement in ANX. In contrast, ASD was more prominently reflected in static functional connectivity. MDD and ADHD demonstrated comparatively weak associations with both structural volumes and static connectivity, with ADHD showing only selective hyperconnectivity between the FPN and DMN. In comparison, dynamic features revealed additional and disorder-relevant alterations beyond these conventional measures. ADHD was marked by a significant reduction in dwell time, indicating diminished state persistence that was not captured by structural or static connectivity metrics.

To quantify the relative discriminative contribution of different feature modalities, we supplemented our study with comparative diagnostic classification experiments across feature types. We implemented a support vector machine classifier within a nested stratified five-fold cross-validation framework, with performance evaluated exclusively on out-of-fold predictions to obtain unbiased generalization estimates. For each disorder, healthy controls and patients were randomly sampled at a 1:1 ratio with age and sex match to ensure balanced comparisons, and diagnostic performance was quantified using the area under the receiver operating characteristic curve (AUC) (Fig. 4f). The relative sensitivity of distinct feature categories was highly consistent with the pattern observed in Fig. 4e. Brain dynamics–derived features exhibited superior discriminative performance compared with structural and static functional measures. In particular, dwell time achieved the highest classification accuracy in ADHD (AUC = 0.640) and ANX (AUC = 0.649), whereas fractional occupancy showed comparatively stronger performance in MDD (AUC = 0.591). In all three conditions, dynamic features outperformed both structural volume measures (ADHD: AUC = 0.541; ANX: AUC = 0.597; MDD: AUC = 0.559) and static functional connectivity (ADHD: AUC = 0.600; ANX: AUC = 0.619; MDD: AUC = 0.561). By contrast, SCZ displayed a distinct pattern, with structural volume measures yielding the highest discriminative performance (AUC = 0.716), followed by functional connectivity (AUC = 0.648). Notably, these effects reflect relative discriminative signal across feature modalities under matched case–control sampling, rather than model optimization for maximal accuracy for diagnosis.

We further evaluated the necessity and complementarity of these modalities by AUC using a feature integration strategy (Fig. 4g). Starting from brain dynamics features as the baseline representation, structural and functional connectivity measures were incrementally incorporated to assess additional performance gains. Consistent with the disorder-specific trends described above, structural and connectivity features conferred meaningful improvements primarily in ASD and SCZ. For the other disorders, however, the incremental gains were minimal or absent. These results further support the essential role of brain dynamics representations in clinical analysis, demonstrating that they provide unique diagnostic information beyond that captured by conventional structural volume and static connectivity measures.

**Fig. 4.**
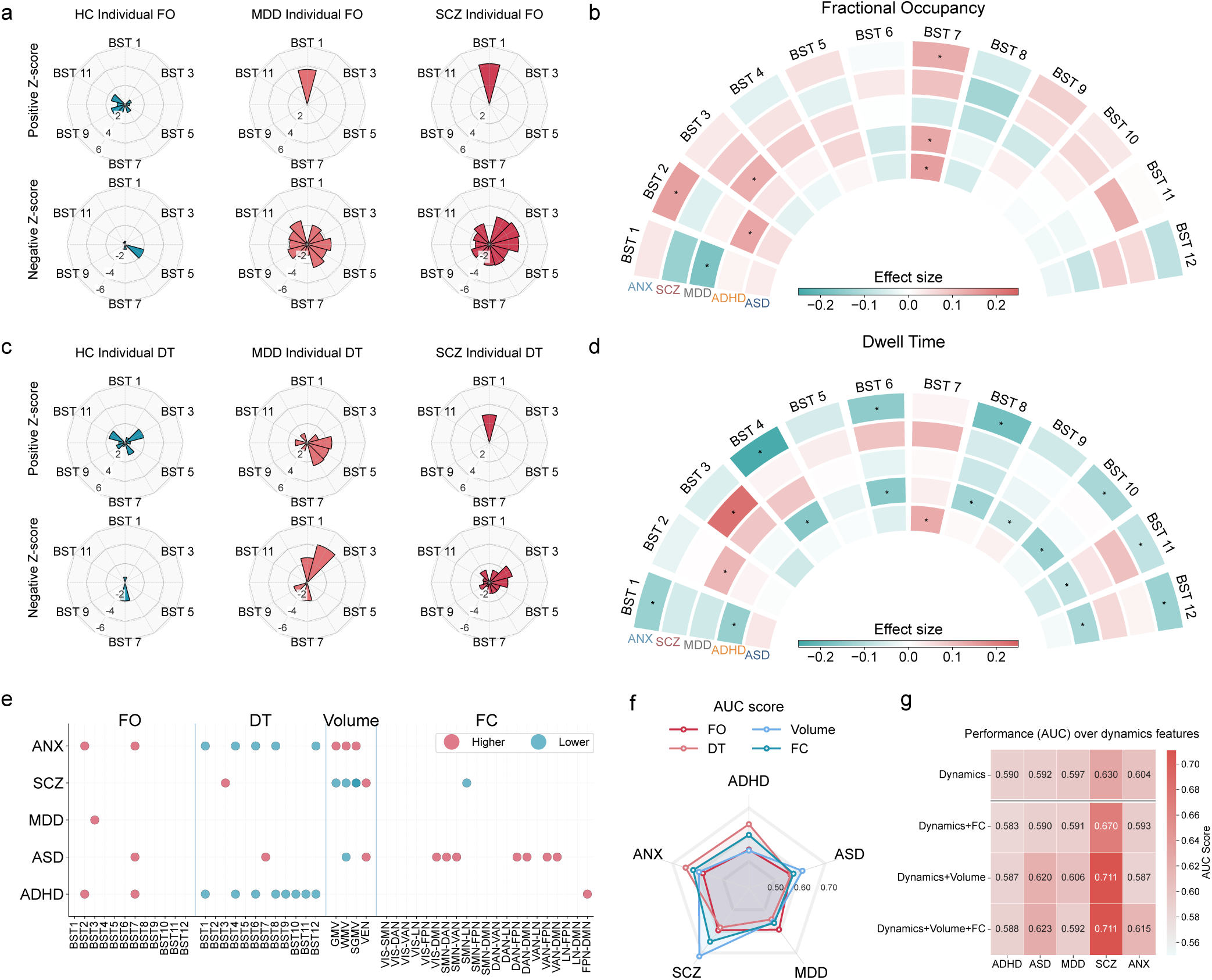
Disorder-specific patterns of deviation in functional brain state token dynamics. Building on the norms, we quantified deviations in brain state dynamics at both the individual and group levels. **a,** Individual-level profiles showing individual-level z-scores of fractional occupancies (FO) relative to the normative reference across brain states. Representative examples from healthy controls, major depressive disorder (MDD), and schizophrenia (SCZ) are illustrated using radar plots. **b,** Group-wise effect size comparison of fractional occupancies relative to the test set across diagnostic groups. Asterisks indicate corrected P < 0.05. **c,** Individual-level deviation profiles showing individual-level z-scores of dwell time (DT). **d,** Group-wise effect size comparison of dwell time relative to the test set. Asterisks indicate corrected P < 0.05. **e,** Cross-disorder comparison of significantly altered features across imaging modalities with significant differences (corrected P < 0.05) for structural volume (GMV, sGMV, WMV), functional connectivity (FC), and brain dynamics metrics (FO and DT). **f,** Classification performance based on stratified nested cross-validation. Area under the curve (AUC) values are shown for each disorder against 1:1 randomly sampled healthy controls, comparing various features. **g,** Additive comparison of classification performance across feature sets. Starting from brain dynamics features (fractional occupancies and dwell time), structural volume and FC measures are progressively added to evaluate changes in AUC across disorders.

### Robust brain-state tokens support clinical translation of estimated norms

A major challenge in neuroimaging analysis is robust generalization to out-of-sample data, requiring stability of both the representation model and normative estimation. We evaluated model transfer on two independent held-out cohorts with unseen scanners: HCP1200, comprising 1,012 healthy control participants, and ABCD, including 367 healthy controls together with 60 ADHD, 56 bipolar disorder, and 54 obsessive–compulsive disorder participants (Fig. 5a), focusing on scanning sites with sufficient sample sizes to ensure stable site-level calibration and enable downstream longitudinal analyses. We showed that recalibrating only site-specific location offsets was sufficient to enable stable transfer of the normative model to these independent cohorts, and in most settings yielded performance comparable to or exceeding that of full refitting of distributional parameters (Fig. 5b). Controlled subsampling in HCP1200, holding out 400 subjects as an independent test set, demonstrated that reliable recalibration could be achieved with as few as 40 to 75 subjects per site. Across 50 repeated runs, ICC reached 0.95 at a calibration sample size of 45 and 0.98 at 100, with corresponding MSE values of 0.982 and 0.972, approaching full-cohort performance (ICC = 0.999; MSE = 0.946; Supplementary Fig. S18). This supports the robustness of our framework in generalization to independent cohorts. Consistent with these findings, transfer closely matched or exceeded full refitting in HCP1200. In ABCD, despite smaller per-site samples and greater heterogeneity, transfer outperformed refitting (Fig. 5b), and most brain-state tokens preserved stable z-score calibration across sites (Supplementary Figs. S19–S20).

This generalization capacity permits individual-level deviation profiling in unseen data (Fig. 5c). Moreover, in group-wise comparison within ABCD, individuals meeting ADHD criteria exhibited significant longitudinal reductions in DMN-decoupling (BST9; Cliff’s delta = −0.21), which is consistent with previous results in Fig. 4e, whereas HC showed stable Z-scores across time (Fig. 5d). Consistently, composite dwell-time deviation, defined as the mean absolute Z-score across 12 brain-state tokens, increased at follow-up in ADHD, indicating broader alterations in brain-state dynamics (Fig. 5e). Slope-based analyses further revealed coordinated reductions across multiple decoupling states, with DMN-decoupling (BST9) remaining significant after correction (Supplementary Fig. S20-S21). In contrast, BP and OCD displayed smaller, non-significant longitudinal changes.

Exploratory time-to-event analyses were performed in the baseline healthy subgroup of the ABCD cohort, comprising 367 individuals who remained clinically stable and 97 individuals who subsequently transitioned to ADHD (n = 30), bipolar disorder (n = 40), or obsessive–compulsive disorder (n = 27) during follow-up. Participants were stratified according to a composite deviation index aggregating fractional occupancy and dwell-time Z-scores across 12 brain-state tokens. Participants were stratified into higher- and lower-deviation groups using an upper-quartile threshold of the composite deviation index. Kaplan–Meier curves (Fig. 5f) showed significant separation between higher- and lower-deviation strata for all three outcomes (log-rank P = 0.0008, 0.0005, and 0.0006), indicating that greater baseline multivariate deviation was associated with elevated transition risk. Individual case profiling (Fig. 5g) further illustrates how deviation-derived indices position individuals along this vulnerability spectrum. These exploratory analyses suggest that normative brain-state deviation encodes structured variability in longitudinal trajectories at the individual level.

**Fig. 5.**
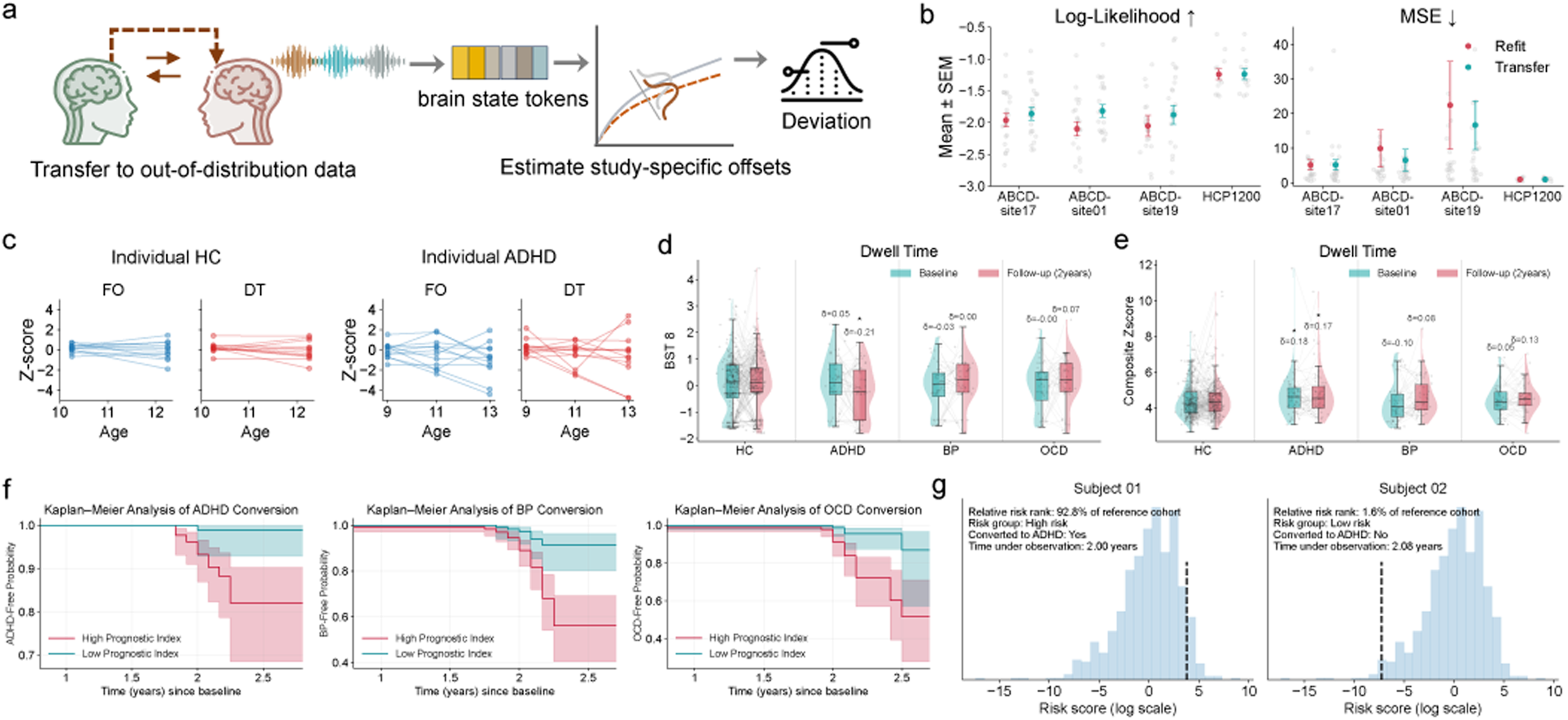
Normative brain state deviations enable out-of-sample transfer and longitudinal analysis. **a**, Transfer of the normative model to independent cohorts by recalibrating site-specific location offsets, enabling generalization. **b,** Comparison of transfer and refitting performance, revealing consistently greater robustness of transfer and pronounced differences in transfer magnitude across sites and scanners. **c,** Transfer-enabled individual deviation profiling in unseen cohorts, permitting individual-level longitudinal analysis. **d–e,** Longitudinal analyses in ABCD revealing reduced DMN-decoupling (BST9) and increased composite dwell-time deviation in ADHD, with other groups remaining stable. **f,** Kaplan–Meier curves stratified by baseline deviation index, indicating elevated transition risk to ADHD, BP and OCD in highly-deviating individuals. **g,** Deviation-based risk stratification highlighting the clinical potential of individualized brain-state profiles.

## Discussion

In this study, we systematically chart large-scale human brain network dynamics and delineate their trajectories across the human lifespan. We show that brain network dynamics can be robustly quantified across cohorts using a set of conserved brain-state tokens, which remain stable across development and aging, and are meaningfully associated with cognitive performance and psychiatric disorders. This unified representation enables large-scale, cross-condition comparisons of brain dynamics and allows us to chart lifespan trajectories of key dynamical features, including fractional occupancy and dwell time. Importantly, deviation scores derived from normative brain dynamics showed robust associations with mental disorders, capturing clinically relevant alterations that may even surpass static markers of brain structure and activity in their sensitivity to disease-related abnormalities. Overall, our study provides a scalable and interpretable framework for characterizing population-level brain dynamics, advancing our understanding of how functional networks organize and adapt across the lifespan in healthy individuals and those with mental disorders.

### Tokenization in brain dynamics

Tokens are discrete representational units that encode contextually meaningful states and form the basic computational primitives in many deep learning models, such as words in language^36^ or patches in vision^37^. In the brain, neural activity likewise unfolds in a continuous manner but organizes into recurrent, metastable patterns of large-scale network configurations. Here, we show that brain dynamics exhibit structured self-organization into discrete brain-state tokens, yielding recurring and interpretable neural configurations analogous to tokenized representations in language and other complex data modalities This framework is grounded in the concept of metastability in dynamical systems, where the brain transiently visits recurring but short-lived states (“attractor ghosts”) that together define its dynamic repertoire. A number of studies on metastability have suggested that brain network dynamics exhibit structured organization^14,17,26^, yet most have been limited to specific datasets or protocols and show weak reproducibility and generalization at the population level. By contrast, NeuroLex achieves better generalizability and discretization performance compared to existing approaches, supporting its ability to extract stable and contextually meaningful state representations of brain dynamics. This scalability enables systematic characterization of brain states across populations and experimental contexts, providing a foundation for studying dynamic brain organization at the population level.

### Lifespan trajectories in brain dynamics

Across the lifespan, brain-state dynamics exhibited moderate age dependence, with most states showing pronounced changes during development and early adulthood. Analyses of fractional occupancy and dwell time consistently indicated that the rate of age-related change attenuated around 20-25 years of age (Supplementary Fig. S9-14), although the precise timing varied across states. Notably, multiple states except BST2, 11, and 12 showed convergent patterns in which age-related gradients approached near-zero values in the mid-twenties in both fractional occupancy and dwell time measures. For example, the limbic-decoupled state (BST7) showed a marked reduction in occurrence before 24 years of age, followed by a gradual increase across adulthood. These indicate that the most pronounced reconfiguration of brain-state dynamics occurs prior to early adulthood, with subsequent changes unfolding at a substantially slower pace. This temporal profile is consistent with prior lifespan neuroimaging studies suggesting that late adolescence and early adulthood mark a transition from rapid developmental reorganization toward more gradual refinement of large-scale brain organization^38–40^.

Beyond early development, brain-state dynamics continued to evolve across middle and later adulthood, with ageing selectively modulating the relative expression of specific states. During midlife, many states exhibited relatively stable trajectories, followed by more pronounced divergences in older age. In particular, the increased occurrence of the limbic-decoupled state (BST7) in older adults suggests progressive changes in interactions between affective and higher-order systems, consistent with age-related alterations reported in frontolimbic brain networks^41,42^. In contrast, the gradual decline of globally synchronized and visual-decoupled states across adulthood points to reduced global coherence and sensory system engagement with increasing age, in line with prior evidence of functional dedifferentiation and declining network efficiency^43,44^.

From a lifespan perspective, these findings suggest that brain-state dynamics are shaped by temporally distinct developmental and ageing-related processes. Early life is characterized by rapid shifts in the prevalence and stability of multiple states, whereas middle adulthood reflects a period of relative stabilization, followed by gradual and state-specific modulation in later life. Such trajectories are consistent with broader theories of brain development and ageing proposing early optimization of large-scale functional dynamics and later adaptive rebalancing under accumulating structural and metabolic constraints^45,46^. The observed age-dependent modulation of brain states may therefore provide a dynamic systems-level substrate linking neurodevelopmental maturation with age-related changes in cognition and behaviour.

### Clinical heterogeneity in the brain dynamics

Combining the integration of brain-state tokenization with normative modelling represents a key advance, as it shifts the analytical focus towards individualized deviation profiles defined relative to population reference norms across brain tokens that represent network dynamics. It makes the approach practical and applicable at the level of the individual.

As illustrated in Fig. 4d, ANX and ADHD were characterized by a global reduction in dwell time, indicating a shift toward diminished temporal persistence of brain states. This pattern points to reduced persistence of engaged brain states and more rapid switching between states, consistent with accounts of ADHD and ANX that emphasize diminished sustained control and increased arousal-driven reactivity, which can destabilize large-scale network configurations and bias the system toward transient, quickly alternating functional states^47,48^. Other diagnostic groups exhibit markedly heterogeneous patterns, with different combinations of fractional occupancy and dwell time abnormalities and varying token-specific signatures, indicating high heterogeneity in brain dynamics across brain disorders.

Beyond disorder-specific differences in overall deviation magnitude, there remains global brain synchronization and segregation across brain disorders, as reflected by shifts across high-to low-coupling brain-state tokens (BST1–BST4). In ASD, deviations were characterized by a redistribution from weakly coupled toward more strongly coupled global states, evidenced by increased fractional occupancy in BST1, BST2 and reduced occupancy in BST3–BST4. MDD exhibited the opposite configuration, with reduced occupancy of highly coupled states (decreased fractional occupancy in BST1) and increased engagement of weakly coupled or segregated states (increased fractional occupancy in BST2–BST4). ANX and ADHD showed a distinct yet partially overlapping profile, marked by increased occupancy of moderately coupled states (BST1–BST2) and reduced occupancy of the weakest coupled state (BST4). Importantly, in addition to these occupancy changes, ANX and ADHD uniquely exhibit a uniform reduction in dwell time across all tokens, indicative of globally shortened state persistence and more rapid switching, whereas in the remaining disorders, dwell time changes largely follow the same direction as fractional occupancy, without evidence for a generalized alteration in temporal stability.

Beyond global coupling patterns, the disorders investigated also exhibited heterogeneous alterations in more specific network-decoupling modes. SCZ, ADHD and ASD both show increased expression of the limbic-decoupled state (BST7), suggesting a shared shift toward reduced integration of limbic circuits within large-scale regulatory networks, a configuration that may reflect impaired affective control and abnormal salience processing^49,50^. Such alterations may point to weakened affective integration and a reduced regulatory influence of limbic signals on higher-order associative systems. In contrast, ADHD displayed pronounced alterations across several single-network decoupling modes, involving the Ventral Attention Network (BST6), the Frontoparietal Network (BST8), and the Default-Mode Network (BST9). These patterns point to a dynamical shift toward increased segregation of ventral attention and frontoparietal control systems from globally integrated states, in line with prior reports of altered interactions between stimulus-driven reorienting and top-down regulatory networks in ADHD^51,52^. This underscores that disorder-related alterations in brain dynamics are not confined to shifts along a single global coupling axis, but instead reflect network-specific deviations in how functional systems transiently disengage and reconfigure over time.

### Strengths and limitations

A central advance of this study is the reframing of large-scale brain dynamics through a tokenization paradigm, establishing a foundation model inspired representation that bridges metastable neural systems with scalable computational abstractions. In doing so, it introduces a complementary lifespan-resolved axis of brain variation, expanding the dimensional framework beyond conventional structural and functional measures to advance system-level understanding of human brain change and disease. Besides, several limitations also warrant consideration. First, although twelve brain-state tokens were reproducible across lifespan, scanners and disorders, they reflect a specific parameterization guided by stability and reproducibility criteria. Alternative model choices may yield additional states or hierarchical decompositions. While spatial patterns were largely consistent across runs (Supplementary Fig. S6), variability in centroid phase-locking patterns suggests partially overlapping or hierarchical organization not fully captured by a finite token set. As a quantization of a continuous dynamical manifold, tokenization assigns configurations to the nearest centroid, potentially introducing boundary effects that influence fine-grained reproducibility and lifespan trajectories. Nonetheless, relative deviation profiles and clinical associations remained stable, including under divergent parcellation schemes (e.g., Yeo-7 vs. Yeo-17; Supplementary Figs. S23–S27), indicating that disease-relevant dynamic deviations are robust to reasonable variations in state specification. Second, to ensure comparability across cohorts with heterogeneous acquisition protocols, we downsampled all timeseries to repetition times to 2 s. Although we found that phase-locking matrices are comparatively robust to changes in temporal resolution (Supplementary Figs. S28–S29), this temporal harmonization primarily facilitates cross-cohort consistency. At the same time, the intrinsic temporal resolution of fMRI, shaped by the hemodynamic response, inevitably constrains sensitivity to rapid state transitions and short-lived neural dynamics. The resulting brain-state tokens therefore reflect large-scale dynamical patterns at the timescale accessible to fMRI rather than neural events at their elementary resolution. Future work will benefit from extending this framework to modalities with higher temporal resolution, such as EEG or MEG, enabling a more fine-grained characterization of fast, transient state dynamics. Third, we mitigated site heterogeneity through the normative modelling, following population-reference strategies used in prior large-scale studies^6,7^. While this approach reduces site-related confounding, residual acquisition and site effects cannot be fully excluded. Finally, the age distribution is uneven, with underrepresentation of infants and individuals in mid-adulthood, representing opportunities for future explorations. In particular, limited infant coverage restricts our ability to characterize early developmental changes in functional brain dynamics and to establish robust normative references for early childhood.

## Conclusion

In this study, we represent continuous neural activity as a finite set of recurrent brain-state tokens and anchoring these representations to population-derived reference norms, our approach moves beyond traditional case-control comparisons toward individualized deviation profiles of brain dynamics. This enables a principled and interpretable quantification of heterogeneity across development and neuropsychiatric conditions while maintaining robustness in large, multi-site datasets. Overall, these findings highlight the added value of a dynamic, normative perspective for identifying both shared and disorder-specific signatures of brain dysfunction. While the present work focuses on large-scale resting-state dynamics to ensure statistical stability and cross-site comparability, the proposed framework is inherently extensible. As larger and more richly sampled lifespan datasets become available, future studies can refine normative references toward higher-resolution state representations and multimodal neuroimaging phenotypes. Overall, brain-state tokenization combined with normative modelling provides a flexible and powerful framework for mapping individual variability in brain dynamics across the lifespan in health and disease, offering translational potential and enabling the longitudinal tracking of individual brain-state trajectories relative to population-level variation.

## Online Methods

### Ethics of the study

Description of informed consent and other ethical procedures is extensively provided in each referenced study, with additional information summarized in Supplementary Table S1. All data processing was conducted within the secure computing environment of the German Network for Bioinformatics Infrastructure (de.NBI), where data storage, and analysis comply with EU General Data Protection Regulation (GDPR) and relevant German data privacy laws was guaranteed. Further this infrastructure is ISO certified, which ensures security and the highest standards.

### Reference cohort

The cohort of individuals was aggregated from 87 scanning sites across multiple studies, including CAMCAN, SALD, SRPBS-OPEN, NKIRS, CORR, ABIDE2, SLIM, SRPBS-1600, ADHD, ds000030, ds001408, ds001747, ds001796, ds002330, ds002785, ds002790, ds003037, ds003346, ds003469, ds003831, ds003974, ds003988, ds004144, ds004169, ds004469, ds004636, ds004648, ds004697, ds004718, ABIDE, and FCON1000. Further details on each study are provided in the corresponding publications (Supplementary Table 1). In total, the reference cohort included 6,787 healthy or non-diagnosed individuals (50.72% females). The participants’ ages ranged from 5 to 88 years (Fig. 1a). Comprehensive information on each scanning site, including sample size, mean age, standard deviation, and sex ratio, is available in Supplementary Table 2. Participants with missing demographic information, missing T1-weighted MRI data, or who had withdrawn from the respective studies were excluded from further analyses. Scans with a total acquisition time exceeding 240 s were included.

### Clinical cohort

As for the clinical datasets, we integrated data from SRPBS-OPEN, NKIRS, SRPBS1600, ds000030, HBN, and ABIDE. To ensure statistical robustness, only clinical groups with more than 100 participants and available diagnostic information were included. Based on these criteria, 1,749 individuals with clinical conditions are included in total and are divided into five diagnostic cohorts: Attention-Deficit/Hyperactivity Disorder (ADHD) (n=625), autism spectrum disorder (ASD) (n=492), Major Depressive Disorder (MDD) (n=273), Schizophrenia (SCZ) (n=125), and generalized anxiety disorder (ANX) (n=234).

### Held-out data

In this study, we aggregated two fully held-out datasets for external evaluation, including task-wise comparisons, behavioural association analyses, and disease-progression analyses. Notably, whereas previous evaluations relied on splits between a reference cohort for training and clinical cohorts with healthy individuals for testing, the held-out datasets used here were acquired at previously unseen scanner sites. This design enables a stringent assessment of out-of-distribution generalization. The first dataset is the HCP1200 cohort, which includes cognitively healthy participants who completed seven task-based fMRI paradigms including working memory, gambling, motor, language, social cognition, relational processing, and emotion processing, as well as resting-state scans. Only participants with available all seven task-fMRI data were included, resulting in a final sample of 1,012 participants. All participants in HCP1200 are healthy individuals, with a repetition time (TR) of 0.72 s.

The second dataset is the ABCD cohort, comprising 634 subjects and 1085 longitudinal fMRI scans, with a TR of 0.8 s. Following a systematic site-level quality assessment, sites exhibiting abnormally elevated global synchronization were excluded, and analyses were further restricted to centres with at least 100 healthy control participants. Three sites (01, 17 and 19) met these criteria and were retained for the analysis. The ABCD dataset was stratified into five groups: healthy controls (HC), ADHD, bipolar disorder (BP), obsessive–compulsive disorder (OCD), and a longitudinal conversion group. ADHD, BP, and OCD groups were defined based on a clinical diagnosis at baseline according to KSADS assessments incorporating concordant parent- and youth-reported evaluations. The conversion group was used exclusively for longitudinal and progression analyses of ADHD, BP and OCD. For progression analyses, individuals were required to be clinically asymptomatic at baseline, operationalized as the absence of any KSADS diagnosis based on both parent- and youth-reported assessments. Longitudinal progression was defined as the emergence of a disorder-specific clinical diagnosis at follow-up. This was applied consistently across ADHD, BP, and OCD outcomes.

### fMRI preprocessing

All functional MRI datasets were preprocessed using fMRIPrep v25.2.3 ensuring a fully standardized and reproducible workflow across sites. The preprocessing steps included slice timing correction, motion correction, susceptibility distortion correction, and spatial normalization to the MNI space using ANTs. Anatomical images were skull-stripped and segmented into gray matter, white matter, and cerebrospinal fluid using FAST (FSL). Functional images were co-registered to their corresponding anatomical images and resampled to 2 mm isotropic MNI space. For the HCP1200 dataset, symmetric preprocessing was adopted to match the left–right acquisition protocol and ensure compatibility with both LR and RL phase-encoding directions.

After the main fMRIPrep pipeline, all datasets underwent ICA-AROMA (Automatic Removal of Motion Artifacts) denoising^53^. We opted for AROMA rather than ICA-FIX, as AROMA does not require site-specific classifier training and thus provides better generalizability and applicability across multi-site cohorts. All preprocessing was performed on the German Network for Bioinformatics Infrastructure (de.NBI) Cloud platform. The AROMA-denoised signals were further nuisance-regressed (including WM, CSF, and six motion parameters) and temporally band-pass filtered (0.01–0.1 Hz). To control for head motion, only scans with mean framewise displacement < 0.3 mm were included in analyses.

### NeuroLex

To achieve better generalization across datasets, we propose a self-supervised learning framework to address the representational inconsistency inherent in traditional clustering approaches, where the absence of an explicit supervisory objective often leads to dataset-specific state spaces. We leverage vector quantization in the latent space to discretize learned representations into a finite vocabulary of brain-state tokens, analogous to tokenization in foundation models, whereby continuous features are mapped into a shared semantic token space. In this regard, we developed NeuroLex that takes as input the phase-locking matrices (PLMs) derived from instantaneous phases of regional fMRI time-series by Hilbert Transform, capturing transient synchronization across brain regions. Each PLM 𝑋_𝑡_ ∈ 𝑅^𝑁×𝑁^is treated as a graph, where nodes correspond to brain regions and edge weights encode phase-locking strength at time 𝑡. The encoder 𝑓_𝜃_(·) consists of a brain network Transformer^54^ with learnable positional embeddings and multi-head self-attention layers:

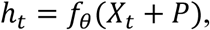

where 𝑃 denotes the positional map and ℎ_𝑡_ ∈ 𝑅^𝐷^ is the latent representation summarizing the instantaneous network topology. The latent vectors are discretized by a vector quantizer 𝑞_𝜙_(·) with a codebook 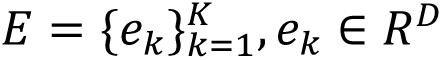. For each ℎ_𝑡_, the nearest embedding is selected as:

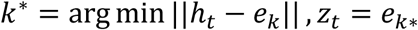

The codebook is initialized by K-means clustering on the first batch and updated through exponential moving averages (EMA) of assignment counts and embedding sums with Laplace smoothing. To prevent codebook collapse, we maintain a usage counter and reinitialize expired embeddings with randomly sampled batch representations once unused for a fixed number of iterations. The quantization is optimized using the straight-through estimator and a commitment loss term:

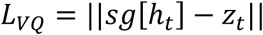

where 𝑠𝑔[·] denotes the stop-gradient operator. The decoder 𝑔_𝜓_(·) maps quantized vectors back to the graph domain via a Transformer stack and reconstructs the corresponding phase-locking matrix:

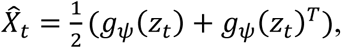

enforcing symmetry in the reconstructed phase-locking patterns. The overall training objective combines reconstruction and vector-quantization losses:

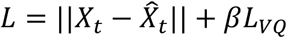

where 𝛽 is a weighting coefficient for the commitment term. This architecture links topology-aware encoding, stable vector quantization, and symmetry-preserving decoding, allowing each fMRI time point to be represented by a discrete brain-state token. These tokens form the basis for quantifying dynamic brain organization in terms of fractional occupancy and dwell time, which are subsequently analysed using lifespan normative modelling.

### Fractional Occupancy

For each participant, the trained NeuroLex model maps every fMRI time point to one of 𝐾 discrete brain-state tokens {𝑠_1_, 𝑠_2_, … , 𝑠_𝑇_}, where 𝑇 is the total number of time frames. The fractional occupancy of state 𝐾 is defined as the proportion of time the brain spends in that state:

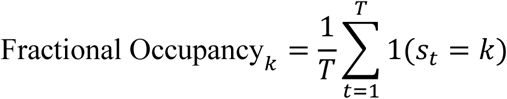

where 1 is the indicator function. This measure reflects the overall prevalence or dominance of each brain state during the recording and serves as a coarse descriptor of the individual’s state distribution. Across participants, fractional occupancy values were used to construct group-level occupancy profiles and lifespan trajectories.

### Dwell time

While fractional occupancy quantifies how often a state occurs, dwell time captures how long it persists once entered. For each subject and state 𝐾, we identify consecutive runs of 𝑠_𝑡_ = 𝑘 and compute their average duration in time frames:

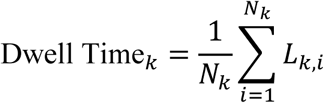

where 𝐿_𝑘,𝑖_ is the length of the 𝑖-th continuous episode of state 𝑘, and 𝑁_𝑘_ is the number of such episodes. When multiplied by the sampling interval of the time series (i.e., the original TR or the post-downsampling interval, when applicable), dwell time yields the mean physical duration (in seconds) for which the brain remains in a given state before transitioning to another.

### Normative modelling

We split the reference cohort (individuals without clinical diagnoses) into training and test sets at the site level. Sites containing diagnostic cases were randomly assigned (50%:50%) to the training and test sets, whereas sites without any diagnostic cases were included exclusively in the training set. Clinical cohorts (ADHD, ASD, MDD, SCZ, and ANX) were held out from model fitting and were only used for evaluation, drawn from the sites assigned to the test split to enable an independent assessment of deviation patterns across disorders. To ensure comparability with the clinical cohorts, individuals in the test set were matched to disease groups by age, sex, and, where possible, site.

Normative models were estimated using the Generalized Additive Models for Location, Scale, and Shape (GAMLSS) framework, implemented in R (version 4.1.2) and accessed via a custom Python–R interface using rpy2. This interface was extended to enable accessible out-of-distribution fits of all GAMLSS parameters designed for transfer learning. The models were fitted with smooth functions of age and sex as predictors, while including the scanning site as a random intercept on the location parameter (𝜇) only, in order to account for site-specific mean offsets while avoiding over-parameterization in centers with limited sample sizes. The response variables were the derived brain-state token features fractional occupancy and dwell time. Models were estimated using the SHASH (Sinh–Arcsinh) family to flexibly capture non-Gaussian distributions and outlier centiles. This approach enables joint modelling of the mean (𝜇), variance (𝜎), skewness (𝜈), and kurtosis (𝜏) parameters as functions of covariates, allowing a comprehensive characterization of lifespan trajectories. Each BST-based feature, denoted by 𝑦, was modelled by:

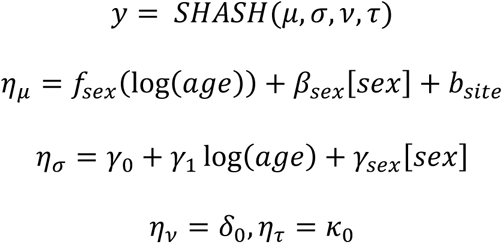

For each test participant, normative deviations (z-scores) were derived by standardizing model predictions relative to the reference cohort. This modelling pipeline was implemented entirely in Python with R-based backend estimation for numerical stability and direct compatibility with the GAMLSS framework. We calculated the z-score (the normalized quantile residual):

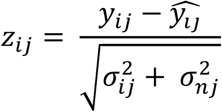

The computation of the z-score includes predicted mean 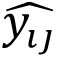, true response 𝑦_𝑖𝑗_, predicted conditional variance 𝜎_𝑖𝑗_ and residual variance 𝜎_𝑛𝑗_ estimated from the reference population. For model evaluation, the GAMLSS-based normative model generated point estimates of the four distributional parameters (𝜇, 𝜎, 𝜈, 𝜏) and corresponding performance metrics, including explained variance, mean squared log-loss, skewness, and kurtosis, computed on the test set containing only non-diagnosed participants. These metrics quantified the model’s goodness of fit and the degree to which predicted distributions captured the empirical variability of the data.

### Group comparisons

We performed classical nonparametric test on the z-scores of clinical cohorts and individuals without diagnosis. To assess the statistical significance, we performed Mann– Whitney U–tests, a non-parametric test that is suitable for comparing two independent samples that are not normally distributed. To control for multiple testing, p-values were adjusted using the false discovery rate (FDR) procedure.

## Data availability

In this study, we used brain imaging from multiple large-scale open-access neuroimaging initiatives, including ABIDE, ABIDE II, ADHD200, CAMCAN, CORR, FCON1000, SALD, SLIM, NKIRS, SRPBS-OPEN, SRPBS1600, and several OpenNeuro datasets (ds000030, ds001408, ds002790, ds003469, ds003831, ds003974, ds003988, ds004349, ds004469, ds004648, ds004697, ds004718), as well as the HCP1200 and ABCD project.

### Code availability

The code used in this study is publicly available on GitHub at https://github.com/podismine/NeuroLex. Actively maintained forks are hosted at https://github.com/MHM-lab, which also provides the GAMLSS-based normative modeling toolkit used in this work. All trained models are shared through these repositories and can be freely downloaded and applied for non-commercial use. Software packages include fMRIPrep v25.2.3 (https://fmriprep.org/en/stable/), ICA-AROMA (https://github.com/maartenmennes/ICA-AROMA), R v4.1.2 (https://www.r-project.org/), MATLAB R2018b (https://www.mathworks.com/products/matlab.html), BrainNetViewer toolbox (https://www.nitrc.org/projects/bnv/), Python v3.10.12 (https://www.python.org/), GAMLSS package v5.4.22 (https://www.gamlss.com/), scikit-learn v1.5.2 (https://scikit-learn.org), Nilearn v0.11.0 (https://nilearn.github.io/stable/index.html), Matplotlib v3.9.2 (https://matplotlib.org/).

## Acknowledgements

We thank all participants and the clinicians and researchers involved in recruitment and assessment. YY and TW gratefully acknowledge support from the Alexander von Humboldt Foundation for YY’s research fellowship. TW acknowledges funding from the German Research Foundation (DFG; Emmy Noether Programme, grant 513851350), the BMBF/DLR project FEDORA (01EQ2403G) and funding from the Carl-Zeiss-Stiftung (P2022-00-087). This work was supported by the BMBF-funded de.NBI Cloud within the German Network for Bioinformatics Infrastructure (de.NBI; grants 031A532B, 031A533A, 031A533B, 031A534A, 031A535A, 031A537A–D, 031A538A). BF acknowledges funding from the Dutch Ministry of Education, Culture and Science of the government of The Netherlands for the NWO Gravitation programme GUTS (grant 024.005.011). The authors used data from the Adolescent Brain Cognitive Development Study (ABCD; https://abcdstudy.org). ABCD data are curated within the NIMH Data Archive (NDA). The ABCD Study is supported by the National Institutes of Health and additional federal partners under the following award numbers: U01DA041048, U01DA050989, U01DA051016, U01DA041022, U01DA051018, U01DA051037, U01DA050987, U01DA041174, U01DA041106, U01DA041117, U01DA041028, U01DA041134, U01DA050988, U01DA051039, U01DA041156, U01DA041028, U01DA041134, U01DA050988, U01DA051039, U01DA041156, U01DA041089, U24DA041123, and U24DA041147. A full list of supporters is available at https://abcdstudy.org/federal-partners.html. A list of participating sites and a complete listing of ABCD Study investigators can be found at https://abcdstudy.org/consortium_members/. ABCD consortium investigators designed and implemented the study and/or provided data but did not participate in the analysis or writing of this manuscript. The views expressed herein are those of the authors and do not necessarily reflect the views of the ABCD consortium, the NIH, or any other affiliated organization.

## Author Contributions

Y.Y. and T.W. conceived the study and designed the overall analytical framework. Y.Y. developed and implemented the NeuroLex tokenization approach, performed data preprocessing, large-scale modelling, and statistical analyses. T.W. guided the methodological development and supervised the project. T.W. and N.S. contributed to the analytical development and supported the statistical analyses. S.D. and T.W. contributed to data curation and dataset development. Y.Y. and T.W. jointly interpreted the results and wrote the first manuscript. All authors provided critical feedback on the interpretation of the results, contributed to manuscript revision, and approved the final submission.

## Roles of funding

The funders had no influence on the study design, analyses or interpretations.

