## Supplementary Information for "A normative reference for large-scale human brain dynamics across the lifespan"

### Table of Contents

Common nomenclature ... 2
Supplementary Methods ... 3
1. Statistics & Reproducibility ... 3
2. Brain Atlases and Parcellation Schemes ... 3
3. NeuroLex Implementation ... 4
Supplementary Results ... 5
4. Determination of the Number of Brain State Tokens ... 5
5. Explanation of Brain State Tokens ... 6
6. Comparative evaluation of brain-state clustering performance ... 8
7. Comparison of split-half reliability performance ... 9
8. Consistency and reliability of brain state clustering ... 10
9. Cross-run stability of brain-state token spatial patterns ... 11
10. Generalization Performance of NeuroLex across the train and test set ... 12
11. Generalization capability to internal test and external datasets ... 14
12. Evaluations of GAMLSS model specifications ... 15
13. Normative trajectories of fractional occupancy across the lifespan ... 16
14. Normative trajectories of dwell time across the lifespan ... 18
15. Structural normative trajectories across the lifespan ... 20
16. Functional connectivity normative trajectories across the lifespan ... 22
17. Sensitivity Analysis of cross-cohort transfer sample size ... 24
18. Cross-cohort transferability of normative models ... 25
19. Application to the ABCD cohort and longitudinal slope analysis ... 26
20. Atlas-level comparison with the Yeo-17 network parcellation ... 28
21. Effects of temporal downsampling on brain state estimation ... 31
Supplementary Tables ... 34
22. Supplementary Table 1. Sources of all the studies used in the study ... 34
23. Supplementary Table 2. Overview of the included datasets ... 38
24. Supplementary Table 3. Rest-task discrimination in the held-out HCP1200 cohort ... 42
25. Supplementary Table 4. Comparison of model fitting across different function configs ... 43
26. Supplementary Table 5. Statistical significance of age-related effects modulated by sex of the lifespan ... 44
27. Supplementary Table 6. Statistical significance of sex-related effects of the lifespan ... 45
28. Supplementary Table 7. Group-wise comparison of normative z-scores between ASD and test HC ... 46
29. Supplementary Table 8. Group-wise comparison of normative z-scores between ADHD and test HC ... 47
30. Supplementary Table 9. Group-wise comparison of normative z-scores between MDD and test HC ... 48
31. Supplementary Table 10. Group-wise comparison of normative z-scores between SCZ and test HC ... 49
32. Supplementary Table 11. Group-wise comparison of normative z-scores between ANX and test HC ... 50
References ... 51

### Common nomenclature

ADHD - Attention deficit hyperactivity disorder

AIC - Akaike Information Criterion

ANX - Anxiety or phobic disorders

ARI - Adjusted rand index

ASD - Autism spectrum disorder

BIC - Bayesian information criterion

CN - Control or cognitively normal

CSF - Ventricular cerebrospinal fluid

DAN - Dorsal attention network

DMN - Default mode network

DT - Dwell time

DX - Diagnosis

FDR - False discovery rate

FO - Fractional occupancy

FPN - Frontoparietal Control Network

GAMLSS - Generalised additive models for location scale and shape

GMV - Total cortical grey matter volume

HC – Healthy control

ICC - Intraclass correlation coefficient

IQR - Interquartile range

LN - Limbic network

MDD - Major depressive disorder

MRI - Magnetic resonance imaging

sGMV - Total subcortical grey matter volume

SCZ – Schizophrenia

SMN - Somatomotor network

VAN - Ventral attention network

VIS - Visual network

WMV - Total cortical white matter volume

### Supplementary Methods

#### 1. Statistics & Reproducibility

All datasets were preprocessed using fMRIPrep (version 25.2.3) within the de.NBI cloud platform to ensure identical preprocessing across cohorts. The preprocessing included spatial normalization, confound regression, and ICA-AROMA-based denoising to minimize motion-related artifacts across multi-site data. The preprocessing code is available here: https://github.com/nipreps/fmriprep and https://github.com/maartenmennes/ICA-AROMA. When translating our models into external datasets or unseen clinical cohorts, particularly in a federated setting, this preprocessing pipeline should be followed.

Functional timeseries were derived from Yeo-7 parcellations[1] and converted into phase-locking matrices. Brain state representations were learned from the reference cohort (healthy or no diagnosis participants) using our NeuroLex model, which can be applied to your data after preprocessing. To ensure reproducibility, we repeated the tokenization procedure with five independent random seeds, computing silhouette and adjusted Rand index (ARI) scores across runs to evaluate stability. Sites with fewer than ten participants or with abnormal phase-locking dynamics were excluded, which should be checked before you deploy our models.

The reference cohort included 6,787 individuals (healthy participants across 31 datasets and 87 scanning sites), split into training (6,070 individuals) and testing sets (717 individuals). No statistical method was used to predetermine sample size; instead, all available participants meeting the quality criteria were included (mean framewise displacement $< 0.3$ mm). All models and analyses were implemented in Python 3.11, using PyTorch 2.3.1, and executed on the de.NBI HPC platform with Singularity containers to ensure computational reproducibility.

#### 2. Brain Atlases and Parcellation Schemes

All analyses in this study were performed using the Yeo 7-network functional atlas[1]. This atlas provides a coarse-grained yet neurobiologically interpretable partition of the cerebral cortex into seven large-scale intrinsic connectomes that capture the dominant modes of spontaneous functional organization. These include the visual network (VIS), the somatomotor network (SMN), the dorsal attention network (DAN), the ventral attention/salience network (VAN), the limbic network (LN), the Frontoparietal control network (FPN), and the default-mode network (DMN). Respectively, they correspond to cortical regions supporting visual perception and sensory encoding, motor execution and somatosensory integration, goal-directed and visuospatial attention, salience detection and network switching, affective and mnemonic integration, executive regulation and working memory, and internally oriented self-referential cognition. A comparison between the Yeo-7 and Yeo-17 parcellations is presented in the Supplementary Fig. S23-S27, demonstrating consistency across atlases. We adopted the Yeo-7 atlas rather than higher-resolution parcellations because our focus was on population-level generalizable brain states. Owing to functional variability across individuals and sites, fine-grained atlases increase the risk of introducing hybrid states. In large, heterogeneous datasets, minor spatial misalignments or scanner-specific noise disproportionately affect smaller parcels, leading to fragmented and more unstable states.

#### 3. NeuroLex Implementation

Neurolex was trained using the Adam optimizer with an initial learning rate of $3 \times 10^{-4}$ and a weight decay of $5 \times 10^{-5}$. The batch size was fixed at 256. Models were trained for 1,000 epochs, and the best-performing checkpoint was selected based on the validation loss. The Transformer encoder consisted of four layers, with four heads per multi-head self-attention module. The hyperparameter $\beta$ was set to 0.1 based on a selected grid search over the values 0.01, 0.05, 0.1, 0.2, 0.5, and 1.0.

### Supplementary Results

#### 4. Determination of the Number of Brain State Tokens

To determine the optimal number of discrete brain states, we systematically varied the number of tokens from 8 to 32 and evaluated the resulting state representations across multiple runs. For each configuration, we assessed clustering quality using two criteria: the Silhouette score and split-half reproducibility ($R^2$). Specifically, for each run, 90% samples of train set were randomly sampled to train the model, while the test set was fixed for evaluation. The Silhouette score was computed on the test set to quantify within-state compactness and between-state separation, whereas $R^2$ was computed to assess clustering consistency and reproducibility across runs. As shown in the left of Supplementary Fig. S1, the Silhouette score increases first and then gradually decreased as the number of tokens increased, indicating that cluster compactness and separability declined with increasing representational granularity. In contrast, $R^2$ decreased, indicating reduced cross-split reliability at higher resolutions. This divergence reflects a fundamental trade-off between representational granularity and population-level stability in discrete brain-state modeling.

To balance these two objectives, we normalized the two metrics into $[0,1]$ and combined them into a composite score with equal weighting. As shown in the right of Supplementary Fig. S1, token numbers in the range of 10 to 14 achieved consistently favorable performance across all criteria. Among these, a token number of 12 yielded the highest composite score, representing an optimal balance between state separability and reproducibility. Consequently, we selected 12 brain states as the optimal resolution for all subsequent analyses. This choice is consistent with previous studies reporting approximately 10–14 metastable brain states in resting-state fMRI[2–6].

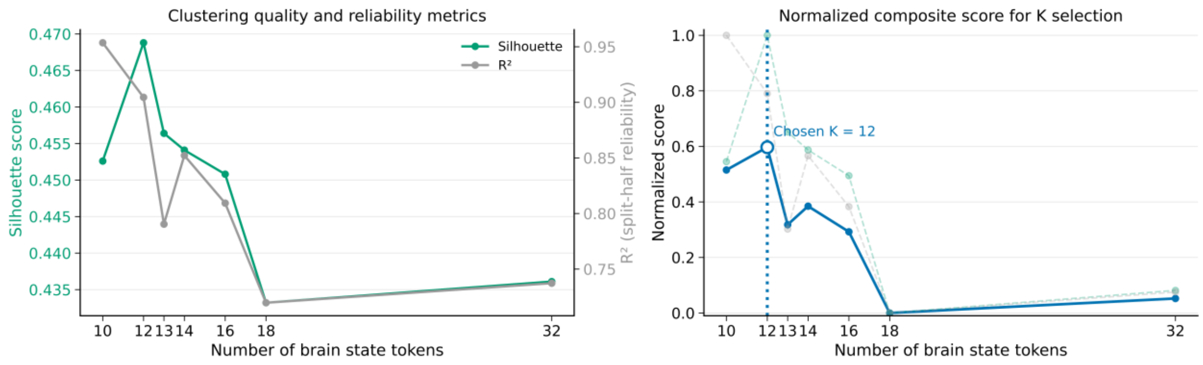

**Supplementary Fig. S1. Selection of the number of brain state tokens. Left,** clustering quality and reliability were evaluated as a function of the number of brain-state tokens, using the Silhouette score and split-half reproducibility ($R^2$). **Right,** normalized metrics and their composite score used for model selection. Both metrics were normalized to the range [0, 1] and combined with equal weighting to balance cluster separability and reproducibility.

#### 5. Explanation of Brain State Tokens

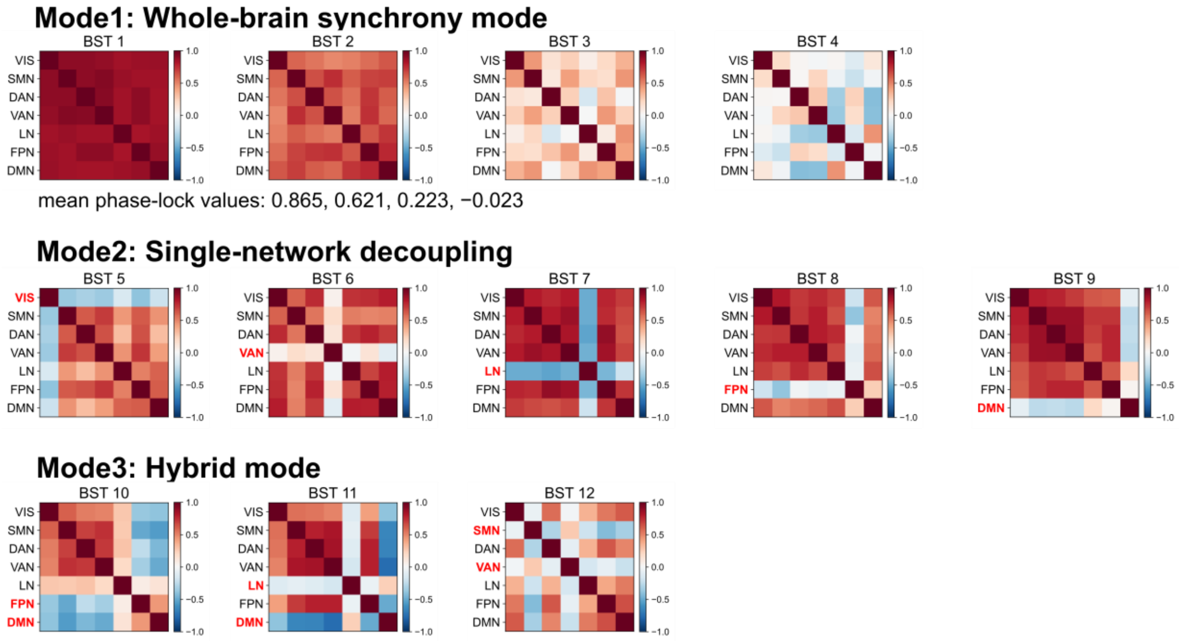

**Supplementary Fig. S2. Interpretation of the 12 brain-state tokens.** We identified 12 recurrent brain-state tokens derived from phase-lock dynamic connectivity patterns and organized them into three dominant modes of large-scale coordination. Mode1 (BST1–BST4) reflects a graded axis of global integration–segregation, Mode2 (BST5–BST9) captures single-network decoupling within otherwise synchronized configurations, and Mode3 (BST10–BST12) represents hybrid coalition states characterized by selective inter-network coupling and competitive segregation.

Here we further elaborate on the interpretability of the 12 brain-state tokens (BSTs), organized into three dominant dynamic modes reflecting distinct principles of large-scale phase coordination among the seven canonical subnetworks.

Mode1: BST1–BST4 form a coherent gradient of global metastable integration. These states exhibit widespread inter-network phase alignment, with synchrony progressively decreasing from BST1 to BST4. The mean inter-subnetwork phase-locking values follow a monotonic decline (0.865, 0.621, 0.223, -0.023), indicating a systematic transition from strongly integrated to weakly or near-segregated configurations. Functionally, these states capture whole-brain coordination and its gradual dissolution. Rather than representing discrete categorical events, Mode1 reflects a continuous modulation axis of global integration, analogous to shifts between highly coherent and more differentiated large-scale system organization.

Mode2 comprises states in which one canonical subnetwork transiently decouples while the remaining networks maintain relatively high synchrony. These configurations preserve global structure but introduce focal functional specialization:

- BST5: Visual network (Vis) decoupling
- BST6: Ventral attention network (VAN) decoupling
- BST7: Limbic network (LN) decoupling
- BST8: Frontoparietal network (FPN) decoupling
- BST9: Default mode network (DMN) decoupling.

These single-network states likely reflect transient functional recalibration within otherwise integrated dynamics. Such patterns may index moments of task-negative disengagement, affective modulation, attentional reorientation, or executive gating, depending on the network involved. Importantly, they demonstrate that metastability is not limited to global synchronization shifts but also includes structured, network-specific departures from coherence.

Mode3 captures hybrid configurations in which specific subnetworks form internally coherent sub-blocks while collectively decoupling from the remaining systems.

- BST10: Global coordination is preserved among most networks, while FPN and DMN jointly decouple from the rest but remain mutually coupled. This configuration may reflect a competitive yet cooperative interaction between executive control and internally oriented processes, potentially indexing internally driven cognitive organization.
- BST11: A similar structure is observed for limbic and DMN networks, suggesting coordinated affective–internal processing dissociated from broader cortical integration.
- BST12: Somatomotor (SMN) and VAN networks form a coupled module segregated from other systems. Given the involvement of motor and salience-related regions, this state may partly reflect embodied or sensorimotor–attentional alignment; alternatively, it may capture movement-related or arousal-related fluctuations that persist even under resting-state conditions.

Collectively, Mode3 illustrates that metastable organization includes structured subcoalitions, not merely global integration or single-network disengagement. These hybrid states reveal competitive-cooperative motifs among cognitive-control, affective, and sensorimotor systems, offering a higher-order view of large-scale dynamic architecture.

#### 6. Comparative evaluation of brain-state clustering performance

We compared the clustering separability of brain state representations across the largest ten independent scanning sites using the silhouette score as an index of within-cluster compactness and between-cluster separation (Supplementary Fig. S3). For each site, silhouette scores were computed across multiple runs for NeuroLex and three baseline methods (GaussianHMM, EiDA, and LEiDA), allowing direct comparison of clustering quality under heterogeneous acquisition conditions. Across the largest ten scanning sites in the test set, NeuroLex consistently achieved higher silhouette scores than the baseline methods. While silhouette values varied across sites, reflecting differences in data characteristics and cohort composition, NeuroLex maintained a clear separation from GaussianHMM and EiDA at every site and generally outperformed LEiDA as well. The distribution of silhouette scores for NeuroLex was also more concentrated, indicating reduced variability across runs compared with the baseline approaches. In contrast, GaussianHMM and EiDA showed systematically lower silhouette scores across sites. LEiDA exhibited intermediate performance, with higher silhouette scores than GaussianHMM and EiDA at some sites but remained consistently below NeuroLex.

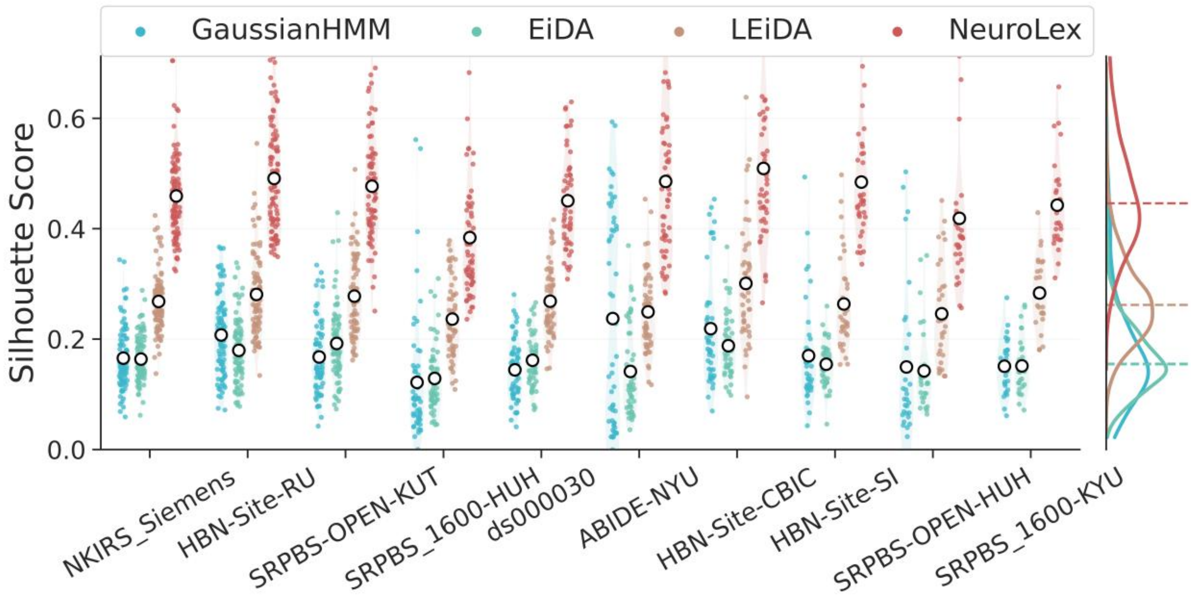

**Supplementary Fig. S3. Cross-site comparison of clustering separability in terms of silhouette scores of brain state representations across ten independent scanning sites for NeuroLex and three baseline methods (GaussianHMM, EiDA, and LEiDA).** Each point represents the silhouette score from a single run, with open circles indicating scanner-wise means. NeuroLex consistently shows higher silhouette scores across scanners, indicating improved cluster separability under heterogeneous acquisitions. Density plots on the right summarize the overall distribution of silhouette scores across all sites for each method.

#### 7. Comparison of split-half reliability performance

We assessed the reliability of brain state representations using split-half reproducibility analyses across the largest ten independent scanner sites (Supplementary Fig. S4). Reliability was quantified at the individual level via split-half similarity (Pearson correlation) and at the group level using intraclass correlation (ICC) respectively. Across sites, NeuroLex showed consistently high split-half reproducibility at both the subject and group levels. Split-half similarity was generally higher and less variable for NeuroLex compared with other methods, indicating more stable individual-level representations across splits. At the group level, NeuroLex also achieved higher ICC values across most sites, reflecting increased consistency of population-level brain state patterns. In contrast, GaussianHMM and EiDA exhibited lower split-half reproducibility with greater variability across sites, while LEiDA showed intermediate reproducibility across both metrics. Although reproducibility values varied across datasets, reflecting differences in acquisition protocols and sample composition, the relative ordering of methods was consistent across sites.

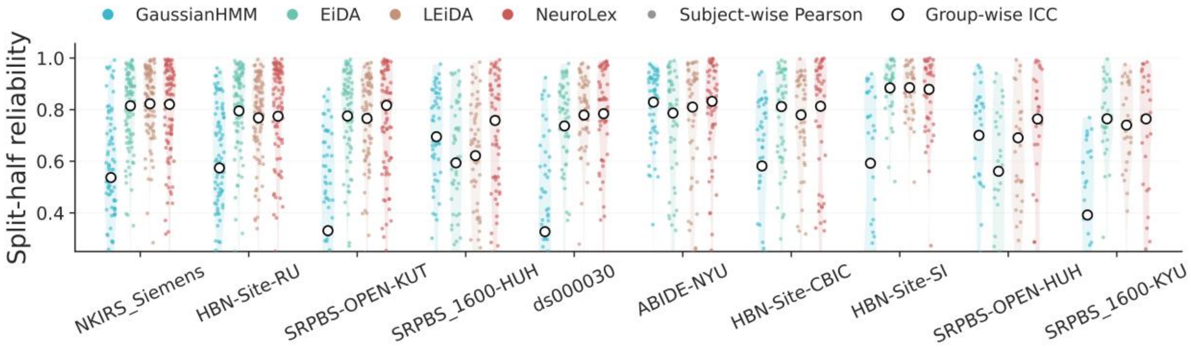

**Supplementary Fig. S4. Split-half reproducibility across ten independent scanning sites for NeuroLex and three baseline methods (GaussianHMM, EiDA, and LEiDA).** Each point represents the result from a single split. Subject-wise reproducibility is quantified using Pearson correlation coefficients (filled circles), and group-wise reliability is quantified using intraclass correlation coefficients (open circles). NeuroLex shows consistently higher reliability across sites at both the individual and group levels. Shaded density plots summarize the overall reliability distributions across all sites.

#### 8. Consistency and reliability of brain state clustering

We evaluated the cross-site consistency of brain state representations using the adjusted Rand index (ARI), which quantifies the agreement between clustering solutions across repeated runs at the index level. ARI scores were computed for NeuroLex and three baseline methods (GaussianHMM, EiDA, and LEiDA) across the largest ten independent scanning sites (Supplementary Fig. S5). Across sites, NeuroLex achieved higher ARI scores than the baseline methods in most cases, indicating more stable clustering assignments across runs. While ARI values varied across sites, reflecting differences in data characteristics and sample composition,

NeuroLex maintained relatively high and stable agreement across all sites. In contrast, GaussianHMM and EiDA exhibited lower ARI scores with greater dispersion, suggesting increased variability in clustering outcomes. LEiDA showed intermediate performance, with ARI scores higher than GaussianHMM and EiDA at several sites but generally lower than those obtained with NeuroLex.

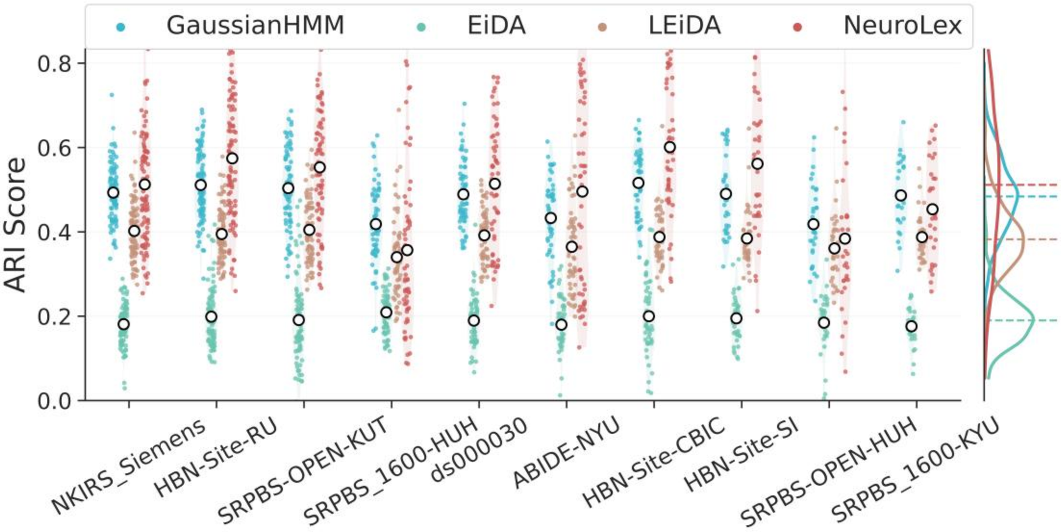

**Supplementary Fig. S5. Adjusted Rand index (ARI) scores across the largest ten independent scanning sites for NeuroLex and three baseline methods (GaussianHMM, EiDA, and LEiDA).** Each point represents the ARI from a single subject, with open circles indicating site-wise means. NeuroLex shows higher ARI scores in most sites, reflecting more stable clustering solutions across repeated runs. Density plots on the right summarize the overall ARI distributions across all sites for each method.

#### 9. Cross-run stability of brain-state token spatial patterns

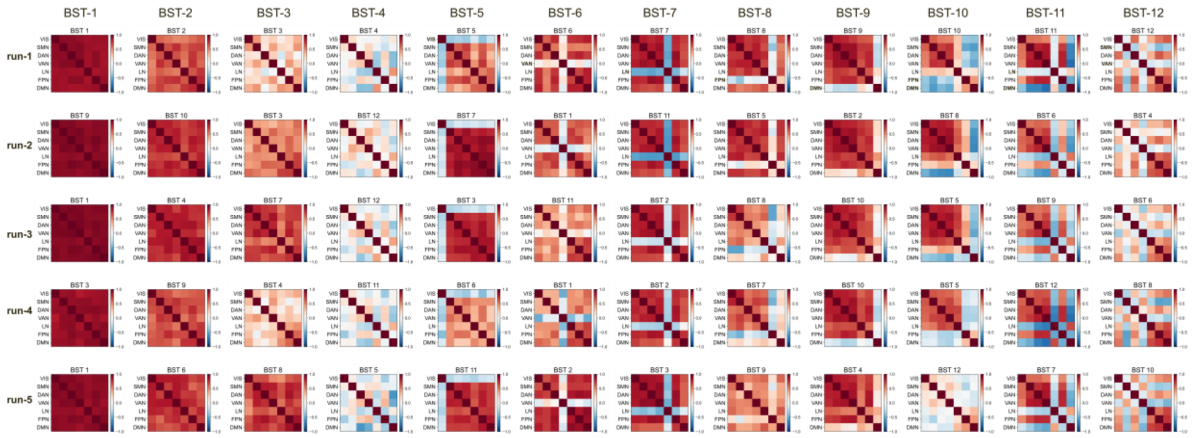

**Supplementary Fig. S6. Cross-run reproducibility of dynamic functional connectivity patterns underlying brain-state tokens.** Although deep latent embeddings are not strictly identical across independent runs, the emergent brain-state tokens preserved their semantic correspondence and exhibited a highly consistent organizational structure.

To examine the robustness of the learned brain-state tokens, we repeated the model with each time sampling 90% data for training using independent random initializations. It is well established that deep learning models operate within high-dimensional non-convex optimization landscapes, and thus the resulting latent embedding geometry is not strictly deterministic across runs. Variations in token centroids and embedding coordinates are therefore expected even under identical hyperparameters and training data. Despite this intrinsic variability at the latent representation level, visualization of the reconstructed dynamic functional connectivity matrices reveals a high degree of spatial concordance across runs (Supplementary Fig. S6). The overall topological structure of network coupling and decoupling patterns is preserved, suggesting that the model consistently converges toward similar dynamical motifs embedded within the data. Notably, states characterized by selective decoupling of a single subnetwork (BST5-BST9) demonstrate particularly strong cross-run stability. These configurations show highly reproducible spatial patterns across all five runs, with minimal qualitative variation in inter-network phase coupling structure. These findings indicate that while exact latent embeddings are not fully deterministic, the emergent brain-state tokens represent reproducible population-level dynamical configurations rather than run-specific artifacts. The consistency observed in the spatial organization of dFC patterns supports the robustness of the NeuroLex tokenization framework and reinforces its interpretability at the level of large-scale functional network dynamics.

#### 10. Generalization Performance of NeuroLex across the train and test set

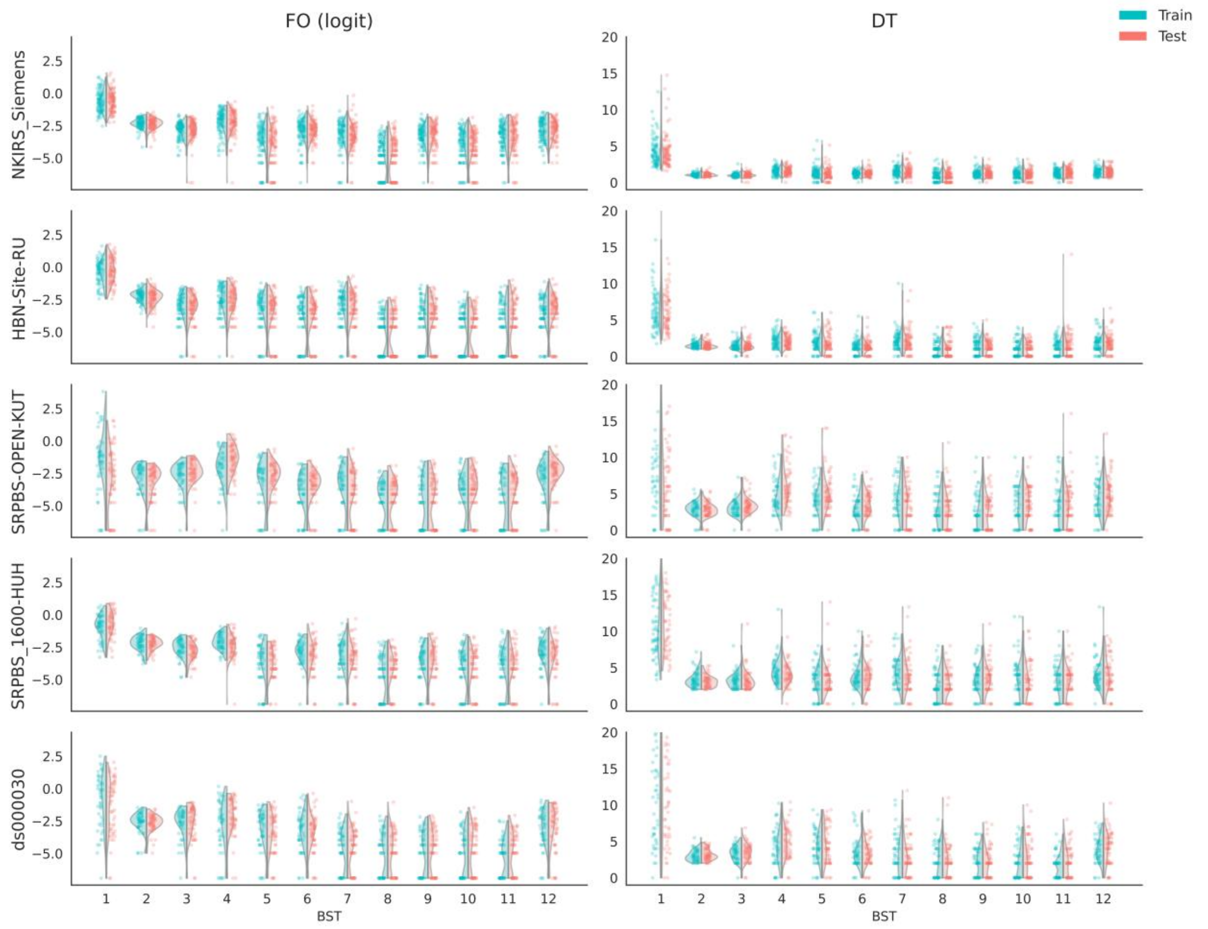

**Supplementary Fig. S7. Train-test generalization of brain state features across sites.** Violin plots show the distributions of fractional occupancy (FO, logit-transformed) and dwell time (DT) for each of the 12 brain state tokens across the largest five independent scanning sites. Training data are shown in green and healthy held-out test data in red. For each site and token, fractional occupancy and dwell time distributions in the test set closely match those in the training set, with no significant differences observed, indicating stable feature distributions across datasets and scanning sites.

To further assess the generalization ability of NeuroLex, we compared the distributions of two state expression features, fractional occupancy (FO) and dwell time (DT), across training and test datasets for each of the 12 BSTs and across the largest five scanning sites (Supplementary Fig. S7). For each site, token-wise fractional occupancy (logit-transformed) and dwell time distributions were visualized separately, with training samples shown in green and test samples shown in red. Across all sites and tokens, the distributions of fractional occupancy and dwell time in the test sets closely matched those observed in the corresponding training sets. No significant differences were detected between training and test distributions for either feature across the 12 brain state tokens, indicating that NeuroLex-derived representations generalize consistently across datasets acquired at different scanners.

#### 11. Generalization capability to internal test and external datasets

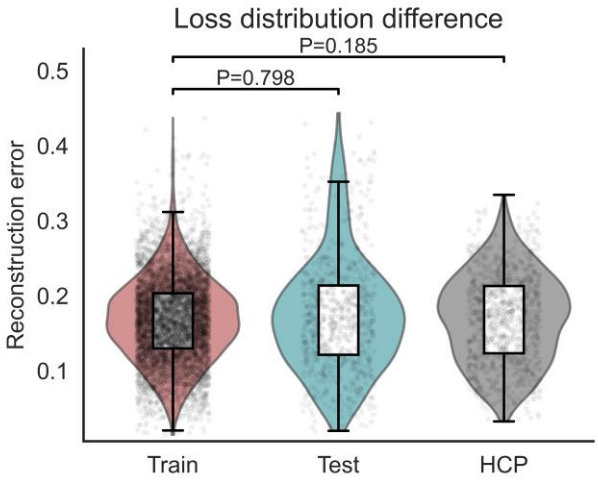

**Supplementary Fig. S8. Reconstruction error distributions across training, internal test, and external cohorts.** Reconstruction error distributions are shown for the training set, internal test set, and independent held-out HCP cohort. Error distributions were highly comparable across subsets, with no significant differences detected between Train and Test (Wilcoxon rank-sum test, P = 0.798) or between Train and HCP (P = 0.185).

To disentangle representation-level generalization from downstream site variability, we assessed model performance using reconstruction error. This metric directly evaluates whether the model overfits the training cohort and whether performance degrades in independent external datasets. We compared the distribution of reconstruction errors across the training set, internal test set, and an independent held-out HCP cohort. As shown in Supplementary Fig. S8, reconstruction error distributions were highly comparable across all three subsets. Nonparametric Wilcoxon rank-sum tests revealed no significant differences between Train and Test (P = 0.798) or between Train and HCP (P = 0.185). Importantly, no systematic inflation of reconstruction error was observed in either the internal test or the external HCP dataset. These findings indicate that the model does not exhibit detectable overfitting to the training cohort and maintains stable performance under cross-cohort evaluation. Collectively, this supports robust generalization of the learned representation beyond scanner- and site-specific characteristics.

#### 12. Evaluations of GAMLSS model specifications

We evaluated the robustness of the normative modeling results by comparing multiple GAMLSS specifications that differed in the parameterization of the scale component and the transformation of age. Across all runs, the location parameter ($\mu$) was specified identically, with the linear predictor given by:

$$\eta_\mu = f_{sex}(\log(age)) + \beta_{sex}[sex] + b_{site}$$

where $f_{sex}(\cdot)$ denotes a sex-stratified smooth function of age. Sex was included as fixed effects, and site was modeled as a random intercept. The smooth term for age was specified using penalized B-splines with a low degree of freedom, allowing for flexible yet stable modeling of age-related effects while avoiding overfitting. We set multiple configures for the fitting, whereas in configs 1–3, the scale parameter ($\sigma$) was specified as:

$$\text{Config-1: } \eta_\sigma = \gamma_0 + \gamma_1 \log(age)$$

$$\text{Config-2: } \eta_\sigma = \gamma_0$$

$$\text{Config-3: } \eta_\sigma = \gamma_0 + \gamma_1 \log(age) + \gamma_{sex}[sex]$$

, while the shape parameters were held constant across runs:

$$\eta_\nu = \delta_0, \eta_\tau = \kappa_0$$

The degrees of freedom and shape parameters had only a minor impact on the results, we therefore fixed them to reduce the risk of overfitting. Configs 4–6 mirrored Configs 1–3, respectively, with $\log(age)$ replaced by linear age in the corresponding terms of the $\sigma$ model.

The results are shown in Supplementary Table S4. Across all model variants, the resulting normative estimates were highly consistent. The empirical distributions of Z-scores remained centered near zero with comparable variance across runs, and higher-order moments (skewness and kurtosis) showed minimal variation. Model fit and predictive performance metrics, including log score, $R^2$, MAE, MSE, RMSE, AIC, and BIC, were also highly similar across specifications, indicating that the normative modeling results were robust to alternative parameterizations of $\sigma$ and to the choice of age transformation. In comparison, config-3 achieved the most favorable performance in average. Specifically, config-3 yielded Z-score distributions closest to the expected standard normal properties, with stable mean and variance and minimal higher-order deviations. In addition, config-3 showed consistently competitive or lower error metrics and information criteria compared with alternative specifications, while maintaining comparable goodness-of-fit across runs. Given its stable performance across a broad set of evaluation criteria, config-3 was selected as the final GAMLSS specification for all subsequent analyses.

#### 13. Normative trajectories of fractional occupancy across the lifespan

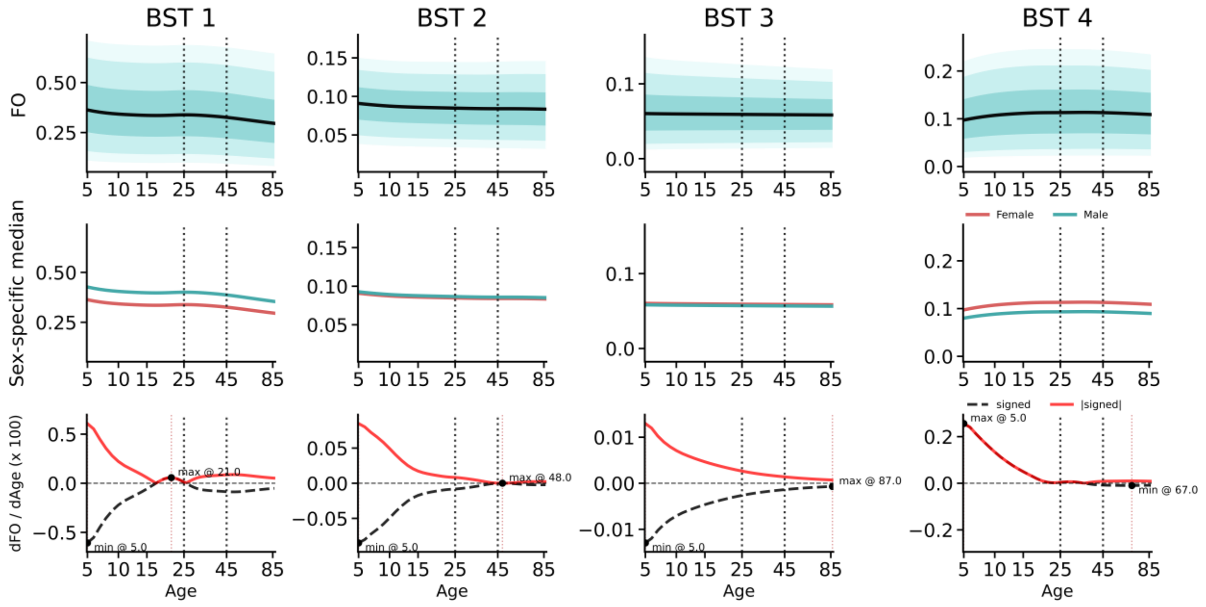

**Supplementary Fig. S9. Lifespan trajectories of fractional occupancy for BST 1–4.** The first row shows the age-dependent centile curves of fractional occupancy (FO) estimated using GAMLSS, including the 5th, 10th, 25th, 50th, 75th, 90th, and 95th percentiles. Shaded regions represent uncertainty around the estimated centiles. The second row depicts sex-specific median trajectories, with males shown in cyan and females in red. The third row illustrates the age-related gradient of FO, where the black curve represents the signed first-order derivative with respect to age (dFO/dAge), and the red curve denotes its absolute magnitude, highlighting periods of rapid change across the lifespan.

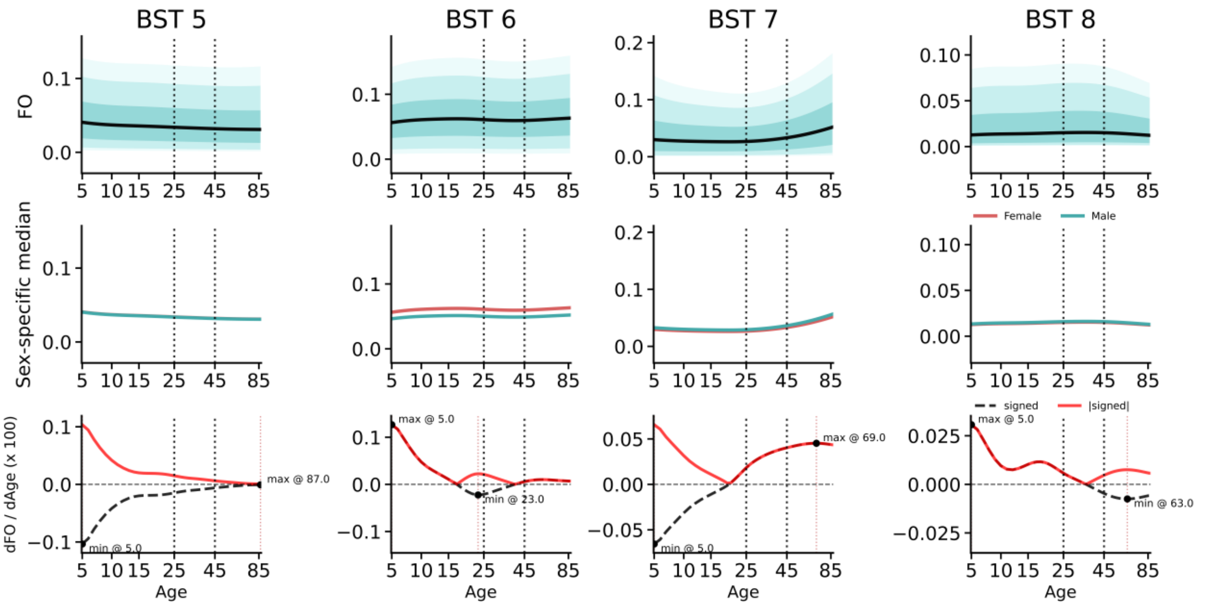

**Supplementary Fig. S10. Lifespan trajectories of brain-state fractional occupancy for BST 5–8.** Visualization follows the same convention as in Fig. S6, showing centile curves (top), sex-specific median trajectories (middle; cyan: male, red: female), and age-related gradients of fractional occupancy (bottom; black: signed gradient, red: absolute gradient).

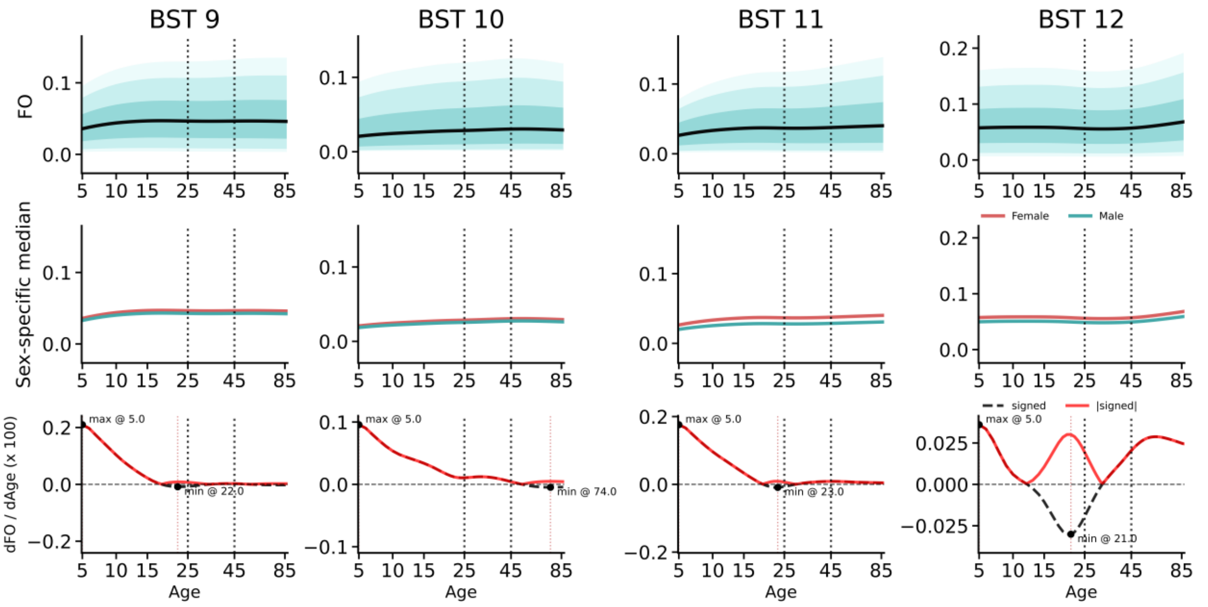

**Supplementary Fig. S11. Lifespan trajectories of brain-state fractional occupancy for BST 9–12.** Visualization follows the same convention as in Fig. S6, showing centile curves (top), sex-specific median trajectories (middle; cyan: male, red: female), and age-related gradients of fractional occupancy (bottom; black: signed gradient, red: absolute gradient).

Across the 12 brain-state tokens, we observed pronounced age-dependent changes in fractional occupancy, with many states exhibiting non-linear developmental trajectories. Notably, several dominant and functionally relevant states showed rapid changes during childhood and adolescence, followed by a relative stabilization in adulthood. Specifically, BST 1, BST 3, BST 4, BST 6, BST 9, and BST 11 displayed marked fractional occupancy changes before approximately 25 years of age, after which their trajectories gradually plateaued, indicating a transition from developmental reorganization to a more stable adult regime. In contrast, other states demonstrated comparatively flatter trajectories across the lifespan, suggesting more age-invariant engagement. Gradient analyses further highlighted that the magnitude of age-related change peaked predominantly in early life, with substantially reduced gradients beyond young adulthood, reinforcing the presence of a critical developmental window for large-scale brain-state reconfiguration. Clear sex-specific differences were observed in a subset of brain states. BST 1, BST 4, BST 6, BST 11, and BST 12 exhibited consistent separation between male and female median trajectories across substantial portions of the lifespan. These differences were most pronounced in early and mid-adulthood and persisted after accounting for site-related variability, suggesting systematic sex effects on the temporal allocation of these brain states rather than transient developmental noise. In contrast, other states showed largely overlapping male and female trajectories, indicating relative sex invariance.

#### 14. Normative trajectories of dwell time across the lifespan

Similar to fractional occupancy, dwell time exhibited pronounced age-dependent changes across multiple brain states, with the most prominent transitions occurring before early adulthood. Several states showed rapid dwell time modulation during childhood and adolescence, followed by a stabilization phase after approximately 25 years of age, indicating a shared developmental turning point in both state occurrence and temporal persistence. In contrast to fractional occupancy, dwell time demonstrated minimal sex-related differences except BST1. Male and female median trajectories largely overlapped in most brain states and age ranges. This suggests that while age strongly shapes the temporal persistence of brain states, sex effects primarily influence how frequently states are entered rather than how long they are maintained once engaged. Overall, fractional occupancy and dwell time capture complementary aspects of brain-state dynamics: fractional occupancy is more sensitive to sex-dependent differences in state engagement, whereas dwell time predominantly reflects age-related maturation and stabilization processes shared across sexes.

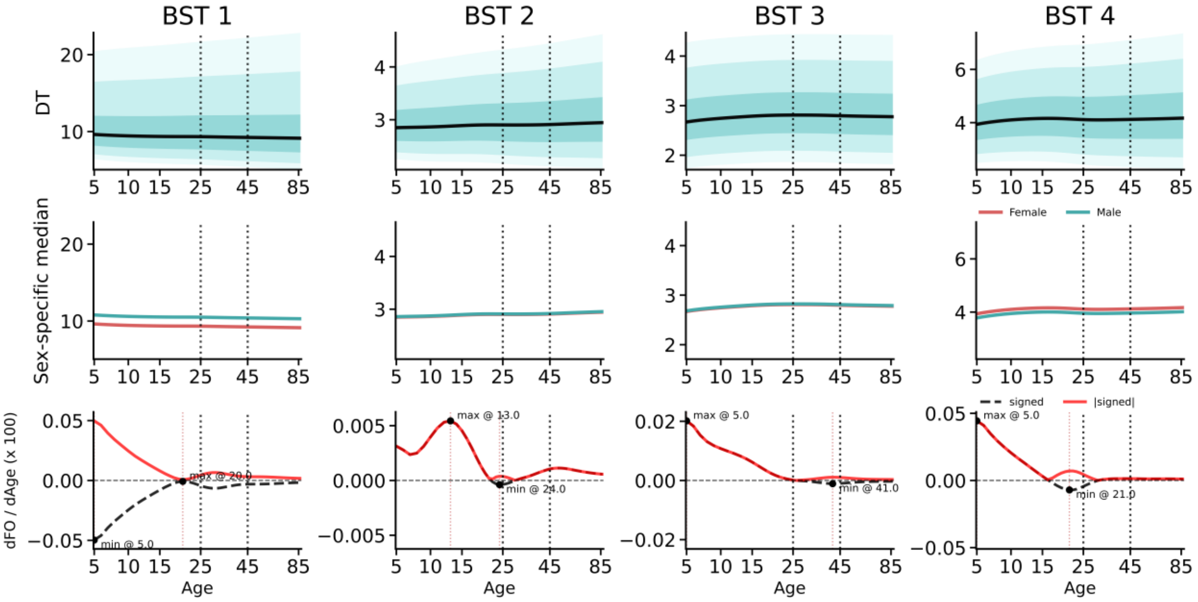

**Supplementary Fig. S12. Lifespan trajectories of dwell time for BST 1–4.** The first row shows the age-dependent centile curves of dwell time (DT) estimated using GAMLSS, including the 5th, 10th, 25th, 50th, 75th, 90th, and 95th percentiles. Shaded regions represent uncertainty around the estimated centiles. The second row depicts sex-specific median trajectories, with males shown in cyan and females in red. The third row illustrates the age-related gradient of DT, where the black curve represents the signed first-order derivative with respect to age (dDT/dAge), and the red curve denotes its absolute magnitude, highlighting periods of rapid change across the lifespan.

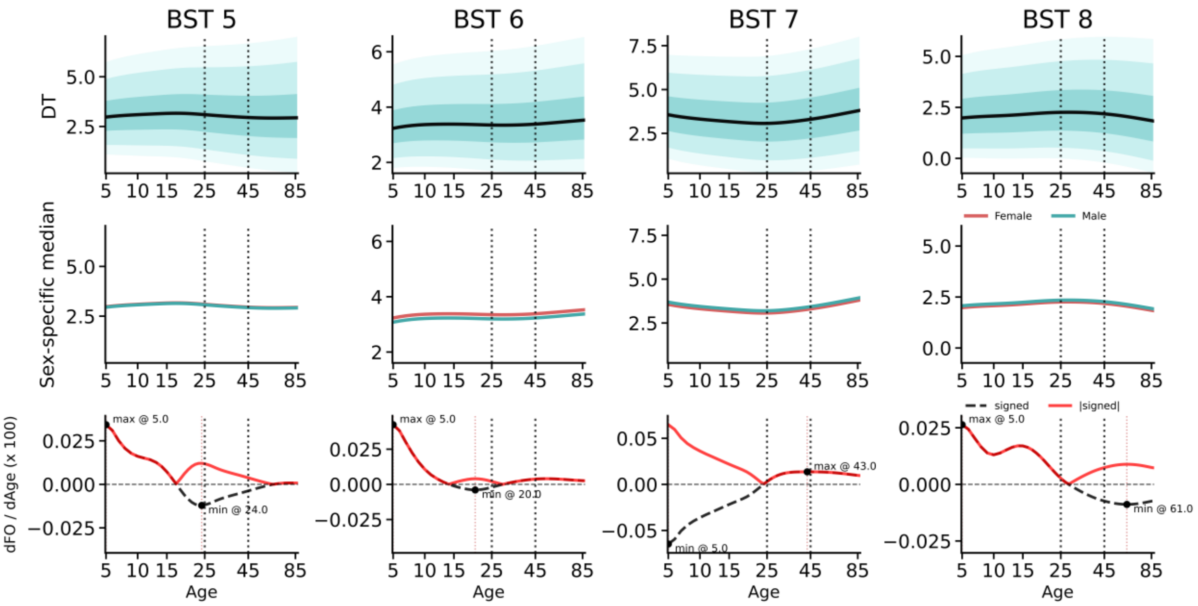

**Supplementary Fig. S13. Lifespan trajectories of brain-state dwell time for BST 5–8.** Visualization follows the same convention as in Fig. S9, showing centile curves (top), sex-specific median trajectories (middle; cyan: male, red: female), and age-related gradients of dwell time (bottom; black: signed gradient, red: absolute gradient).

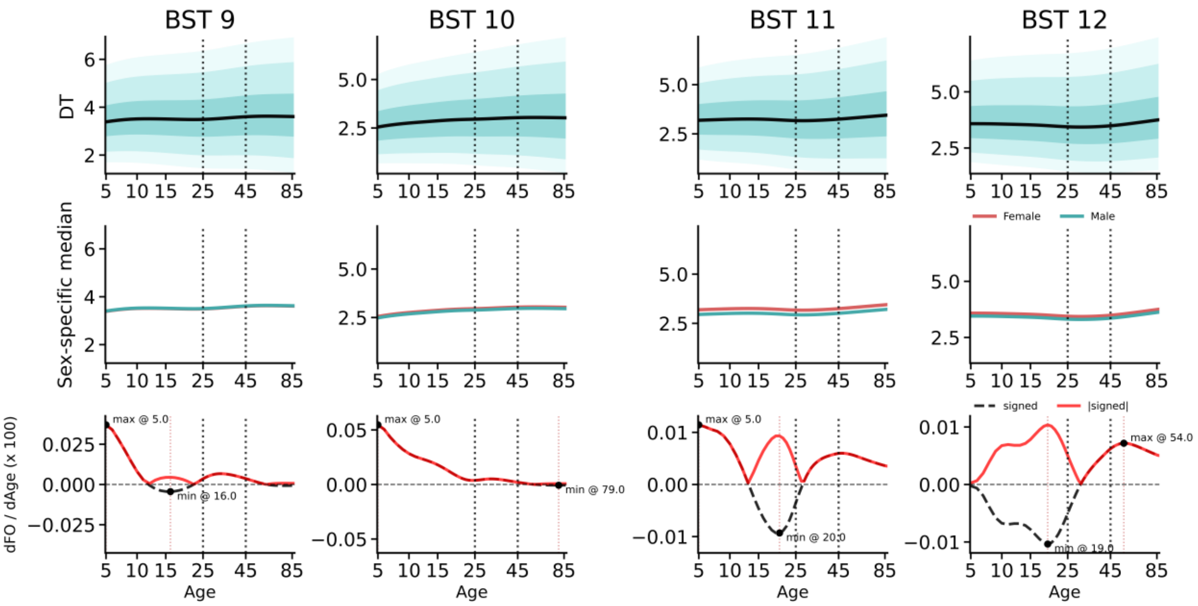

**Supplementary Fig. S14. Lifespan trajectories of brain-state dwell time for BST 9–12.** Visualization follows the same convention as in Fig. S9, showing centile curves (top), sex-specific median trajectories (middle; cyan: male, red: female), and age-related gradients of dwell time (bottom; black: signed gradient, red: absolute gradient).

#### 15. Structural normative trajectories across the lifespan

Using the same multi-site cohort as the functional analyses, we performed normative modeling of global structural brain measures to characterize their lifespan trajectories. Grey matter volume (GMV) showed a largely monotonic decline from early childhood onward, indicating that the sampled age range captures the post-peak phase of gray matter development. This pattern is consistent with previous lifespan studies demonstrating that total cortical gray matter volume reaches its maximum in early childhood, followed by sustained age-related cortical thinning. White matter volume (WMV) and subcortical gray matter volume (SGMV) exhibited a non-monotonic trajectory, characterized by an initial increase during childhood, a peak in late childhood to early adolescence, and a gradual decline across adulthood and aging. This prolonged maturational profile contrasts with the earlier maturation of GMV and reflects extended myelination and subcortical development processes. In contrast, ventricular cerebrospinal fluid (VEN) increased steadily across the lifespan, with an accelerated rise in later life, consistent with global brain tissue loss and ventricular expansion associated with aging. Sex-stratified trajectories revealed systematic differences in absolute volume levels between males and females, while preserving highly similar age-related trends across all structural measures. These results are consistent with previous studies[7] and form a baseline for comparison. Note however, that we also provide a tool that allows for transfer of GAMLSS models in a federated fashion, which has not been implemented before.

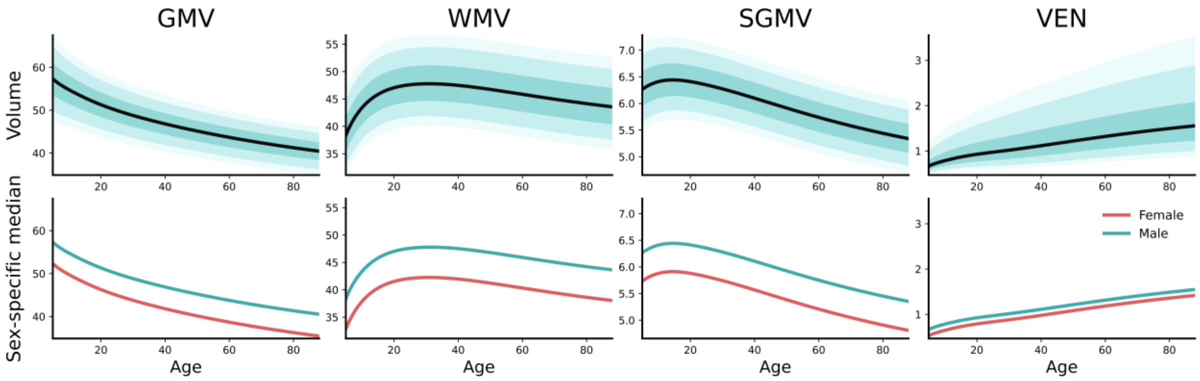

**Supplementary Fig. S15. Lifespan normative trajectories of structural brain measures.** Normative models were fitted to gray matter volume (GMV), white matter volume (WMV), subcortical gray matter volume (SGMV), and ventricle cerebrospinal fluid (VEN), using the same multi-site cohort as the fMRI analyses. Top row shows the population-level median trajectory (black line) with shaded bands indicating the estimated centile ranges. The bottom row depicts sex-specific median trajectories for females (red) and males (cyan). GMV exhibits a monotonic decline across the sampled age range, consistent with early childhood peak followed by progressive cortical thinning. In contrast, WMV and

SGMV show an early life increase with a peak in late childhood to early adolescence, followed by gradual decline in adulthood. ventricular volume increases steadily with age, reflecting age-related brain tissue loss and ventricular enlargement. All models account for site-related variability within the normative modeling framework.

#### 16. Functional connectivity normative trajectories across the lifespan

Moreover, we performed normative modeling of large-scale functional connectivity derived from the Yeo7 parcellation to characterize lifespan trajectories of static network-level interactions. For each participant, we quantified within-network and between-network connectivity across the seven canonical systems (visual, somatomotor, dorsal attention, ventral attention/salience, limbic, frontoparietal control, and default mode networks). Consistent with our brain dynamics findings, functional connectivity exhibited pronounced age-related changes prior to approximately 25 years, with the majority of connections showing a relatively rapid decline during late adolescence and early adulthood. Across much of the lifespan, connectivity patterns were either relatively stable or mildly decreasing from childhood onward, reflecting early functional maturation followed by progressive stabilization and subsequent attenuation. Notably, coupling between the limbic network and other large-scale systems demonstrated a marked age-related reduction. This pattern converges with our dynamic state analysis: BST7, characterized by limbic decoupling from the rest of the brain, showed increased fractional occupancy. The concordance between static connectivity and dynamic brain-state metrics therefore indicates a lifespan-wide decline in limbic integration, particularly evident in adulthood. In addition, at the global level, we observed an overall weakening of functional connectivity strength across networks. This trend parallels our BST1 findings, in which globally coupled brain states decreased in frequency, suggesting a progressive reduction in whole-brain integration. Sex-stratified trajectories further revealed systematic differences in connectivity strength across multiple networks. Compared with the dynamic metrics, static functional connectivity appeared more sensitive to sex-related effects, although the overarching age-related patterns were preserved in both males and females.

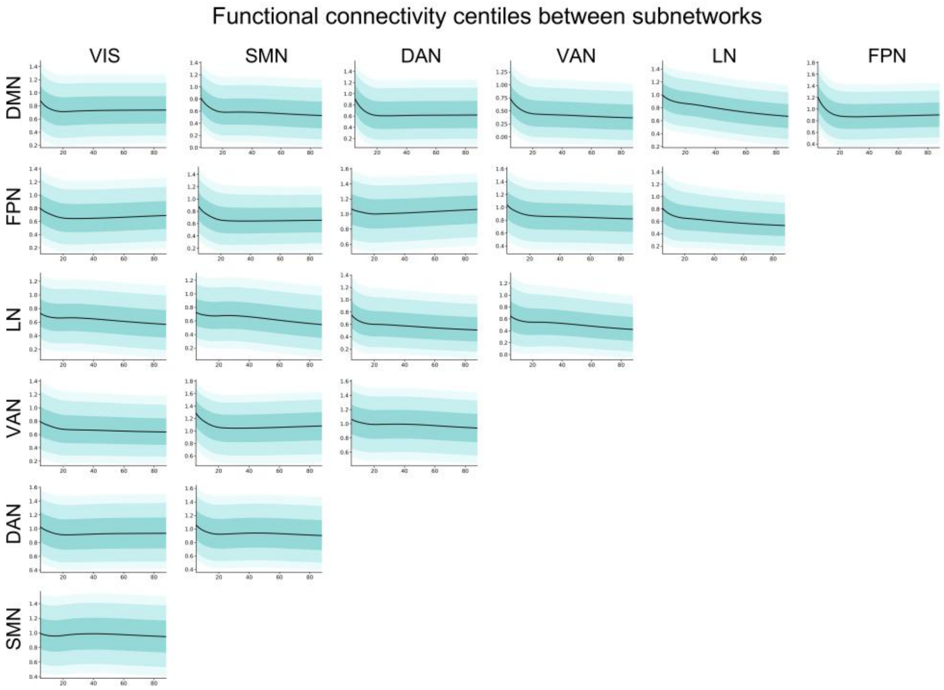

**Fig. S16. Lifespan normative trajectories of functional connectivity between subnetworks.** Normative models were fitted to the connectivity strength using the same multi-site cohort as the brain dynamics analyses. The plot shows the population-level median trajectory (black line) with shaded bands indicating the estimated centile ranges.

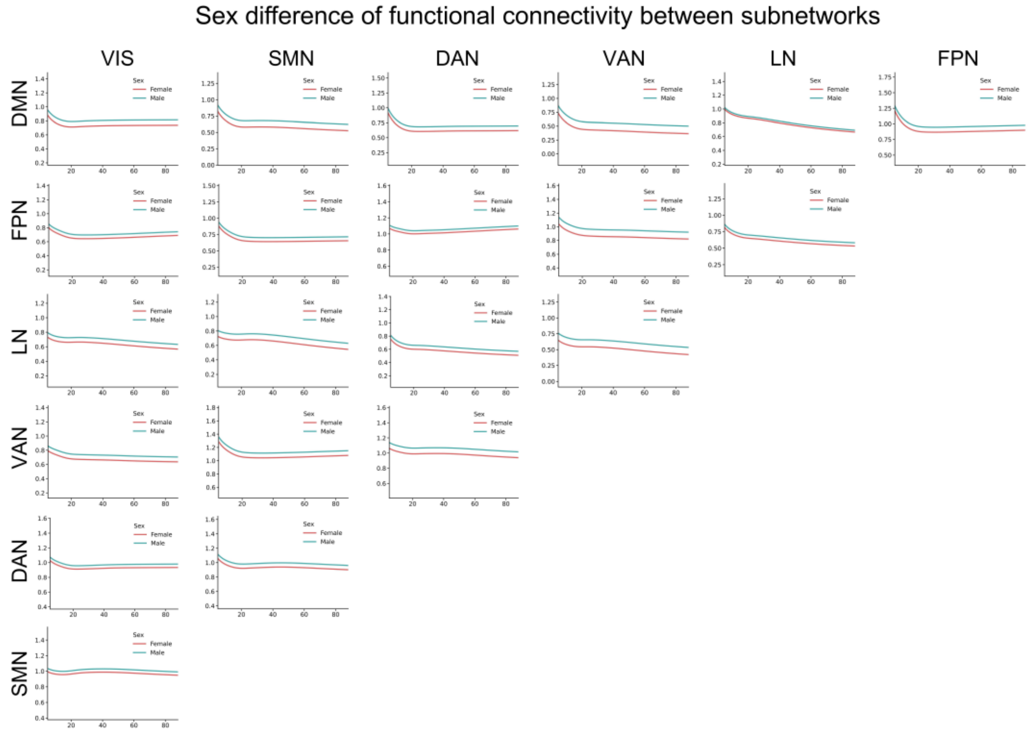

**Fig. S17. Lifespan normative trajectories of functional connectivity between subnetworks.** Normative models were fitted to the connectivity strength using the same multi-site cohort as the brain dynamics analyses. The plot shows sex-specific median trajectories for females (red) and males (cyan).

#### 17. Sensitivity Analysis of cross-cohort transfer sample size

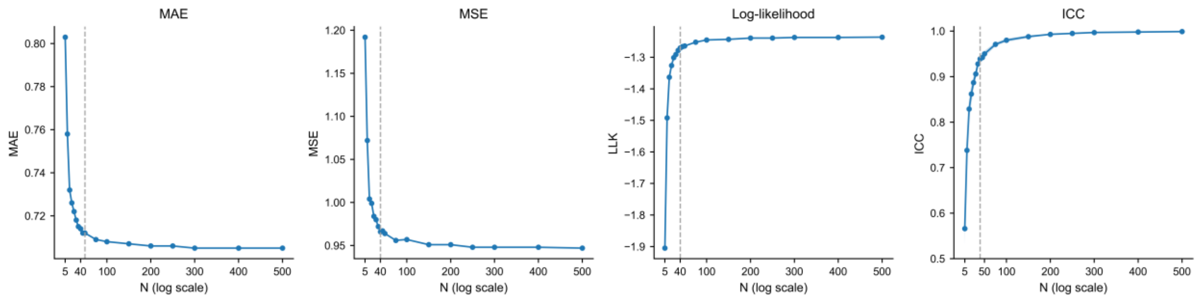

**Supplementary Fig. S18. Sample-size effects on cross-cohort transfer stability of the GAMLSS-based normative model.** Transfer stability was quantified using mean absolute error (MAE), mean squared error (MSE), log-likelihood (Logarithmic Score), and intraclass correlation coefficient (ICC).

To quantify the sample-size requirements for cross-cohort transfer of the GAMLSS-based normative framework, we conducted a controlled calibration analysis in the HCP1200 cohort (n = 1,012). An independent subset of 400 participants was fixed as a stable evaluation set, while the remaining 612 individuals served as the calibration pool. From this pool, we repeatedly subsampled varying numbers of participants to estimate study-specific location offsets ($\mu$) while keeping the distributional parameters otherwise fixed. For each candidate sample size, calibration subsets were bootstrap-sampled and the full transfer procedure was repeated 50 times to characterize variability and robustness across runs. Model performance in held-out healthy participants was quantified using mean absolute error (MAE), mean squared error (MSE), the Logarithmic Score (log-likelihood), and intraclass correlation coefficient (ICC) between observed values and predicted conditional means (μ). As illustrated in Supplementary Fig. S18, MAE and MSE exhibited a monotonic decrease with increasing calibration sample size, with a pronounced inflection around ~40 samples, beyond which improvements progressively attenuated, approaching an asymptotic regime. In parallel, both Logarithmic Score and ICC increased with sample size and demonstrated clear stabilization beyond approximately 40 participants. Notably, ICC exceeded 0.950 at 45 calibration samples and reached 0.980 at 100 samples, indicating near-ideal agreement between calibrated predictions and empirical observations. Overall, these results suggest that, under high-quality acquisition conditions such as HCP1200, approximately 40 participants are sufficient to achieve stable offset recalibration for cross-cohort transfer, whereas ~100 samples provide near-optimal calibration performance with minimal residual variability.

#### 18. Cross-cohort transferability of normative models

Violin plots show the distributions of normative z-scores for each brain state token across the reference cohort, the test healthy controls (TestSet), and two independent held-out external datasets (HCP1200 and ABCD), assessing the transferability of our norms out-of-distribution. Each point represents an individual. Black horizontal bars indicate the median z-score of each cohort, and the dashed line denotes zero deviation from the normative reference. Overall, z-score distributions are centered around zero across cohorts, indicating stable transfer of the normative model to unseen healthy populations. A small number of brain state show weak while marginally significant shifts compared to the reference set (asterisk), which likely reflect partial site mismatch and increased sampling variability.

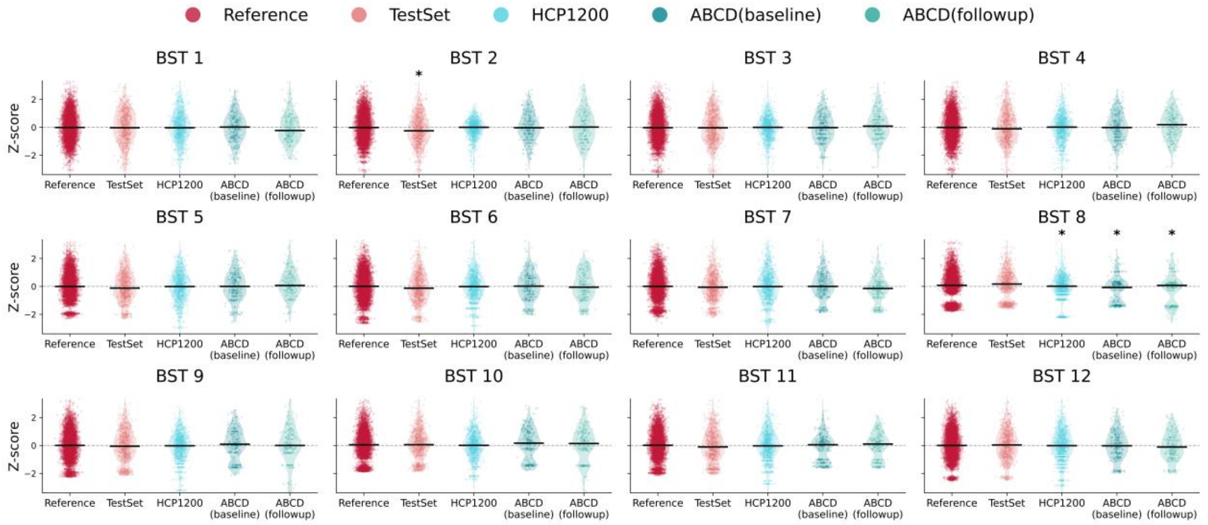

**Supplementary Fig. S19. Z-score distributions of fractional occupancy comparing the reference cohort, the internal test set, and site-transfer evaluations on HCP1200 and ABCD test cohorts.**

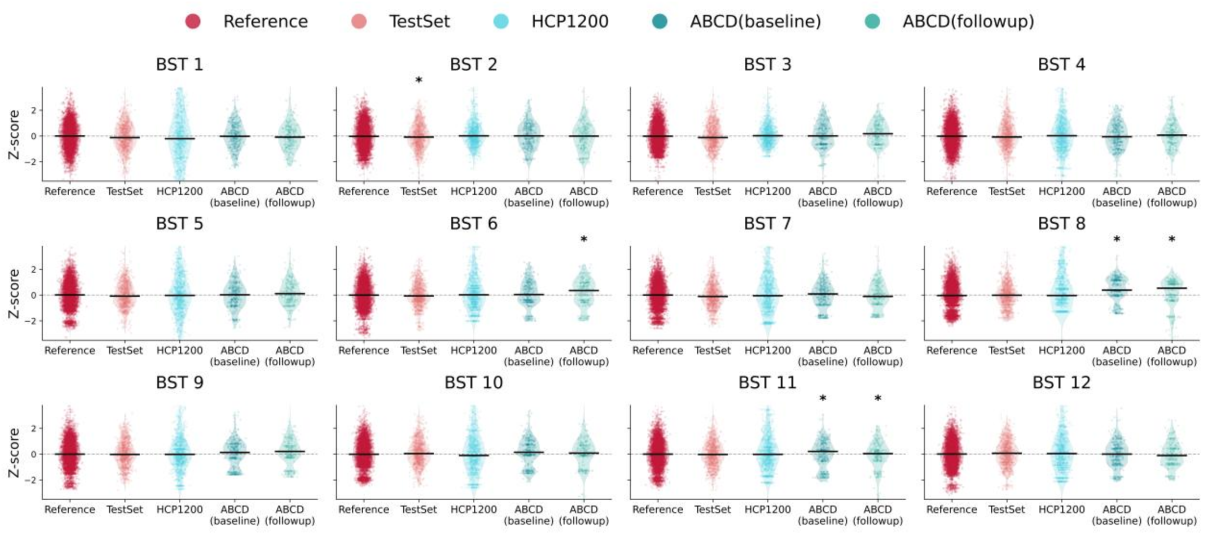

**Supplementary Fig. S20. Z-score distributions of dwell time comparing the reference cohort, the internal test set, and site-transfer evaluations on HCP1200 and ABCD test cohorts.**

#### 19. Application to the ABCD cohort and longitudinal slope analysis

We further implemented a transfer procedure when applying the normative model to the held-out ABCD cohort. Because the ABCD acquisition was performed with a repetition time (TR) of 0.8 s, time series were downsampled to 2 s to ensure harmonization with the reference model. The ABCD healthy-control cohort comprised 367 individuals and 589 scans (mean scans per subject = 1.6). For transfer calibration, half of the baseline scans were randomly selected to estimate study-specific location offsets, ensuring that each site contributed at least 40 scans for calibration; the remaining scans were retained for independent testing. For longitudinal slope analyses, change was quantified as $\Delta z/\Delta age$ for each subject, and group differences were assessed using two-sided Mann–Whitney U tests.

Supplementary Figs. S21–S22 present the slope-based longitudinal analyses in ADHD, BP, and OCD. Red indicates a positive Cliff's δ effect size (greater increase in $\Delta z/\Delta age$), whereas green indicates a relative decrease. Brain-state tokens reaching statistical significance in either fractional occupancy or dwell time are marked with an asterisk. Owing to limited sample sizes, only ADHD demonstrated a statistically significant alteration after multiple-comparison correction (corrected $P < 0.05$), specifically in the dwell time of BST9. Other tokens showed marginal significance, yet several exhibited effect sizes approaching 0.2. Notably, the ADHD slope patterns were broadly consistent with the cross-sectional findings in Fig. 4, with decreasing dwell time across multiple states (BST6, BST8, BST9, BST10, BST11) and a relative increase in fractional occupancy of BST7, indicating a progressive shift toward reduced temporal persistence and altered state occupancy, consistent with a tendency toward more labile and rapidly reconfiguring large-scale network dynamics in ADHD. In contrast, BP and OCD did not exhibit robust, convergent longitudinal effects and instead showed heterogeneous patterns of change. BP exhibited heterogeneous, state-specific trajectories, with changes primarily involving BST5 as well as BST3 and BST6, without evidence for a coherent global shift in dwell time across tokens. In OCD, longitudinal alterations were most apparent in the VAN-decoupling state, suggesting progressive modulation of ventral attention–related interactions. Given the role of the ventral attention network in salience detection and stimulus-driven reorienting, such VAN-linked dynamical shifts may reflect evolving imbalances between externally driven attentional capture and internal control processes that are central to obsessive–compulsive symptomatology.

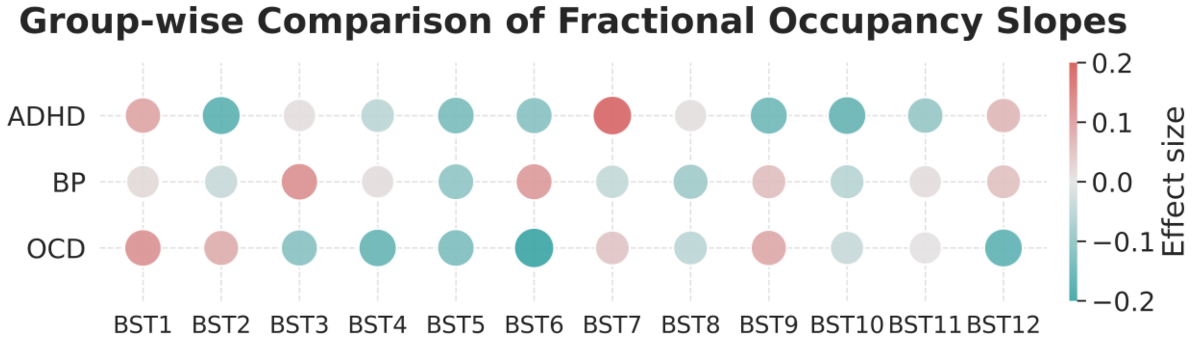

**Supplementary Fig. S21. Z-score distributions of dwell time comparing the reference cohort, the internal test set, and site-transfer evaluations on HCP1200 and ABCD test cohorts.**

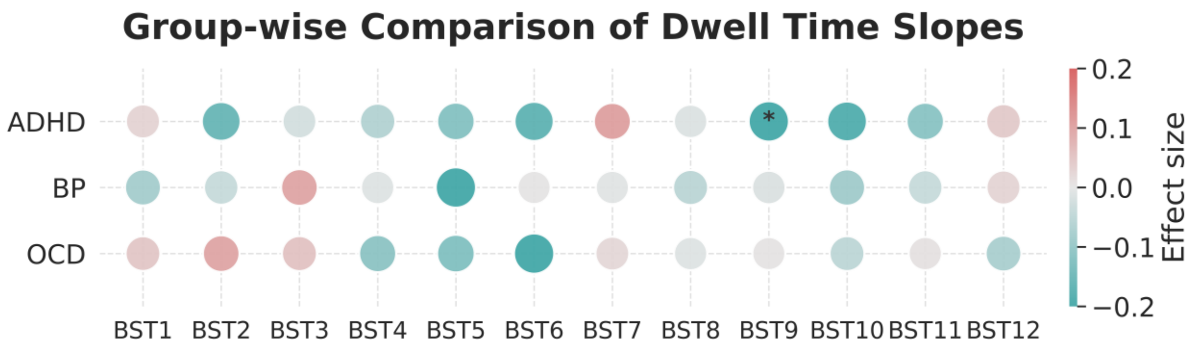

**Supplementary Fig. S22. Z-score distributions of dwell time comparing the reference cohort, the internal test set, and site-transfer evaluations on HCP1200 and ABCD test cohorts.**

#### 20. Atlas-level comparison with the Yeo-17 network parcellation

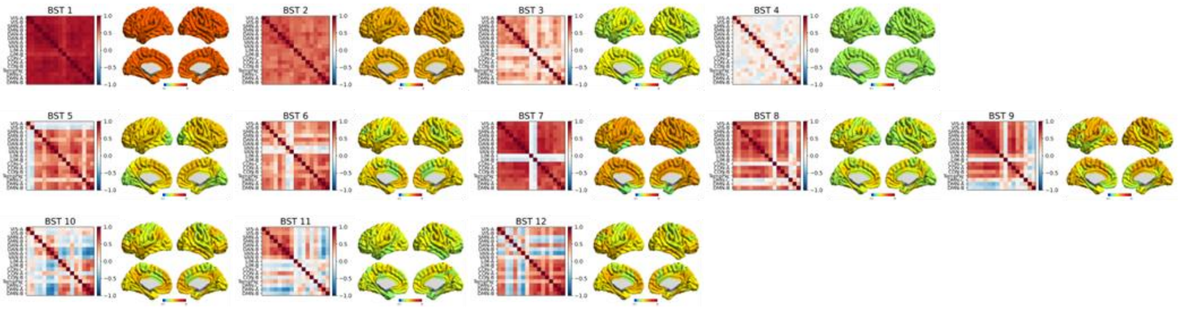

**Supplementary Fig. S23 Characteristic brain state patterns identified by NeuroLex in terms of Yeo-17.** For each state, the phase-lock matrix and corresponding spatial patterns are shown, together with global phase-locking strength.

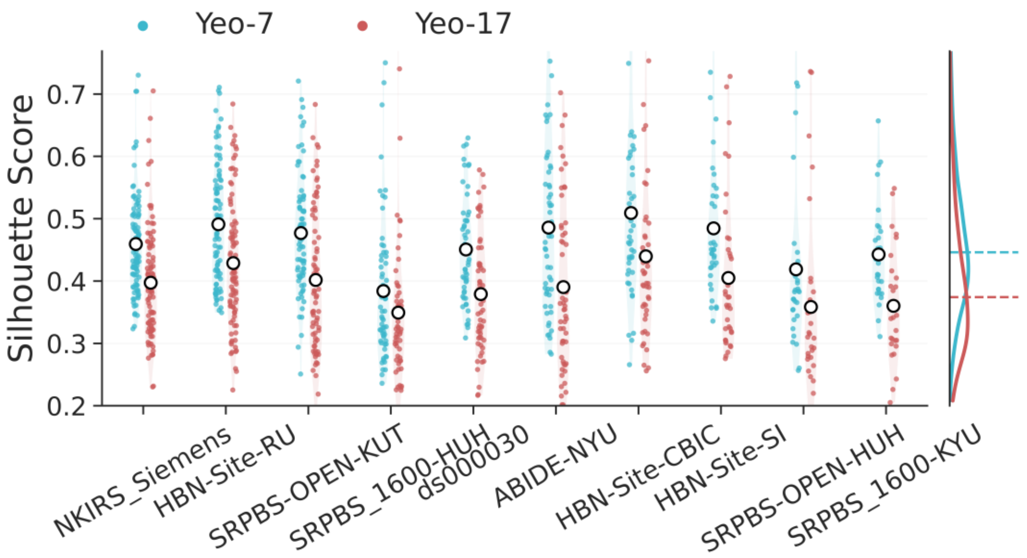

**Supplementary Fig. S24. Comparison of Yeo7 and Yeo17 in terms of Silhouette score across the largest 10 test scanner sites.**

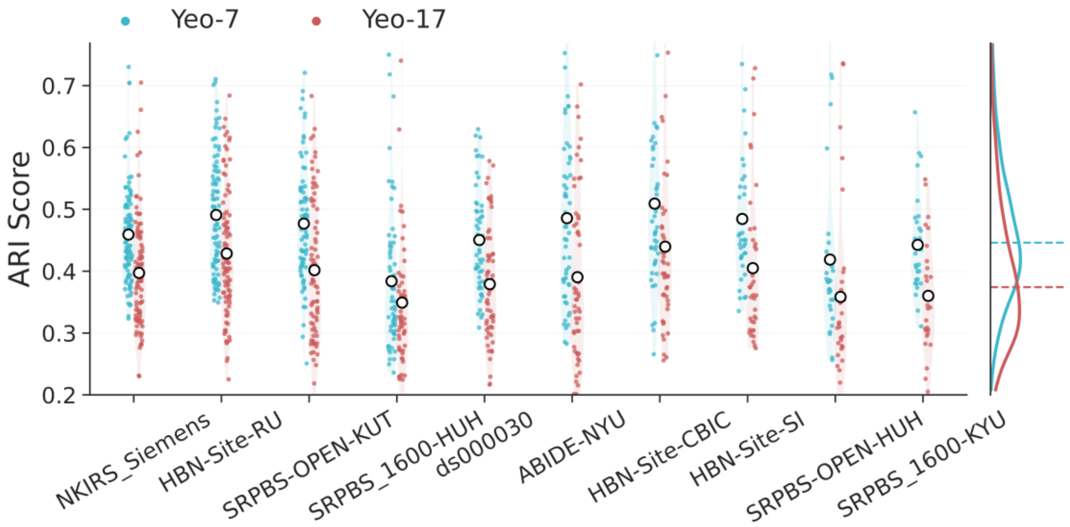

**Supplementary Fig. S25. Comparison of Yeo7 and Yeo17 in terms of ARI score across the largest 10 test scanner sites.**

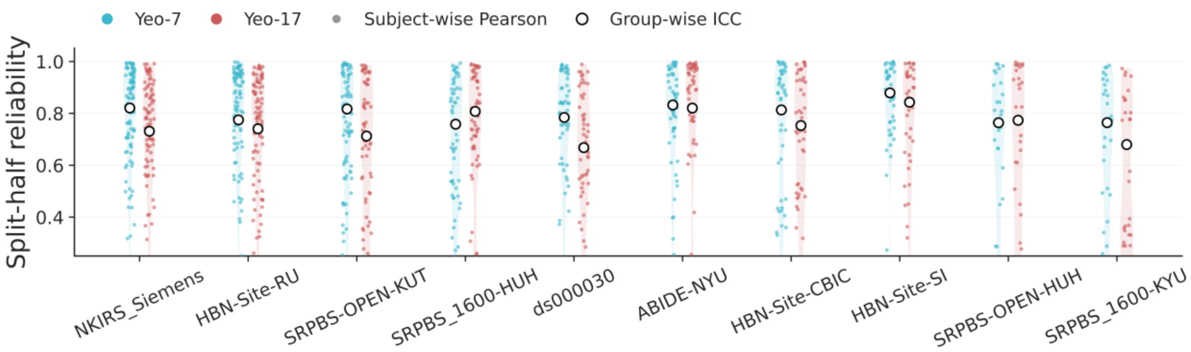

**Supplementary Fig. S26. Comparison of Yeo7 and Yeo17 in terms of Split-half reliability score across the largest 10 test scanner sites.**

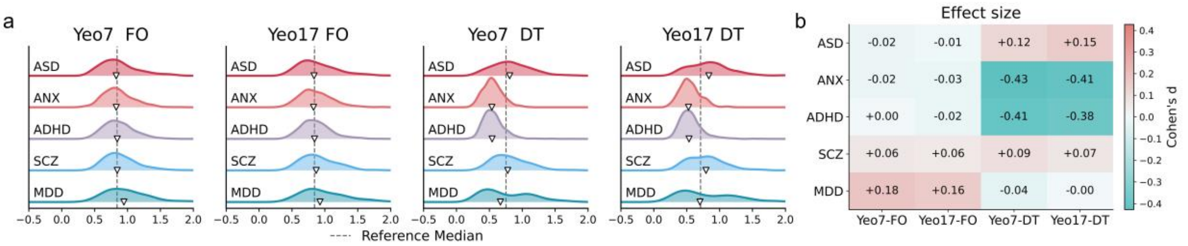

**Supplementary Fig. S27. Comparison of Yeo7 and Yeo17 in terms of composite z-score across five diagnostic groups.** a. Distribution of composite z-score of fractional occupancy and dwell time. b. Effect size (cliff's delta) comparing each disease group against the healthy control of the test set.

Supplementary Fig. S23 illustrates the spatial distributions of the learned brain-state tokens under the Yeo-17 network parcellation. By visual inspection, these patterns show a high degree of concordance with those reported in Fig. 2b, indicating that the learned tokens are robust to the choice of parcellation scheme. Consistent with the main results, we observe a gradual transition from globally highly coupled states to more weakly coupled configurations across BST1 to BST4. In contrast, BST5 to BST9 are characterized by decoupling patterns centered on specific core functional networks, reflecting increased functional segregation. Finally, BST10 to BST12 exhibit mixed or transitional configurations, combining features of multiple networks and resembling intermediate brain states. Notably, BST8 and BST9 show differences between the Yeo-7 and Yeo-17 representations. While these states primarily correspond to single-network decoupling patterns (BST8: FPN; BST9: DMN under Yeo-7), additional involvement of the default mode or limbic networks emerges under the finer Yeo-17 parcellation. We suggest that this effect is also caused by the hierarchical organization of large-scale brain networks. Finer parcellations resolve sub-network structure and inter-network boundaries more explicitly, which can lead to partial overlap between spatial patterns across closely related networks rather than indicating fundamentally distinct brain states. Despite these minor variations, the overall spatial motifs and their functional interpretations remain highly consistent across parcellations. Across more than 10,000 individuals, these findings support the existence of reproducible and population-level brain-state tokens, indicating a set of broadly shared and generalizable brain dynamic states across individuals and datasets.

Supplementary Figs. S24–S26 further demonstrates the quantitative results between Yeo-& and Yeo-17. Across atlas resolutions, we observed a systematic reduction in representation-level consistency when moving from Yeo7 to Yeo17, as reflected by lower Silhouette scores, ARI, and split-half reliability across the largest 10 test sites. This trend is caused as increasing the number of parcels introduces greater pattern uncertainty and reduces the stability of token-to-network correspondence. Coarse-grained functional systems can be further subdivided into multiple finer-scale configurations. Under higher-resolution parcellations, the number of plausible hierarchical decompositions increases, which in turn leads to greater ambiguity in token assignment and reduced agreement metrics. For this reason, we adopted the Yeo7 parcellation as the primary atlas in the main analyses, balancing interpretability at the systems level with improved numerical stability and cross-site robustness.

Despite modest changes in representation-level agreement across atlas resolutions, downstream disease sensitivity remained highly stable. Supplementary Fig. S27 summarizes diagnostic sensitivity derived from Yeo7- and Yeo17-based representations using high-level normative deviation metrics quantified by a composite |Z| score. To minimize potential confounding from site effects in cross-atlas comparison, we computed the composite deviation as the L1-norm mean of absolute z-scores rather than centile-based Mahalanobis distance[8], to overcome site-induced biases arising from differences in the covariance structure underlying the Mahalanobis metric. In Fig. S27a, the distribution of composite |Z| across diagnostic groups is shown, with dashed lines indicating the reference median and lower-triangular markers denoting group-specific medians to facilitate direct comparison of deviation magnitude. Fig. S27b presents the corresponding Cliff's δ effect sizes computed relative to matched test healthy controls. The results further confirm cross-resolution consistency, as disease-specific deviation patterns were preserved between Yeo7 and Yeo17 parcellations, with only minor numerical variations.

#### 21. Effects of temporal downsampling on brain state estimation

To examine whether interpolation involved in temporal downsampling influences the estimation of brain-state dynamics, we conducted a comparative analysis using the largest two independent resting-state fMRI datasets with different native temporal resolutions: SRPBS-OPEN (raw TR = 1.0 s) and NKIRS (raw TR = 0.64 s). For each dataset, we contrasted an interpolation-based downsampling strategy with a stride-based subsampling approach that retained every second time point without interpolation. We then assessed the impact of these two approaches on key dynamic features, including fractional occupancy and dwell time, across the twelve brain-state tokens.

Across both datasets, the two downsampling strategies yielded highly similar distributional profiles for fractional occupancy and dwell time. For each brain-state token, the overall shapes of the distributions and their central tendencies were largely preserved between interpolation-based and non-interpolated stride-based downsampling. This was consistently observed across different BSTs, suggesting that the inclusion of interpolation during downsampling does not systematically alter the relative prevalence or temporal persistence of the inferred brain states. Despite the difference in native temporal resolution between SRPBS-OPEN and NKIRS, the qualitative agreement between the interpolated and non-interpolated downsampling approaches was consistent across sites. Any observed differences were small, unsystematic, and did not follow a coherent direction across brain-state tokens or dynamic measures. These results indicate that the estimated brain-state tokens and their associated dynamic features are largely insensitive to whether temporal downsampling is performed with or without interpolation, supporting the robustness and generalizability of the proposed brain-state modeling framework across datasets with different acquisition parameters. Another possible suspect is that the phase-locking values may partially mitigate the impact of limited temporal resolution, as they capture relative phase relationships rather than relying strongly on instantaneous signal amplitudes.

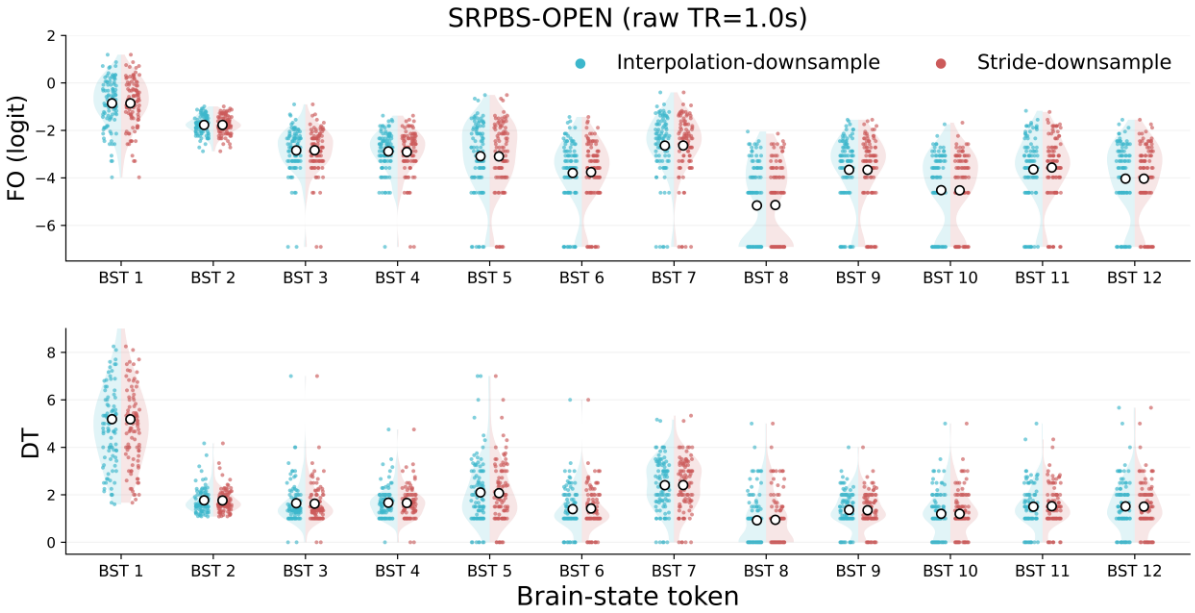

**Supplementary Fig. S28. Comparison of interpolation-based and stride-based downsampling on brain-state dynamics in terms of SRPBS-OPEN scanning site.** Distributions of fractional occupancy (FO, top) and dwell time (DT, bottom) are shown for the twelve brain-state tokens (BST1–BST12) estimated from the SRPBS-OPEN dataset (raw TR = 1.0 s). For each brain-state token, split violin plots summarize the distributions obtained using two different temporal downsampling strategies: interpolation-based downsampling (cyan, left half) and stride-based downsampling by retaining every second timepoint (red, right half). Individual dots represent subject-level estimates, while white circles with black outlines denote the mean value for each condition.

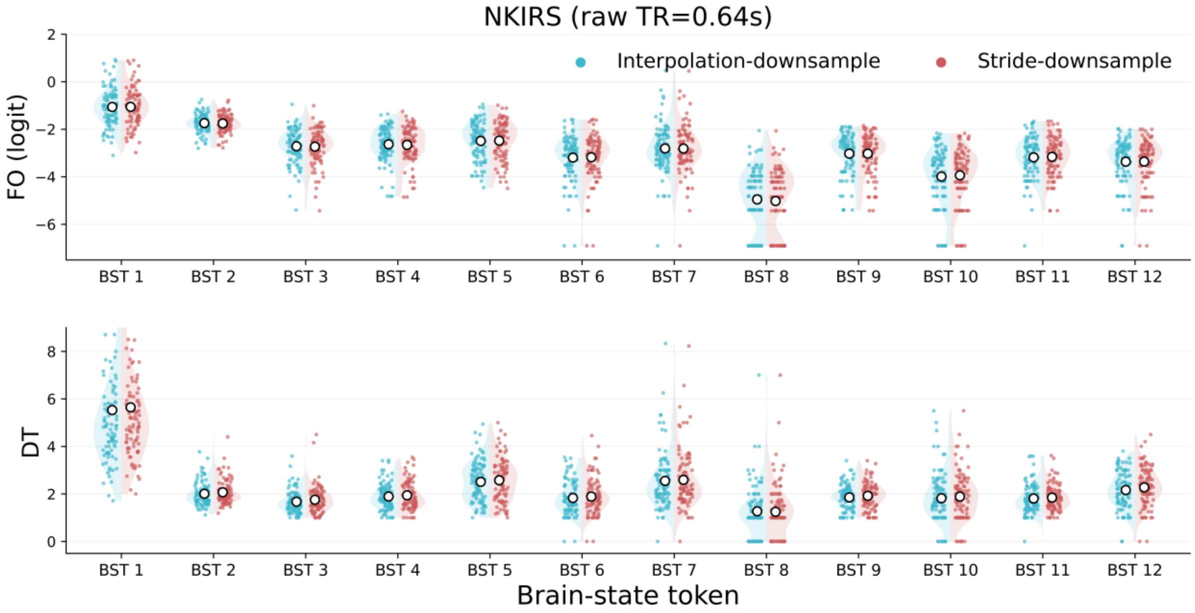

**Supplementary Fig. S29. Comparison of interpolation-based and stride-based downsampling on brain-state dynamics in terms of NKIRS scanning site.** Distributions of fractional occupancy (FO, top) and dwell time (DT, bottom) are shown for the twelve brain-state tokens (BST1–BST12) estimated from the NKIRS dataset (raw TR = 0.64 s). For each brain-state token, split violin plots summarize the distributions obtained using two different temporal downsampling strategies: interpolation-based downsampling (cyan, left half) and stride-based downsampling by retaining every three timepoint (red, right half). Individual dots represent subject-level estimates, while white circles with black outlines denote the mean value for each condition.

### Supplementary Tables

#### 22. Supplementary Table 1. Sources of all the studies used in the study

| Datasets | Sources | Comments | References |
|---|---|---|---|
| Autism Brain Imaging Dataset Exchange (ABIDE-1) | http://fcon_1000.projects.nitrc.org/ | Primary support for the work by Adriana Di Martino was provided by the NIMH (K23MH087770) and the Leon Levy Foundation. Primary support for the work by Michael P. Milham and the INDI team was provided by gifts from Joseph P. Healy and the Stavros Niarchos Foundation to the Child Mind Institute, as well as by an NIMH award to MPM (R03MH096321). | [9] |
| Autism Brain Imaging Dataset Exchange II (ABIDE-2) | http://fcon_1000.projects.nitrc.org/ | Primary support for the work by Adriana Di Martino and her team was provided by the National Institute of Mental Health (NIMH 5R21MH107045). Primary support for the work by Michael P. Milham and his team provided by the National Institute of Mental Health (NIMH 5R21MH107045); Nathan S. Kline Institute of Psychiatric Research). Additional Support was provided by gifts from Joseph P. Healey, Phyllis Green and Randolph Cowen to the Child Mind Institute. | |
| Cam-CAN | https://camcan-archive.mrc-cbu.cam.ac.uk/dataaccess/ | Data collection and sharing for this project was provided by the Cambridge Centre for Ageing and Neuroscience (CamCAN). CamCAN funding was provided by the UK Biotechnology and Biological Sciences Research Council (grant number BB/H008217/1), together with support from the UK Medical Research Council and University of Cambridge, UK. | [10,11] |
| Consortium for Reliability and Reproducibility (CORR) | http://fcon_1000.projects.nitrc.org/ | The National Institute on Drug Abuse (NIDA) and the National Natural Science Foundation of China (NSFC) have been instrumental in the CoRR collaboration providing the necessary funding and manpower to build the foundation of the project along with the Child Mind Institute, the Institute of Psychology, Chinese Academy of Sciences and the Nathan Kline Institute. | [12] |
| FCON1000 | https://fcon_1000.projects.nitrc.org/fcpClassic/FcpTable.html | This study used data from the 1000 Functional Connectomes Project (FCP). We thank the investigators and participants for making these data publicly available. | [13] |
| Healthy Brain Network (HBN) | https://fcon_1000.projects.nitrc.org/indi/cmi_healthy_brain_network/ | The Healthy Brain Network and its collaborative initiatives are supported by philanthropic contributions from the following individuals, foundations and organizations: Margaret Bilotti; Brooklyn Nets; Agapi and Bruce Burkard; James Chang; Phyllis Green and Randolph Cowen; Grieve Family Fund; Susan Miller and Byron Grote; Sarah and Geoff Gund; George Hall; Jonathan M. Harris Family Foundation; Joseph P. Healey; The Hearst Foundations; Eve and Ross Jaffe; Howard & Irene Levine Family Foundation; Rachael and Marshall Levine; George and Nitzia Logothetis; Christine and Richard Mack; Julie Minskoff; Valerie Mnuchin; Morgan Stanley Foundation; Amy and John Phelan; Roberts Family Foundation; Jim and Linda Robinson Foundation, Inc.; Linda and Richard Schaps; Zibby Schwarzman; Abigail Pogrebin and David Shapiro; Stavros Niarchos Foundation; Preethi Krishna and Ram Sundaram; Amy and John Weinberg; Donors to the 2013 Child Advocacy Award Dinner Auction; Donors to the 2012 Brant Art Auction. | [14] |

| | | | |
|---|---|---|---|
| NKIRS | https://fcon_1000.projects.nitrc.org/indi/enhanced/index.html | Principal support for the enhanced NKI-RS project is provided by the NIMH BRAINS R01MH094639-01 (PI Milham). Funding for key personnel also provided in part by the New York State Office of Mental Health and Research Foundation for Mental Hygiene. Funding for the decompression and augmentation of administrative and phenotypic protocols provided by a grant from the Child Mind Institute (1FDN2012-1). Additional personnel support provided by the Center for the Developing Brain at the Child Mind Institute, as well as NIMH R01MH081218, R01MH083246, and R21MH084126. Project support also provided by the NKI Center for Advanced Brain Imaging (CABI), the Brain Research Foundation, and the Stavros Niarchos Foundation. | [15,16] |
| SALD | https://fcon_1000.projects.nitrc.org/indi/retro/sald.html | This data repository was supported by the National Natural Science Foundation of China (31470981; 31571137; 31500885), National Outstanding young people plan, the Program for the Top Young Talents by Chongqing, the Fundamental Research Funds for the Central Universities (SWU1509383,SWU1509451,SWU1609177), Natural Science Foundation of Chongqing (cstc2015jcyjA10106), Fok Ying Tung Education Foundation (151023) , General Financial Grant from the China Postdoctoral Science Foundation (2015M572423, 2015M580767), Special Funds from the Chongqing Postdoctoral Science Foundation (Xm2015037, Xm2016044), Key research for Humanities and social sciences of Ministry of Education (14JJD880009). | [17] |
| SLIM | https://fcon_1000.projects.nitrc.org/indi/retro/southwestuni_qiu_index.html | This data repository was supported by: The National Natural Science Foundation of China (31271087; 31470981; 31571137; 31500885). National Outstanding young people planthe Program for the Top Young Talents by Chongqing, the Fundamental Research Funds for the Central Universities (SWU1509383,SWU1509451). Natural Science Foundation of Chongqing (cstc2015jcyjA10106). Fok Ying Tung Education Foundation (151023). General Financial Grant from the China Postdoctoral Science Foundation (2015M572423, 2015M580767). Special Funds from the Chongqing Postdoctoral Science Foundation (Xm2015037). Key research for Humanities and social sciences of Ministry of Education(14JJD880009). | [18] |
| Enhanced Nathan Kline Institute - Rockland Sample (NKI) | http://fcon_1000.projects.nitrc.org/ | Principal support for the enhanced NKI-RS project is provided by the NIMH BRAINS R01MH094639-01 (PI Milham). Funding for key personnel also provided in part by the New York State Office of Mental Health and Research Foundation for Mental Hygiene. Funding for the decompression and augmentation of administrative and phenotypic protocols provided by a grant from the Child Mind Institute (1FDN2012-1). Additional personnel support provided by the Center for the Developing Brain at the Child Mind Institute, as well as NIMH R01MH081218, R01MH083246, and R21MH084126. Project support also provided by the NKI Center for Advanced Brain Imaging (CABI), the Brain Research Foundation, and the Stavros Niarchos Foundation. | [19] |
| SRPBS-OPEN | https://bicr-resource.atr.jp/srpbsopen/ | Data used in the preparation of this work were obtained from the DecNef Project Brain Data Repository (https://bicr-resource.atr.jp/srpbsopen/) gathered by a consortium as part of the Japanese Strategic Research Program for the Promotion of Brain Science (SRPBS) supported by the Japanese Advanced Research and Development Programs for Medical Innovation (AMED) | [20] |
| SRPBS-1600 | https://bicr-resource.atr.jp/srpbs1600/ | Data used in the preparation of this work were obtained from the DecNef Project Brain Data Repository (https://bicr-resource.atr.jp/srpbsopen/) gathered by a consortium as part of the Japanese Strategic Research Program for the Promotion of Brain Science (SRPBS) supported by the Japanese Advanced Research and Development Programs for Medical Innovation (AMED) | [20] |
| HCP1200 | https://www.humanconnectome.org/study/hcp-young-adult/document/1200-subjects-data-release | Data were provided [in part] by the Human Connectome Project, WU-Minn Consortium (Principal Investigators: David Van Essen and Kamil Ugurbil; 1U54MH091657) funded by the 16 NIH Institutes and Centers that support the NIH Blueprint for Neuroscience Research; and by the McDonnell Center for Systems Neuroscience at Washington University. | [21] |

| | | | |
|---|---|---|---|
| ADNI | https://adni.loni.usc.edu/ | Data collection and sharing for this project was funded by the Alzheimer's Disease Neuroimaging Initiative(ADNI) (National Institutes of Health Grant U01 AG024904) and DOD ADNI (Department of Defense awardnumber W81XWH-12-2-0012). ADNI is funded by the National Institute on Aging, the National Institute ofBiomedical Imaging and Bioengineering, and through generous contributions from the following: AbbVie,Alzheimer's Association; Alzheimer's Drug Discovery Foundation; Araclon Biotech; BioClinica, Inc.; Biogen;Bristol-Myers Squibb Company; CereSpir, Inc.; Cogstate; Eisai Inc.; Elan Pharmaceuticals, Inc.; Eli Lilly andCompany; EuroImmun; F. Hoffmann-La Roche Ltd and its affiliated company Genentech, Inc.; Fujirebio; GEHealthcare; IXICO Ltd.; Janssen Alzheimer Immunotherapy Research & Development, LLC.; Johnson &Johnson Pharmaceutical Research & Development LLC.; Lumosity; Lundbeck; Merck & Co., Inc.; MesoScale Diagnostics, LLC.; NeuroRx Research; Neurotrack Technologies; Novartis PharmaceuticalsCorporation; Pfizer Inc.; Piramal Imaging; Servier; Takeda Pharmaceutical Company; and TransitionTherapeutics. The Canadian Institutes of Health Research is providing funds to support ADNI clinical sitesin Canada. Private sector contributions are facilitated by the Foundation for the National Institutes of Health(www.fnih.org). The grantee organization is the Northern California Institute for Research and Education,and the study is coordinated by the Alzheimer's Therapeutic Research Institute at the University of SouthernCalifornia. ADNI data are disseminated by the Laboratory for Neuro Imaging at the University of SouthernCalifornia. | [22] |
| ds000030 | https://openfmri.org/dataset/ds000030/ | This data was obtained from the OpenNeuro database. This work was supported by the Consortium for Neuropsychiatric Phenomics (NIH Roadmap for Medical Research grants UL1-DE019580, RL1MH083268, RL1MH083269, RL1DA024853, RL1MH083270, RL1LM009833, PL1MH083271, and PL1NS062410). | [23] |
| ds001408 | https://openneuro.org/datasets/ds001408 /versions/1.0.1 | This data was obtained from the OpenNeuro database. This work was supported by the EEGInfo Association. | [24] |
| ds001747 | https://openneuro.org/datasets/ds001747 /versions/1.0.0 | This data was obtained from the OpenNeuro database. | [25] |
| ds001796 | https://openneuro.org/datasets/ds001796 /versions/1.5.0 | This data was obtained from the OpenNeuro database. | [26] |
| ds002330 | https://openneuro.org/datasets/ds002330 /versions/1.1.0 | This data was obtained from the OpenNeuro database. | [27] |
| ds002785 | https://openneuro.org/datasets/ds002785 /versions/2.0.0 | This data was obtained from the OpenNeuro database. | [28] |
| ds002790 | https://openneuro.org/datasets/ds002790 /versions/2.0.0 | This data was obtained from the OpenNeuro database. | [28] |
| ds003037 | https://openneuro.org/datasets/ds003037 /versions/1.0.0 | This data was obtained from the OpenNeuro database. | [29] |
| ds003346 | https://openneuro.org/datasets/ds003346 /versions/1.1.2 | This data was obtained from the OpenNeuro database. | [30] |
| ds003469 | https://openneuro.org/datasets/ds003469 /versions/1.0.0 | This data was obtained from the OpenNeuro database. This dataset was provided by Ekaterina V. Pechenkova, Yana R. Panikratova, Maria A. Fomina, Alena D. Rumshiskaya, Darya A. Bazhenova, Liudmila A. Makovskaya, Irina S. Lebedeva, Valentin E. Sinitsyn. This research funded by RFBR grant 18-00-01598(18-00-01592). | [31] |
| ds003831 | https://openneuro.org/datasets/ds003831 /versions/1.0.0 | This data was obtained from the OpenNeuro database. | [32] |
| ds003974 | https://openneuro.org/datasets/ds003974 /versions/3.0.0 | This data was obtained from the OpenNeuro database. | [33–35] |

| | | | |
|---|---|---|---|
| ds003988 | https://openneuro.org/datasets/ds003988/versions/1.0.0 | This data was obtained from the OpenNeuro database. | 33–35 |
| ds004144 | https://openneuro.org/datasets/ds004144/versions/1.0.2 | This data was obtained from the OpenNeuro database. This study is supported by Xochitl Duque, Psychology Department of the Instituto Nacional de Psiquiatria, and the Mexican Foundation for Fibromyalgia for their help recruiting. Ethics Approval: Research Ethics Committee of the Instituto Nacional de Psiquiatria. | 36 |
| ds004169 | https://openneuro.org/datasets/ds004169/versions/1.0.6 | This data was obtained from the OpenNeuro database. This study is supported by National Institute of Child Health and Human Development (R01 HD050735), National Health and Medical Research Council (486682, 1009064) | 37 |
| ds004469 | https://openneuro.org/datasets/ds004469/versions/1.1.2 | This data was obtained from the OpenNeuro database. | 38 |
| ds004636 | https://openneuro.org/datasets/ds004636/versions/1.0.2 | This data was obtained from the OpenNeuro database. This work was supported by the National Institutes of Health (NIH) Science of Behavior Change Common Fund Program through an award administered by the National Institute for Drug Abuse (NIDA) (UH2DA041713; PIs: Marsch, LA & Poldrack, RA). This study was approved by the Stanford Institutional Review Board (IRB protocol #39322) | 39 |
| ds004648 | https://openneuro.org/datasets/ds004648/versions/1.0.0 | This data was obtained from the OpenNeuro database. | 40 |
| ds004697 | https://openneuro.org/datasets/ds004697/versions/1.0.2 | This data was obtained from the OpenNeuro database. | 41 |
| ds004718 | https://openneuro.org/datasets/ds004718/versions/1.1.2 | This data was obtained from the OpenNeuro database. | 42 |

#### 23. Supplementary Table 2. Overview of the included datasets

| | Train Set | | | | Test Set | | | | Clinical Set | | | |
|---|---|---|---|---|---|---|---|---|---|---|---|---|
| Site | N | F/M | Age (mean) | Age (std) | N | F/M | Age (mean) | Age (std) | N | F/M | Age (mean) | Age (std) |
| ABIDE-NYU-Siemens | 48 | 12/36 | 15.25 | 6.39 | 54 | 14/40 | 15.44 | 6.41 | 77 | 10/67 | 14.16 | 6.98 |
| ABIDE-UM_1 | 25 | 7/18 | 13.48 | 3.34 | 23 | 8/15 | 14.09 | 3.16 | 33 | 7/26 | 13.06 | 2.49 |
| ABIDE-UM_2 | 11 | 1/10 | 16.18 | 4.21 | 8 | 0/8 | 16.5 | 4.07 | 12 | 1/11 | 14.67 | 1.37 |
| ABIDE-LEUVEN_2 | 10 | 3/7 | 13.8 | 1.48 | 10 | 2/8 | 13.9 | 1.6 | 12 | 2/10 | 13.58 | 0.9 |
| ABIDE-CALTECH | 11 | 2/9 | 30.55 | 11.72 | 8 | 2/6 | 25.62 | 10.8 | 19 | 4/15 | 27 | 10.34 |
| ABIDE-PITT | 11 | 2/9 | 19 | 6.39 | 10 | 0/10 | 19.6 | 6.67 | 22 | 3/19 | 18.95 | 7.38 |
| ABIDE-SDSU | 12 | 4/8 | 13.75 | 2.09 | 10 | 2/8 | 13.5 | 1.84 | 12 | 0/12 | 14.58 | 1.73 |
| ABIDE-STANFORD | 10 | 2/8 | 9.5 | 1.27 | 6 | 2/4 | 9.33 | 2.16 | 17 | 4/13 | 9.47 | 1.62 |
| ABIDE-TRINITY | 11 | 0/11 | 17.82 | 4.21 | 12 | 0/12 | 16.08 | 3.06 | 22 | 0/22 | 17.27 | 3.4 |
| ABIDE-USM | 20 | 0/20 | 21.05 | 7.4 | 20 | 0/20 | 22.45 | 7.37 | 44 | 0/44 | 22.77 | 7.78 |
| ABIDE-YALE | 14 | 4/10 | 12.21 | 3.02 | 9 | 3/6 | 12.44 | 3.09 | 8 | 1/7 | 14.5 | 2.07 |
| ABIDE2-ABIDEII-GU_1 | 20 | 10/10 | 9.95 | 1.67 | 22 | 13/9 | 9.95 | 1.94 | 25 | 2/23 | 10.76 | 1.48 |
| ABIDE2-ABIDEII-IU_1 | 12 | 3/9 | 24.08 | 5.25 | 8 | 2/6 | 23.25 | 4.62 | 19 | 4/15 | 25.05 | 9.56 |
| ABIDE2-ABIDEII-ONRC_2 | 18 | 8/10 | 24.78 | 3.59 | 17 | 7/10 | 23.24 | 3.6 | 22 | 2/20 | 21.64 | 3.79 |
| ABIDE2-ABIDEII-SDSU_1 | 13 | 1/12 | 13 | 3.16 | 10 | 1/9 | 12.6 | 3.03 | 28 | 6/22 | 12.89 | 3.15 |
| ABIDEII-NYU | 15 | 2/13 | 9.73 | 4.23 | 13 | 0/13 | 8.15 | 2.03 | 68 | 6/62 | 8.34 | 5 |
| CAMCAN-MRC_Cognition_and_Brain_Sciences_Unit | 425 | 205/220 | 49.59 | 18.04 | | | | | | | | |
| CORR-BNU_1 | 57 | 27/30 | 23.05 | 2.29 | | | | | | | | |
| CORR-BNU_2 | 61 | 28/33 | 20.89 | 0.9 | | | | | | | | |
| CORR-BNU_3 | 47 | 23/24 | 22.4 | 1.95 | | | | | | | | |
| CORR-HNU_1 | 30 | 15/15 | 24.37 | 2.41 | | | | | | | | |
| CORR-IACAS | 25 | 13/12 | 26.72 | 5.2 | | | | | | | | |

| | | | | |
|---|---|---|---|---|
| CORR-IBA_TRT | 30 | 15/15 | 25.5 | 6.36 |
| CORR-IPCAS_1 | 30 | 21/9 | 20.9 | 1.77 |
| CORR-IPCAS_3 | 28 | 19/9 | 21.18 | 1.89 |
| CORR-IPCAS_4 | 20 | 10/10 | 23.15 | 1.6 |
| CORR-IPCAS_5 | 22 | 0/22 | 18.32 | 0.48 |
| CORR-IPCAS_8 | 11 | 5/6 | 57.64 | 4.13 |
| CORR-JHNU | 30 | 9/21 | 23.27 | 3.69 |
| CORR-MRN | 33 | 17/16 | 23.24 | 10.34 |
| CORR-NYU_1 | 25 | 15/10 | 29.44 | 8.64 |
| CORR-SWU_1 | 20 | 14/6 | 21.55 | 1.76 |
| CORR-SWU_2 | 27 | 18/9 | 20.96 | 1.65 |
| CORR-SWU_3 | 23 | 15/8 | 20.43 | 1.65 |
| CORR-SWU_4 | 230 | 230/0 | 20.04 | 1.29 |
| CORR-UM | 61 | 43/18 | 65.33 | 6.35 |
| CORR-UPSM_1 | 77 | 37/40 | 15.08 | 2.71 |
| FCON1000_AnnArbor_a | 23 | 0/23 | 20.22 | 7.65 |
| FCON1000_AnnArbor_b | 35 | 0/35 | 47.09 | 26.14 |
| FCON1000_Atlanta | 25 | 0/25 | 29.2 | 8.61 |
| FCON1000_Bangor | 18 | 0/18 | 23.83 | 5.44 |
| FCON1000_Beijing_Zang | 196 | 0/196 | 21.17 | 1.83 |
| FCON1000_ICBM | 77 | 0/77 | 41.49 | 16.54 |
| FCON1000_Milwaukee_b | 45 | 0/45 | 53.33 | 5.6 |
| FCON1000_NYU_TRT | 25 | 15/10 | 29.44 | 8.64 |
| FCON1000_NewHaven_a | 19 | 9/10 | 31 | 10.33 |
| FCON1000_NewHaven_b | 16 | 8/8 | 26.87 | 6.28 |
| FCON1000_NewYork_a | 34 | 16/18 | 30.76 | 9.09 |
| FCON1000_NewYork_a_ADHD | 23 | 0/23 | 33.22 | 8.77 |
| FCON1000_NewYork_b | 20 | 0/20 | 29.75 | 9.94 |
| FCON1000_Newark | 18 | 0/18 | 23.28 | 1.56 |

| | | | | | | | | | | | | |
|---|---|---|---|---|---|---|---|---|---|---|---|---|
| FCON1000_Orangeburg | 15 | 0/15 | 39 | 10.74 | | | | | | | | |
| FCON1000_Oulu | 103 | 67/33 | 21.52 | 0.57 | | | | | | | | |
| FCON1000_Oxford | 21 | 0/21 | 29 | 3.89 | | | | | | | | |
| FCON1000_PaloAlto | 15 | 7/8 | 12.13 | 3.09 | | | | | | | | |
| FCON1000_Pittsburgh | 14 | 5/9 | 36.29 | 8.71 | | | | | | | | |
| FCON1000_Queensland | 19 | 0/19 | 25.95 | 3.88 | | | | | | | | |
| HBN-Site-CBIC | 32 | 17/15 | 11.28 | 3.8 | 26 | 12/14 | 12.31 | 3.53 | 503 | 202/301 | 11.22 | 3.39 |
| HBN-Site-CUNY | 12 | 2/10 | 11.38 | 4.53 | 9 | 2/7 | 10.8 | 1.3 | 100 | 40/60 | 11.62 | 3.23 |
| HBN-Site-RU | 52 | 20/32 | 11.62 | 4 | 53 | 19/34 | 11.32 | 3.82 | 339 | 124/215 | 11.52 | 3.56 |
| HBN-Site-SI | 27 | 14/13 | 12.26 | 4.13 | 28 | 15/13 | 11.5 | 3.85 | 91 | 37/54 | 12.23 | 3.92 |
| NKIRS_Siemens | 94 | 65/29 | 37.88 | 23.61 | 89 | 60/29 | 38.85 | 23.15 | 86 | 60/26 | 36.21 | 18.53 |
| SALD | 424 | 260/164 | 43.15 | 17.21 | | | | | | | | |
| SLIM | 569 | 323/246 | 20.09 | 1.28 | | | | | | | | |
| SRPBS-OPEN-ATV | 39 | 10/29 | 22.69 | 2.2 | | | | | | | | |
| SRPBS-OPEN-HRC | 22 | 14/8 | 44.82 | 12.05 | 26 | 21/5 | 39.35 | 11.06 | 15 | 10/5 | 39.87 | 11.59 |
| SRPBS-OPEN-HUH | 34 | 20/14 | 32.56 | 10.61 | 32 | 17/15 | 36.75 | 15.01 | 57 | 25/32 | 43.33 | 12.18 |
| SRPBS-OPEN-KUT | 75 | 30/45 | 37.55 | 14.25 | 76 | 32/44 | 35.01 | 12.57 | 57 | 29/28 | 42.18 | 11.04 |
| SRPBS_1600-HUH | 49 | 34/15 | 38.37 | 12.54 | 61 | 38/23 | 37.28 | 13.26 | 70 | 33/37 | 42.8 | 12.12 |
| SRPBS_1600-KYU | 46 | 18/28 | 28.85 | 9.58 | 29 | 9/20 | 28.97 | 8.32 | 45 | 19/26 | 37.58 | 9.83 |
| ds000030 | 55 | 28/27 | 31.09 | 8.73 | 54 | 25/29 | 30.19 | 7.84 | 75 | 26/49 | 33.45 | 9.82 |
| ds001408 | 43 | 43/0 | 26.74 | 6.35 | | | | | | | | |
| ds001747 | 89 | 49/40 | 21.65 | 2.5 | | | | | | | | |
| ds001796 | 61 | 47/14 | 32.1 | 7.69 | | | | | | | | |
| ds002330 | 64 | 36/28 | 26.63 | 4.22 | | | | | | | | |
| ds002785 | 201 | 116/85 | 21.82 | 1.82 | | | | | | | | |
| ds002790 | 221 | 126/95 | 21.58 | 1.81 | | | | | | | | |
| ds003037 | 62 | 7/55 | 33.94 | 8.39 | | | | | | | | |
| ds003346 | 52 | 52/0 | 30.35 | 8.38 | | | | | | | | |
| ds003469 | 69 | 47/23 | 24.25 | 5.54 | | | | | | | | |

| | | | | |
|---|---|---|---|---|
| ds003831 | 62 | 42/20 | 38.21 | 14.35 |
| ds003974 | 39 | 22/17 | 22.54 | 4.7 |
| ds003988 | 46 | 24/22 | 25.8 | 4.88 |
| ds004144 | 29 | 29/0 | 41.83 | 6.2 |
| ds004169 | 901 | 566/335 | 20.87 | 3.92 |
| ds004469 | 35 | 26/9 | 33.2 | 12.35 |
| ds004636 | 103 | 68/35 | 23.65 | 5.42 |
| ds004648 | 87 | 0/87 | 29.64 | 5.41 |
| ds004697 | 53 | 27/26 | 7.3 | 0.46 |
| ds004718 | 40 | 32/8 | 69.38 | 3.73 |

24. Supplementary Table 3. Rest-task discrimination in the held-out HCP1200 cohort

| Task | Method | Accuracy | Sensitivity | Specificity | AUC |
|---|---|---|---|---|---|
| WM | GaussianHMM | 0.6536±0.0180 | 0.6739±0.0307 | 0.6333±0.0345 | 0.7130±0.0176 |
| | LEiDA | 0.7228±0.0144 | 0.7529±0.0279 | 0.6927±0.0232 | 0.7989±0.0085 |
| | EiDA | 0.7277±0.0268 | 0.7332±0.0354 | 0.7223±0.0326 | 0.8030±0.0178 |
| | NeuroLex | 0.7401±0.0120 | 0.7658±0.0158 | 0.7144±0.0380 | 0.8258±0.0193 |
| Gambling | GaussianHMM | 0.7144±0.0196 | 0.7065±0.0148 | 0.7223±0.0470 | 0.7791±0.0201 |
| | LEiDA | 0.6823±0.0228 | 0.6878±0.0223 | 0.6769±0.0466 | 0.7609±0.0166 |
| | EiDA | 0.7110±0.0162 | 0.7144±0.0242 | 0.7075±0.0185 | 0.7825±0.0221 |
| | NeuroLex | 0.7307±0.0143 | 0.7539±0.0298 | 0.7075±0.0365 | 0.8034±0.0241 |
| Language | GaussianHMM | 0.6616±0.0141 | 0.6937±0.0207 | 0.6295±0.0265 | 0.7227±0.0200 |
| | LEiDA | 0.7772±0.0165 | 0.7628±0.0091 | 0.7915±0.0316 | 0.8539±0.0108 |
| | EiDA | 0.7683±0.0280 | 0.7588±0.0395 | 0.7777±0.0194 | 0.8485±0.0243 |
| | NeuroLex | 0.7930±0.0230 | 0.7955±0.0174 | 0.7905±0.0329 | 0.8727±0.0186 |
| Motor | GaussianHMM | 0.6610±0.0084 | 0.6986±0.0315 | 0.6235±0.0326 | 0.7189±0.0117 |
| | LEiDA | 0.6695±0.0341 | 0.6977±0.0372 | 0.6412±0.0364 | 0.7166±0.0304 |
| | EiDA | 0.6596±0.0321 | 0.6858±0.0333 | 0.6333±0.0388 | 0.7139±0.0338 |
| | NeuroLex | 0.6779±0.0306 | 0.6878±0.0300 | 0.6679±0.0525 | 0.7472±0.0334 |
| Relational | GaussianHMM | 0.7391±0.0219 | 0.7016±0.0277 | 0.7767±0.0446 | 0.8155±0.0159 |
| | LEiDA | 0.7460±0.0171 | 0.7510±0.0169 | 0.7411±0.0250 | 0.8222±0.0208 |
| | EiDA | 0.7505±0.0161 | 0.7342±0.0261 | 0.7669±0.0346 | 0.8267±0.0177 |
| | NeuroLex | 0.7579±0.0203 | 0.7658±0.0374 | 0.7499±0.0360 | 0.8444±0.0194 |
| Emotion | GaussianHMM | 0.6927±0.0195 | 0.7095±0.0194 | 0.6759±0.0299 | 0.7539±0.0167 |
| | LEiDA | 0.6156±0.0190 | 0.6354±0.0411 | 0.5959±0.0106 | 0.6744±0.0271 |
| | EiDA | 0.6853±0.0282 | 0.7006±0.0382 | 0.6699±0.0500 | 0.7464±0.0274 |
| | NeuroLex | 0.7194±0.0202 | 0.7331±0.0397 | 0.7055±0.0450 | 0.7943±0.0214 |
| Social | GaussianHMM | 0.6694±0.0224 | 0.6798±0.0316 | 0.6591±0.0219 | 0.7394±0.0160 |
| | LEiDA | 0.6566±0.0264 | 0.6651±0.0317 | 0.6482±0.0627 | 0.7007±0.0229 |
| | EiDA | 0.6917±0.0101 | 0.6917±0.0089 | 0.6916±0.0236 | 0.7490±0.0079 |
| | NeuroLex | 0.6759±0.0267 | 0.6620±0.0150 | 0.6898±0.0441 | 0.7330±0.0224 |

#### 25. Supplementary Table 4. Comparison of model fitting across different function configs

| Metric | Config-1 | Config-2 | Config-3 | Config-4 | Config-5 | Config-6 |
|---|---|---|---|---|---|---|
| Z Scores Mean | **-0.024** | -0.027 | -0.026 | -0.027 | -0.028 | -0.027 |
| Z Scores SD | 1.081 | 1.073 | 1.073 | 1.080 | 1.074 | **1.072** |
| Skewness | **0.078** | 0.091 | 0.080 | 0.084 | 0.093 | 0.086 |
| Excess Kurtosis | **0.157** | 0.176 | 0.176 | 0.167 | 0.171 | 0.181 |
| Normality Test Statistic | **0.987** | 0.986 | **0.987** | 0.986 | 0.986 | **0.987** |
| Logarithmic Score | -1.659 | -1.664 | **-1.658** | -1.663 | -1.665 | -1.662 |
| R Squared | 0.108 | 0.108 | **0.111** | 0.106 | 0.106 | 0.108 |
| MAE | 1.181 | 1.181 | **1.180** | 1.182 | 1.182 | 1.181 |
| MSE | 4.518 | 4.490 | **4.471** | 4.518 | 4.501 | 4.472 |
| RMSE | 1.737 | 1.735 | **1.733** | 1.738 | 1.737 | 1.734 |
| AIC | 22162.640 | 22175.273 | **22150.342** | 22165.221 | 22177.415 | 22152.963 |
| BIC | 22772.940 | 22777.325 | **22764.302** | 22776.915 | 22781.407 | 22768.827 |

#### 26. Supplementary Table 5. Statistical significance of age-related effects modulated by sex of the lifespan

| | Fractional Occupancy | | | Dwell Time | | |
|---|---|---|---|---|---|---|
| | T | P | Corrected P | T | P | Corrected P |
| BST1 | -3.257 | 0.001 | 0.014 | -1.792 | 0.073 | 0.878 |
| BST2 | -1.968 | 0.049 | 0.589 | 1.469 | 0.142 | 1.000 |
| BST3 | -0.757 | 0.449 | 1.000 | 0.537 | 0.591 | 1.000 |
| BST4 | 0.551 | 0.582 | 1.000 | 0.065 | 0.948 | 1.000 |
| BST5 | -3.979 | <0.001 | 0.001 | -3.090 | 0.002 | 0.024 |
| BST6 | -0.493 | 0.622 | 1.000 | 1.292 | 0.196 | 1.000 |
| BST7 | 10.505 | <0.001 | <0.001 | 3.384 | 0.001 | 0.009 |
| BST8 | 2.713 | 0.007 | 0.080 | -1.287 | 0.198 | 1.000 |
| BST9 | 0.551 | 0.582 | 1.000 | 2.482 | 0.013 | 0.157 |
| BST10 | 4.046 | 0.000 | 0.001 | 3.605 | 0.000 | 0.004 |
| BST11 | 2.485 | 0.013 | 0.156 | 1.411 | 0.158 | 1.000 |
| BST12 | 1.530 | 0.126 | 1.000 | 0.735 | 0.463 | 1.000 |

#### 27. Supplementary Table 6. Statistical significance of sex-related effects of the lifespan

| | Fractional Occupancy | | | Dwell Time | | |
|---|---|---|---|---|---|---|
| | T | P | Corrected P | T | P | Corrected P |
| BST1 | -12.094 | <0.001 | <0.001 | -15.537 | <0.001 | <0.001 |
| BST2 | -2.114 | 0.035 | 0.414 | -0.694 | 0.488 | 1.000 |
| BST3 | 2.390 | 0.017 | 0.203 | -0.745 | 0.456 | 1.000 |
| BST4 | 11.960 | <0.001 | <0.001 | 4.940 | <0.001 | <0.001 |
| BST5 | 0.294 | 0.769 | 1.000 | 0.647 | 0.518 | 1.000 |
| BST6 | 10.602 | <0.001 | <0.001 | 5.397 | <0.001 | <0.001 |
| BST7 | -3.428 | 0.001 | 0.007 | -2.832 | 0.005 | 0.056 |
| BST8 | -2.005 | 0.045 | 0.540 | -2.133 | 0.033 | 0.396 |
| BST9 | 4.126 | <0.001 | <0.001 | -0.367 | 0.714 | 1.000 |
| BST10 | 4.279 | <0.001 | <0.001 | 1.740 | 0.082 | 0.983 |
| BST11 | 11.706 | <0.001 | <0.001 | 5.926 | <0.001 | <0.001 |
| BST12 | 7.148 | <0.001 | <0.001 | 3.820 | <0.001 | 0.002 |

28. Supplementary Table 7. Group-wise comparison of normative z-scores between ASD and test HC

| feature type | feature | Effect size | Tval | P | Corrected P |
|---|---|---|---|---|---|
| Fractional Occupancy | BST1 | 0.029 | 0.938 | 0.348 | 0.638 |
| | BST2 | 0.050 | 1.599 | 0.110 | 0.364 |
| | BST3 | -0.037 | -1.165 | 0.244 | 0.537 |
| | BST4 | -0.023 | -0.732 | 0.464 | 0.638 |
| | BST5 | 0.014 | 0.436 | 0.663 | 0.729 |
| | BST6 | -0.047 | -1.506 | 0.132 | 0.364 |
| | BST7 | 0.151 | 4.851 | <0.001 | <0.001 |
| | BST8 | -0.055 | -1.772 | 0.077 | 0.364 |
| | BST9 | 0.002 | 0.050 | 0.960 | 0.880 |
| | BST10 | -0.010 | -0.303 | 0.762 | 0.762 |
| | BST11 | -0.017 | -0.546 | 0.585 | 0.715 |
| | BST12 | -0.024 | -0.769 | 0.442 | 0.638 |
| Dwell Time | BST1 | 0.039 | 1.211 | 0.226 | 0.497 |
| | BST2 | -0.004 | -0.126 | 0.900 | 0.825 |
| | BST3 | -0.064 | -2.038 | 0.042 | 0.230 |
| | BST4 | -0.013 | -0.426 | 0.670 | 0.794 |
| | BST5 | -0.009 | -0.286 | 0.775 | 0.794 |
| | BST6 | -0.043 | -1.364 | 0.173 | 0.476 |
| | BST7 | 0.132 | 4.223 | <0.001 | <0.001 |
| | BST8 | 0.008 | 0.261 | 0.794 | 0.794 |
| | BST9 | 0.023 | 0.738 | 0.461 | 0.794 |
| | BST10 | -0.014 | -0.442 | 0.659 | 0.794 |
| | BST11 | -0.055 | -1.751 | 0.080 | 0.294 |
| | BST12 | -0.014 | -0.456 | 0.649 | 0.794 |

29. Supplementary Table 8. Group-wise comparison of normative z-scores between ADHD and test HC

| feature type | feature | Effect size | Tval | P | Corrected P |
|---|---|---|---|---|---|
| Fractional Occupancy | BST1 | 0.019 | 0.644 | 0.520 | 0.520 |
| | BST2 | 0.140 | 4.676 | <0.001 | <0.001 |
| | BST3 | 0.042 | 1.396 | 0.163 | 0.271 |
| | BST4 | 0.016 | 0.532 | 0.595 | 0.541 |
| | BST5 | 0.062 | 2.058 | 0.040 | 0.099 |
| | BST6 | -0.059 | -1.953 | 0.051 | 0.102 |
| | BST7 | 0.136 | 4.518 | <0.001 | <0.001 |
| | BST8 | -0.020 | -0.654 | 0.513 | 0.520 |
| | BST9 | 0.024 | 0.788 | 0.431 | 0.520 |
| | BST10 | -0.004 | -0.118 | 0.906 | 0.755 |
| | BST11 | -0.020 | -0.654 | 0.513 | 0.520 |
| | BST12 | -0.073 | -2.434 | 0.015 | 0.050 |
| Dwell Time | BST1 | -0.137 | -4.602 | <0.001 | <0.001 |
| | BST2 | 0.018 | 0.582 | 0.560 | 0.204 |
| | BST3 | -0.015 | -0.480 | 0.631 | 0.210 |
| | BST4 | -0.157 | -5.290 | <0.001 | <0.001 |
| | BST5 | -0.030 | -1.003 | 0.316 | 0.141 |
| | BST6 | -0.155 | -5.211 | <0.001 | <0.001 |
| | BST7 | 0.026 | 0.868 | 0.385 | 0.154 |
| | BST8 | -0.126 | -4.201 | 0.000 | <0.001 |
| | BST9 | -0.080 | -2.650 | 0.008 | 0.004 |
| | BST10 | -0.132 | -4.411 | <0.001 | <0.001 |
| | BST11 | -0.087 | -2.877 | 0.004 | 0.002 |
| | BST12 | -0.113 | -3.779 | <0.001 | <0.001 |

#### 30. Supplementary Table 9. Group-wise comparison of normative z-scores between MDD and test HC

| feature type | feature | Effect size | Tval | P | Corrected P |
|---|---|---|---|---|---|
| Fractional Occupancy | BST1 | -0.170 | -4.161 | <0.001 | <0.001 |
| | BST2 | 0.020 | 0.477 | 0.633 | 0.633 |
| | BST3 | 0.123 | 3.071 | 0.002 | 0.011 |
| | BST4 | 0.076 | 1.851 | 0.065 | 0.130 |
| | BST5 | 0.054 | 1.302 | 0.194 | 0.242 |
| | BST6 | -0.003 | -0.080 | 0.937 | 0.785 |
| | BST7 | -0.056 | -1.321 | 0.187 | 0.242 |
| | BST8 | -0.087 | -2.165 | 0.031 | 0.103 |
| | BST9 | -0.063 | -1.524 | 0.128 | 0.214 |
| | BST10 | 0.021 | 0.496 | 0.620 | 0.633 |
| | BST11 | -0.003 | -0.072 | 0.942 | 0.785 |
| | BST12 | 0.077 | 1.878 | 0.061 | 0.130 |
| Dwell Time | BST1 | -0.064 | -1.650 | 0.099 | 0.219 |
| | BST2 | 0.114 | 2.899 | 0.004 | 0.043 |
| | BST3 | 0.090 | 2.283 | 0.023 | 0.097 |
| | BST4 | 0.090 | 2.228 | 0.026 | 0.097 |
| | BST5 | 0.005 | 0.128 | 0.899 | 0.824 |
| | BST6 | -0.031 | -0.790 | 0.430 | 0.591 |
| | BST7 | 0.008 | 0.199 | 0.842 | 0.824 |
| | BST8 | -0.067 | -1.699 | 0.090 | 0.219 |
| | BST9 | -0.024 | -0.600 | 0.549 | 0.671 |
| | BST10 | -0.009 | -0.227 | 0.821 | 0.824 |
| | BST11 | 0.040 | 1.008 | 0.314 | 0.493 |
| | BST12 | 0.059 | 1.508 | 0.132 | 0.242 |

### 31. Supplementary Table 10. Group-wise comparison of normative z-scores between SCZ and test HC

| feature type | feature | Effect size | Tval | P | Corrected P |
|---|---|---|---|---|---|
| Fractional Occupancy | BST1 | -0.136 | -2.782 | 0.006 | 0.071 |
| | BST2 | -0.045 | -0.821 | 0.413 | 0.513 |
| | BST3 | 0.079 | 1.472 | 0.143 | 0.343 |
| | BST4 | 0.044 | 0.830 | 0.408 | 0.513 |
| | BST5 | -0.037 | -0.682 | 0.496 | 0.541 |
| | BST6 | 0.034 | 0.581 | 0.562 | 0.562 |
| | BST7 | 0.097 | 1.647 | 0.102 | 0.305 |
| | BST8 | -0.120 | -2.035 | 0.044 | 0.174 |
| | BST9 | 0.045 | 0.795 | 0.428 | 0.513 |
| | BST10 | 0.060 | 1.024 | 0.308 | 0.513 |
| | BST11 | 0.122 | 2.098 | 0.038 | 0.174 |
| | BST12 | 0.058 | 1.045 | 0.298 | 0.513 |
| Dwell Time | BST1 | -0.057 | -1.084 | 0.280 | 0.612 |
| | BST2 | 0.002 | 0.043 | 0.966 | 0.886 |
| | BST3 | 0.221 | 4.303 | <0.001 | <0.001 |
| | BST4 | 0.016 | 0.289 | 0.773 | 0.886 |
| | BST5 | 0.028 | 0.511 | 0.610 | 0.886 |
| | BST6 | 0.093 | 1.682 | 0.095 | 0.260 |
| | BST7 | 0.113 | 1.910 | 0.058 | 0.260 |
| | BST8 | -0.055 | -0.969 | 0.334 | 0.612 |
| | BST9 | -0.007 | -0.125 | 0.901 | 0.886 |
| | BST10 | 0.005 | 0.099 | 0.921 | 0.886 |
| | BST11 | 0.099 | 1.753 | 0.082 | 0.260 |
| | BST12 | 0.018 | 0.324 | 0.747 | 0.886 |

32. Supplementary Table 11. Group-wise comparison of normative z-scores between ANX and test HC

| feature type | feature | Effect size | T | P | Corrected P |
|---|---|---|---|---|---|
| Fractional Occupancy | BST1 | 0.038 | 0.966 | 0.335 | 0.390 |
| | BST2 | 0.147 | 3.675 | 0.000 | 0.003 |
| | BST3 | 0.035 | 0.888 | 0.375 | 0.390 |
| | BST4 | -0.039 | -0.990 | 0.322 | 0.390 |
| | BST5 | 0.048 | 1.196 | 0.232 | 0.387 |
| | BST6 | -0.011 | -0.268 | 0.789 | 0.717 |
| | BST7 | 0.128 | 3.151 | 0.002 | 0.009 |
| | BST8 | -0.055 | -1.361 | 0.174 | 0.348 |
| | BST9 | 0.074 | 1.809 | 0.071 | 0.178 |
| | BST10 | 0.035 | 0.860 | 0.390 | 0.390 |
| | BST11 | 0.007 | 0.160 | 0.873 | 0.728 |
| | BST12 | -0.085 | -2.100 | 0.036 | 0.121 |
| Dwell Time | BST1 | -0.135 | -3.690 | 0.000 | 0.000 |
| | BST2 | -0.031 | -0.817 | 0.414 | 0.188 |
| | BST3 | -0.047 | -1.289 | 0.198 | 0.099 |
| | BST4 | -0.242 | -6.716 | <0.001 | <0.001 |
| | BST5 | -0.054 | -1.450 | 0.147 | 0.082 |
| | BST6 | -0.147 | -3.886 | <0.001 | <0.001 |
| | BST7 | 0.017 | 0.471 | 0.638 | 0.266 |
| | BST8 | -0.184 | -5.056 | <0.001 | <0.001 |
| | BST9 | -0.063 | -1.736 | 0.083 | 0.052 |
| | BST10 | -0.115 | -3.083 | 0.002 | 0.002 |
| | BST11 | -0.083 | -2.238 | 0.026 | 0.018 |
| | BST12 | -0.130 | -3.585 | <0.001 | <0.001 |
